# Bexobrutideg: A Selective, Catalytic Degrader of Bruton’s Tyrosine Kinase Overcomes Inhibitor Resistance and Suppresses Autoantibody-Mediated Disease

**DOI:** 10.64898/2026.08.01.740538

**Authors:** Mark Noviski, Paul Auger, Danielle Bautista, Nivetha Brathaban, Brandon Bravo, Robert Cass, Ganesh Cherala, Thomas C. Fung, Stefan Gajewski, Dhwani Haria, Zewen Jiang, Dane Karr, Aileen Kelly, Daisuke Kato, Zef A. Konst, Aishwarya Kumar, Mitchell Lavarias, Jun Ma, Filippo Marchioni, Joel McIntosh, Jenny McKinnell, Jeffrey T. Mihalic, Ratul Mukerji, Brent Murphy, Nell Narasappa, Ge Peng, Panga Jaipal Reddy, Daniel W. Robbins, Ryan Rountree, Spriha Singh, Ying Siow Tan, Austin Tenn-McClellan, Dahlia R. Weiss, Jeffrey Wu, Jordan Ye, Stephanie Yung, Davorka Messmer, Cristiana Guiducci, Arthur T. Sands, Gwenn M. Hansen, Frederick Cohen

## Abstract

Bruton’s tyrosine kinase (BTK) transduces B-cell receptor (BCR), Toll-like receptor (TLR), and Fc receptor (FcR) signaling, and overactivation of these pathways drives B-cell malignancies and antibody-mediated autoimmune disease. Small molecule inhibitors block the enzymatic functions of BTK, but this inhibition is undermined by resistance mutations, several of which abolish BTK’s kinase activity yet promote oncogenic signaling through BTK scaffolding functions. We report the discovery and characterization of bexobrutideg (NX-5948), a heterobifunctional degrader that recruits cereblon (CRBN) to selectively degrade BTK while sparing molecular glue neosubstrates. We demonstrate that bexobrutideg acts catalytically, degrading thousands of copies of BTK per molecule per hour, and this event-driven pharmacology renders it resilient to mutations that confer resistance to both covalent- and noncovalent-inhibitors. Bexobrutideg is orally bioavailable, driving deep and durable BTK degradation across species. Bexobrutideg demonstrates strong efficacy in wild-type and ibrutinib-resistant lymphoma models and robustly suppresses pathway activation in models of autoimmune disease.

## Introduction

Bruton’s tyrosine kinase (BTK), a member of the TEC family of non-receptor tyrosine kinases, is an essential effector of BCR signaling. This pathway governs B cell development, activation, and survival, and chronic activation of BCR signaling is a hallmark of many hematological malignancies (Woyach et al., 2012; Pal Singh et al., 2018). BTK additionally mediates TLR/IL-1R and FcR signaling, and inappropriate activation of these BTK-dependent pathways drives diverse autoimmune, inflammatory, and allergic disorders (Brunner et al., 2005; Neys et al., 2021). BTK therefore represents a therapeutic target across both oncology and immunology (Woyach et al., 2012; Pal Singh et al., 2018).

Covalent BTK inhibitors such as ibrutinib, acalabrutinib, and zanubrutinib are approved for the treatment of mantle cell lymphoma (MCL), chronic lymphocytic leukemia (CLL), small lymphocytic lymphoma (SLL), Waldenström’s macroglobulinemia, and marginal zone lymphoma (Wen et al., 2021). These ATP-competitive small molecules form a covalent bond to cysteine 481 in the kinase domain (Neys et al., 2021; Wen et al., 2021). Despite their transformative impact on the treatment of these malignancies, acquired resistance frequently arises, leading to relapse and progression (Salvaris et al., 2025). Mutation of C481, the most common treatment-emergent resistance mutation, prevents covalent bond formation, thereby reducing target occupancy and efficacy of covalent inhibitors (Woyach et al., 2014; Woyach et al., 2017). Noncovalent inhibitors such as pirtobrutinib were developed to retain binding in the presence of C481 mutations, and pirtobrutinib is approved for the treatment of relapsed/refractory MCL and CLL/SLL. However, additional kinase-domain mutations (V416L, T474I, L528W, M437R) arise during pirtobrutinib therapy and confer cross-resistance to covalent inhibitors (Wang et al., 2022). Critically, several of these mutations abolish BTK’s enzymatic activity while sustaining BCR signaling and BTK-dependent growth, reflecting a kinase-independent scaffolding role of BTK in oncogenic signaling (Wang et al., 2022; Dhami et al., 2022; Montoya et al., 2024). Beyond oncology, the covalent inhibitors remibrutinib and rilzabrutinib are approved for the treatment of antihistamine refractory chronic spontaneous urticaria and persistent/chronic immune thrombocytopenia, respectively (Jan et al., 2025; Labanca et al., 2025). However, there remains an unmet need for agents that can produce deeper and more sustained clinical responses in broader patient populations.

Targeted protein degradation is a modality distinct from small molecule inhibition that can eliminate all functions of a given protein, encompassing both kinase and scaffolding functions in the case of BTK (Pogash & Fletcher, 2025; Békés et al., 2022; Li et al., 2024). BTK degraders are heterobifunctional small molecules that recruit BTK into the proximity of an E3 ligase (CRBN in the case of bexobrutideg, the substrate receptor of the CRL4 E3 ubiquitin ligase), driving polyubiquitylation and proteasomal degradation of BTK (Cromm & Crews, 2017; Buhimschi et al., 2018; Zhang et al., 2022). Because this event-driven pharmacology requires only transient target engagement, a well-designed degrader can tolerate substantial losses in binding affinity that would severely reduce the efficacy of inhibitors (Békés et al., 2022).

Small molecules that bind CRBN can act as molecular glues and degrade neosubstrates such as the transcription factors IKZF1 (Ikaros) and IKZF3 (Aiolos), enhancing T-cell activation and IL-2 release (Fuchs, 2023). CRBN neosubstrate degradation may contribute to degrader efficacy in some malignancies, but it is undesirable in CLL and settings defined by excessive immune activation, such as autoimmune disease. We therefore sought to design a potent and selective BTK degrader by joining an optimized BTK ligand with a CRBN binder using a linker engineered to eliminate immunomodulatory activity.

Here we report the discovery and pre-clinical characterization of bexobrutideg (NX-5948). We quantify its catalytic efficiency using a framework that corrects for artifacts that typically confound in vitro degrader studies, namely, non-specific binding and glutarimide ring opening. We define the structural basis for its activity against resistance mutations, and we show how selective degradation of BTK has the potential to translate to tumor growth inhibition and suppression of inflammation across multiple therapeutic indications. The medicinal chemistry optimization that produced the bexobrutideg BTK ligand was reported previously for the related degrader NX-2127 (zelebrudomide) (Robbins et al., 2024). Here we focus on the redesign of the linker and CRBN moiety, changes that confer selective BTK degradation and differentiated pharmacology.

## Results

### Design of a selective, neosubstrate-sparing BTK degrader

Bexobrutideg emerged from optimization of the preclinical degrader, NRX-0492, which pairs a BTK ligand with a 5-substituted pomalidomide-analog that retains the molecular-glue activity of pomalidomide (Zhang et al., 2023). Further optimization of NRX-0492 yielded zelebrudomide (NX-2127), a clinical-stage dual degrader of BTK and IKZF1/3 (Robbins et al., 2024). Dual degradation is advantageous in certain lymphomas but undesirable in CLL and autoimmune indications, so our optimization focused on modifications that eliminated neosubstrate activity while maintaining potent BTK degradation (Table 1).

**Table 1.**
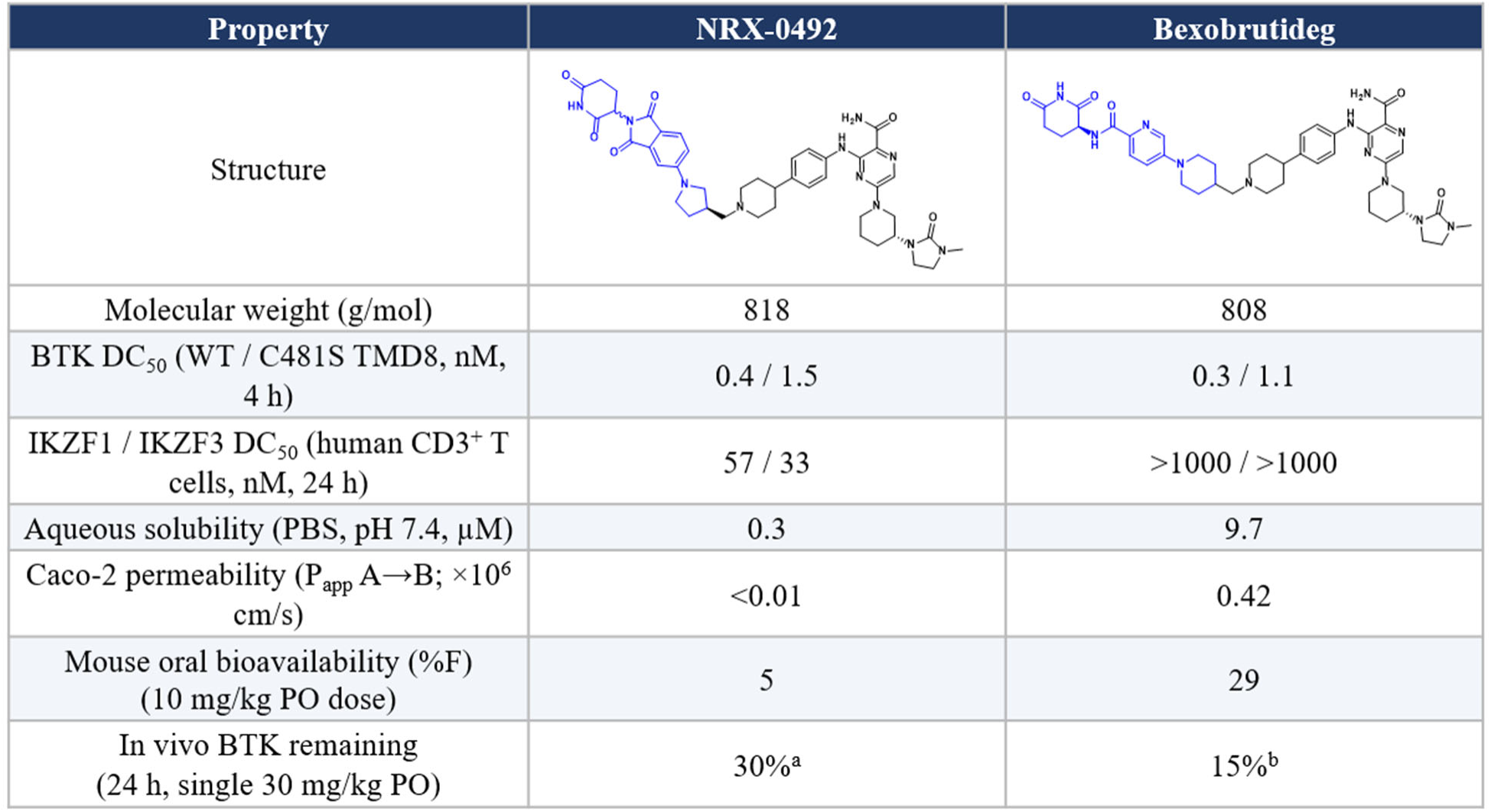
Lead optimization from the preclinical degrader NRX-0492 to bexobrutideg proceeded through redesign of the CRBN binder to preserve potent BTK degradation while eliminating IKZF1/3 neosubstrate activity; the BTK ligand was reported previously (Robbins et al., 2024; Jia et al., 2015). PD assessed by flow cytometry in blood in ^a^BALB/c and ^b^CD-1 mice.

| Property | NRX-0492 | Bexobrutideg |
| --- | --- | --- |
| Structure |  |  |
| Molecular weight (g/mol) | 818 | 808 |
| BTK DC<sub>50</sub> (WT / C481S TMD8, nM, 4 h) | 0.4 / 1.5 | 0.3 / 1.1 |
| IKZF1 / IKZF3 DC<sub>50</sub> (human CD3<sup>+</sup> T cells, nM, 24 h) | 57 / 33 | >1000 / >1000 |
| Aqueous solubility (PBS, pH 7.4, μM) | 0.3 | 9.7 |
| Caco-2 permeability (P<sub>app</sub> A→B; ×10<sup>6</sup> cm/s) | <0.01 | 0.42 |
| Mouse oral bioavailability (%F) (10 mg/kg PO dose) | 5 | 29 |
| In vivo BTK remaining (24 h, single 30 mg/kg PO) | 30%<sup>a</sup> | 15%<sup>b</sup> |

Retaining glutarimide as the CRBN-binding pharmacophore, we replaced the phthalimide with single aromatic rings bearing bridging heteroatoms, appending aryl-amide, lactam, aryl-amine, and aryl-ether groups linked through a 4-alkylpiperidine to the BTK ligand (Supplementary Table 1). These degraders retained CRBN affinity below 1 µM and catalyzed potent BTK degradation in TMD8 cells (DC_50_ 0.4–1.1 nM; D_max_ >94%), while degrading less than 50% of IKZF1/3 in T and B cells at concentrations up to 1 µM, confirming that CRBN affinity and neosubstrate activity are separable (Supplementary Table 1; Supplementary Table 2). Screening in mice at a single oral dose of 30 mg/kg identified a lead compound with high exposure and efficient BTK knockdown; resolution of its diastereomers gave the eutomer bexobrutideg, with defined (S) stereochemistry at the glutarimide chiral center. The distomer degraded BTK markedly less well in vivo (51% versus 15% BTK remaining at 10 mg/kg), and no interconversion of the two isomers was observed in blood or hepatocytes (Supporting Information; Supplementary Table 1; Teo et al., 2003; Mori et al., 2018; Amako et al., 2025). Relative to NRX-0492, the redesigned CRBN binder improved aqueous solubility (9.7 versus 0.3 µM) and permeability, raising oral bioavailability from 5% to 29% (Table 1).

### Bexobrutideg selectively degrades BTK and suppresses B cell activation

In primary human B cells, bexobrutideg induced concentration- and time-dependent BTK degradation (DC_50_ = 0.010 nM and DC_90_ = 0.043 nM at 24 h) and potently suppressed BCR-driven activation, as measured by CD69 induction after anti-IgM stimulation (Fig. 1a-b). Selective BTK degradation was confirmed by global proteomics in TMD8 cells (Fig. 1c; Tohda et al., 2006); the modest decrease in CFLAR (c-FLIP) is a downstream consequence of BTK loss rather than a direct off-target effect, as the inhibitor ibrutinib also decreases CFLAR expression (Nagel et al., 2015). Consistent with a selective, neosubstrate-sparing design, bexobrutideg neither enhanced nor suppressed primary human T cell activation, in contrast to the molecular glues lenalidomide and pomalidomide (Fig. 1d; Robbins et al., 2024). Bexobrutideg thus behaves as a selective BTK degrader without functional immunomodulatory activity or off-target suppression of T cell activation.

**Figure 1.**
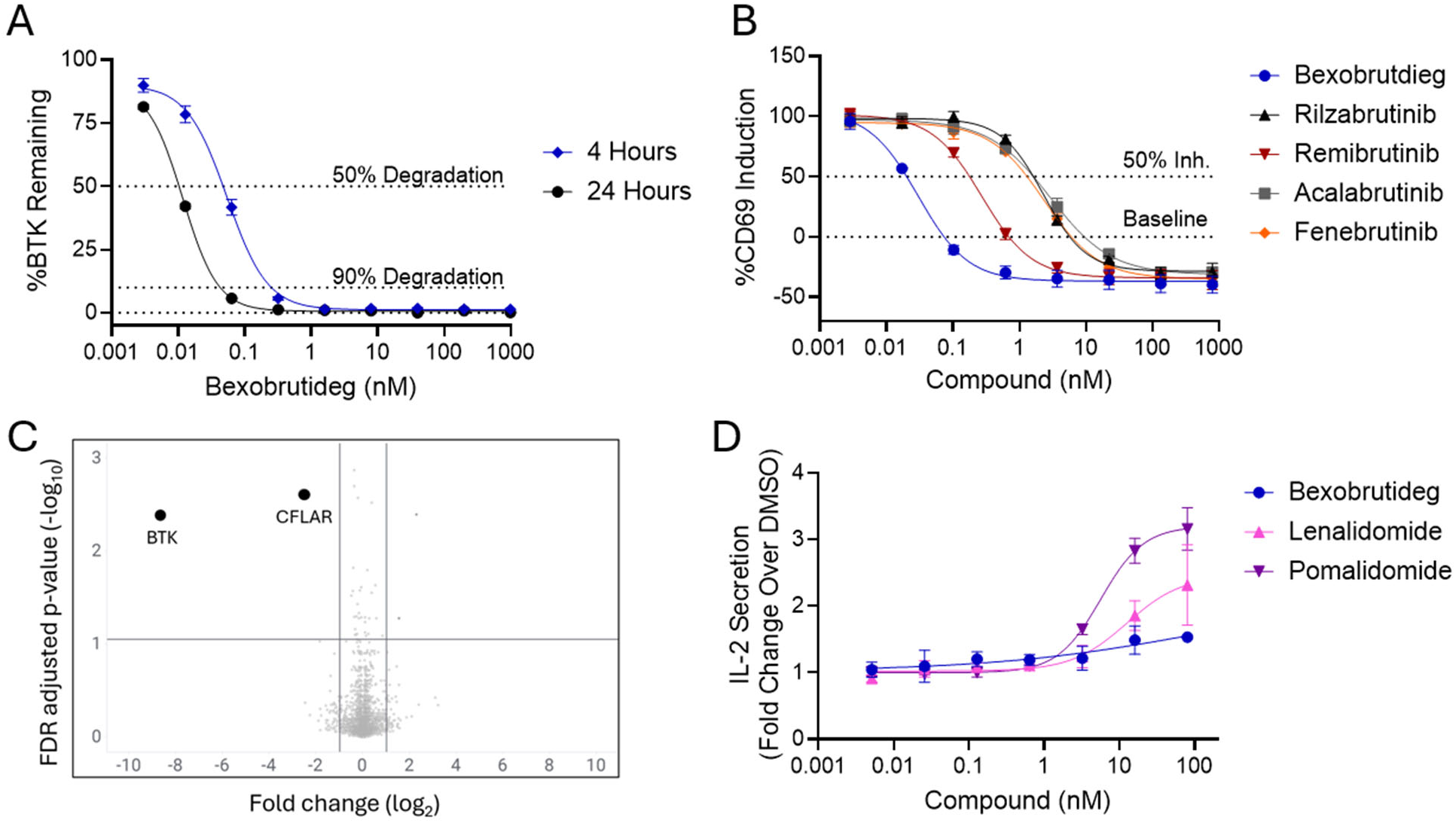
On-target activity and selectivity of bexobrutideg. (a) Human PBMCs from four donors were treated with bexobrutideg for 4 or 24 h, and BTK degradation in B cells was quantified by flow cytometry (b) CD69 induction on B cells after 4 h compound pre-treatment and 19 h stimulation with soluble anti-IgM (30 µg/mL). (c) Global proteomics in TMD8 cells following 6 h treatment with 50 nM bexobrutideg. (d) IL-2 secretion by primary T cells upon 1 hour of compound pre-incubation followed by 24 hours of stimulation with anti-CD3/anti-CD28 in the continued presence of compound; lenalidomide/pomalidomide results were previously reported (Robbins et al., 2024). Mean ± SEM are displayed.

### Structural basis for activity against inhibitor-resistance mutations

Single amino acid changes in BTK’s kinase domain drive clinical resistance by lowering inhibitor-binding affinity and, in certain cases, by conferring kinase-independent scaffold signaling that sustains cell growth (Montoya et al., 2024). Mutations at C481 prevent covalent bond formation, and T474I, V416L, and L528W mutations confer resistance to noncovalent inhibitors and cross-resistance to covalent agents (Wang et al., 2022). X-ray structures of the BTK-binding moiety of bexobrutideg bound to the kinase domain of wild-type or mutant BTK explain its broad mutation tolerance (Fig. 2a-b). The ligand engages the hinge of the kinase domain without extending past the gatekeeper residue T474, avoiding this resistance hotspot (Fig. 2a). The C481 side chain points away from the ligand, so substitutions there minimally perturb binding. Mutations of key hydrophobic residues in the catalytic (V416, L528) and regulatory (M437) spines impair kinase activity and induce scaffold signaling. While the M437R mutation is distal to the ligand binding site and has only negligible effect on ligand affinity (Fig. 2c) the bulky substitutions in the C-spine (V416L and L528W) result in a loss in binding affinity, but BTK’s conformational flexibility allows for displacement of the N-Lobe to still accommodate ligand binding (Fig. 2b; Montoya et al., 2024). We investigated the biochemical and cellular activity of bexobrutideg against these mutants, using pirtobrutinib as a reference inhibitor (Fig. 2c-g). The two agents were equipotent against wild-type BTK in a biochemical probe-displacement assay, and both lost more than tenfold binding potency against several mutants (Fig. 2g). Changes in bexobrutideg binding affinity were confirmed by SPR (Figure S1). However, the downstream consequences of reduced binding potency differed markedly between the molecules. While pirtobrutinib retained the ability to kill TMD8 cells harboring C481S and C481R mutations, cells harboring T474I, V416L, and L528W mutations were resistant to pirtobrutinib (Fig. 2e). By contrast, bexobrutideg retained the ability to degrade mutant BTK with sub-nM activity and high D_max_, potently killing cells harboring each mutation (Fig. 2d-f). For pirtobrutinib, a 17-fold loss in binding to the T474I mutant corresponded with a near-complete loss of cell killing, potentially exacerbated by the hyperactive kinase activity of this mutant (Montoya et al., 2024). Despite a similar drop in binding potency, bexobrutideg retained the ability to kill V416L, C481R, and L528W mutant cells. This highlights an advantage of the degrader modality: a greater resilience to reductions in binary binding affinity compared to small molecule inhibitors.

**Figure 2.**
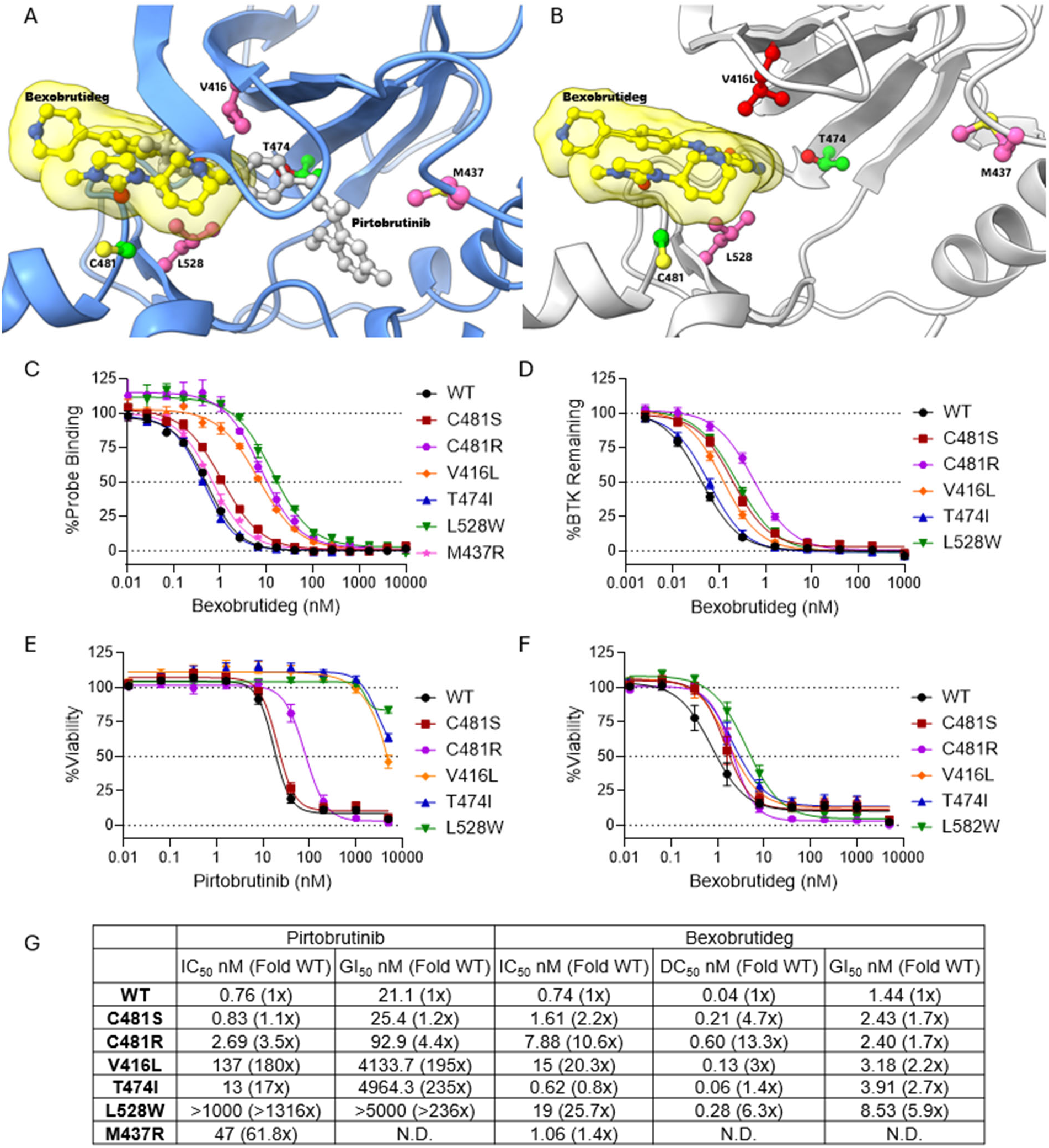
Structural basis for bexobrutideg activity against inhibitor-resistant BTK mutants. (a) BTK-binding moiety of bexobrutideg (yellow) bound to BTK (PDB 8EJB), superimposed on the pirtobrutinib-bound C-lobe (PDB 8FLL); mutation side chains colored by consequence (green, tolerated for kinase activity and ligand binding; purple, ligand binding preserved but kinase activity lost) (Montoya et al., 2024). (b) Bexobrutideg BTK-binding moiety bound to V416L BTK (PDB 36QG). The Cβ atom of residue 416 is displaced by 1.8Å in this mutant compared to 2.2Å in the previously reported L528W mutant BTK. (c) Probe-displacement potency against WT and mutant BTK. (d) BTK degradation in TMD8 cells (24 h), assessed by flow cytometry. (e-f) Viability of TMD8 cells after 72 hours of incubation, as assessed by CellTiter-Glo 2.0 (g) Summary IC_50_/DC_50_/GI_50_ and fold-change vs. WT. Standard deviation and standard error can be found in Supplementary Table 3.

### A quantitative framework for calculating the catalytic efficiency of a degrader

Bexobrutideg’s resilience in the context of mutations that reduce binding affinity reflects its catalytic, event-driven pharmacology. Measuring the catalytic efficiency if a degrader in cellular assays, however, can be confounded by two artifacts impacting the accuracy of dose response assays that are technically challenging to correct: rapid nonspecific binding of degrader to plastic surfaces, and hydrolytic opening of the glutarimide ring (Fig. 3). Adapting the approach of Lynch et al. (2024) to better address these liabilities, we pre-equilibrated compound with media and employed a reverse time-course format to hold exposure stable during the 1-4 h window when most degradation occurs. BTK degradation was strictly concentration- and time-dependent (Fig. 3b-c); when expressed as a function of in vitro compound exposure (concentration × time), degradation curves superimposed independent of incubation time (Fig. 3d), with a hook effect only at 0.5 h.

**Figure 3.**
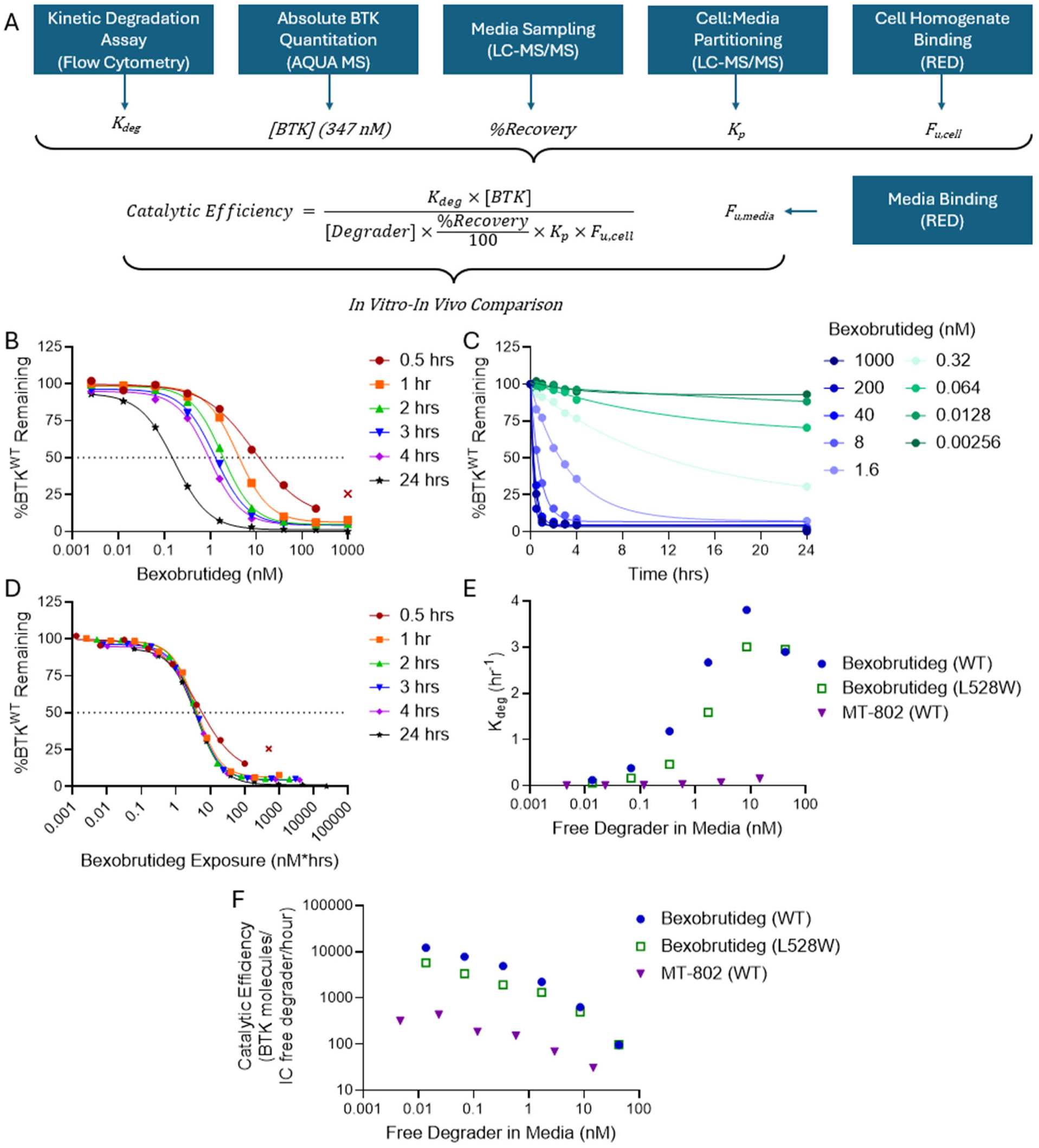
Catalytic activity of bexobrutideg and reference degrader MT-802. (a) Overview of approach taken to quantify catalytic rate. (b) Degradation potency in BTK^WT^ TMD8 cells across incubation times. (c) Degradation kinetics in BTK^WT^. (d) BTK degradation as a function of in vitro bexobrutideg exposure. (e-f) K_deg_ and catalytic efficiency for bexobrutideg and MT-802 against BTK^WT^ and BTK^L528W^.

To convert the degradation rate constant (K_deg_, Fig. 3e) into catalytic efficiency (Fig. 3f), we measured several additional parameters: cell:media partitioning (K_p_), free fractions in media and cell homogenate (F_u,media_, F_u,cell_), compound recovery, and the intracellular concentration of BTK in TMD8 cells. Our calculated BTK concentration of 347 nM closely matched the 360 nM reported previously (Lynch et al., 2024). Bexobrutideg degrades up to 10,000 copies of BTK per free intracellular degrader per hour at free media concentrations of 0.01-0.1 nM (Fig. 3f), a concentration experienced by circulating tumor cells in the first few hours after oral administration of bexobrutideg in the clinic (Palmer et al. 2025; Danilov et al. 2026). Upon administration of the first 600 mg dose (the recommended Phase 2 dose), peak free plasma concentrations reach approximately 0.1 nM, corresponding to ∼5 × 10¹⁰ bexobrutideg molecules/mL. Given a catalytic efficiency of 10,000 BTK molecules degraded per bexobrutideg molecule per hour, the estimated BTK degradation rate at peak concentration is ∼5 × 10¹⁴ BTK molecules/mL/hour (calculated as catalytic efficiency × free plasma concentration). This predicted degradation rate exceeds the estimated BTK burden (∼350 nM, equivalent to ∼6 × 10¹¹ BTK molecules/mL) by several orders of magnitude. Collectively, these estimates predict near-complete BTK degradation within 3 to 4 hours following the first dose. Notably, the observed clinical pharmacodynamic response is consistent with this prediction (Danilov et al., 2026).

To enable direct cross-comparison with Lynch et al., we also quantified the catalytic efficiency of MT-802 (Buhimschi et al., 2018). Our values of 184 and 151 copies/hour at 8 and 40 nM were comparable to the 303 and 56 copies/hour reported by Lynch et al. at 5 and 50 nM. Notably, our K_deg_ values were substantially lower, but this was offset by correcting for loss of MT-802 to nonspecific binding and/or decomposition (Fig. 3f, Supporting Information). The magnitude of this phenomenon, ∼91% loss for bexobrutideg and ∼98% loss for MT-802, argues that nonspecific binding and compound instability must be accounted for in quantitative degrader studies. We further quantified the kinetics and catalytic efficiency of BTK^L528W^ degradation by bexobrutideg, which was modestly slower than in BTK^WT^ but still more rapid and more efficient than MT-802 (Fig. 3d,e). This high catalytic activity explains how bexobrutideg achieves deep degradation and pathway modulation at sub-stoichiometric levels of drug.

### Cross-species pharmacokinetics and pharmacodynamics

Bexobrutideg showed low clearance in mouse and moderate-to-high clearance in rat, dog, and monkey (Supplementary Table 4). Plasma protein binding varied widely across preclinical species (1.2% free in mouse versus 20.3% free in monkey), and oral bioavailability and C_max_ declined in higher species. Despite this, the catalytic, event-driven nature of degradation translated into deep pharmacodynamic effects. Single oral doses of 3, 10, and 30 mg/kg in mice reduced circulating B cell BTK to 30%, 26%, and 15% of baseline at 24 h (Fig. 4a-b), and a 3 mg/kg dose in non-human primates drove robust degradation despite sub-nanomolar free drug exposure (Fig. 4c-d). Degradation was rapid in both species, reaching maximum within two hours, so the low free drug exposure in higher species was sufficient to advance the program.

**Figure 4.**
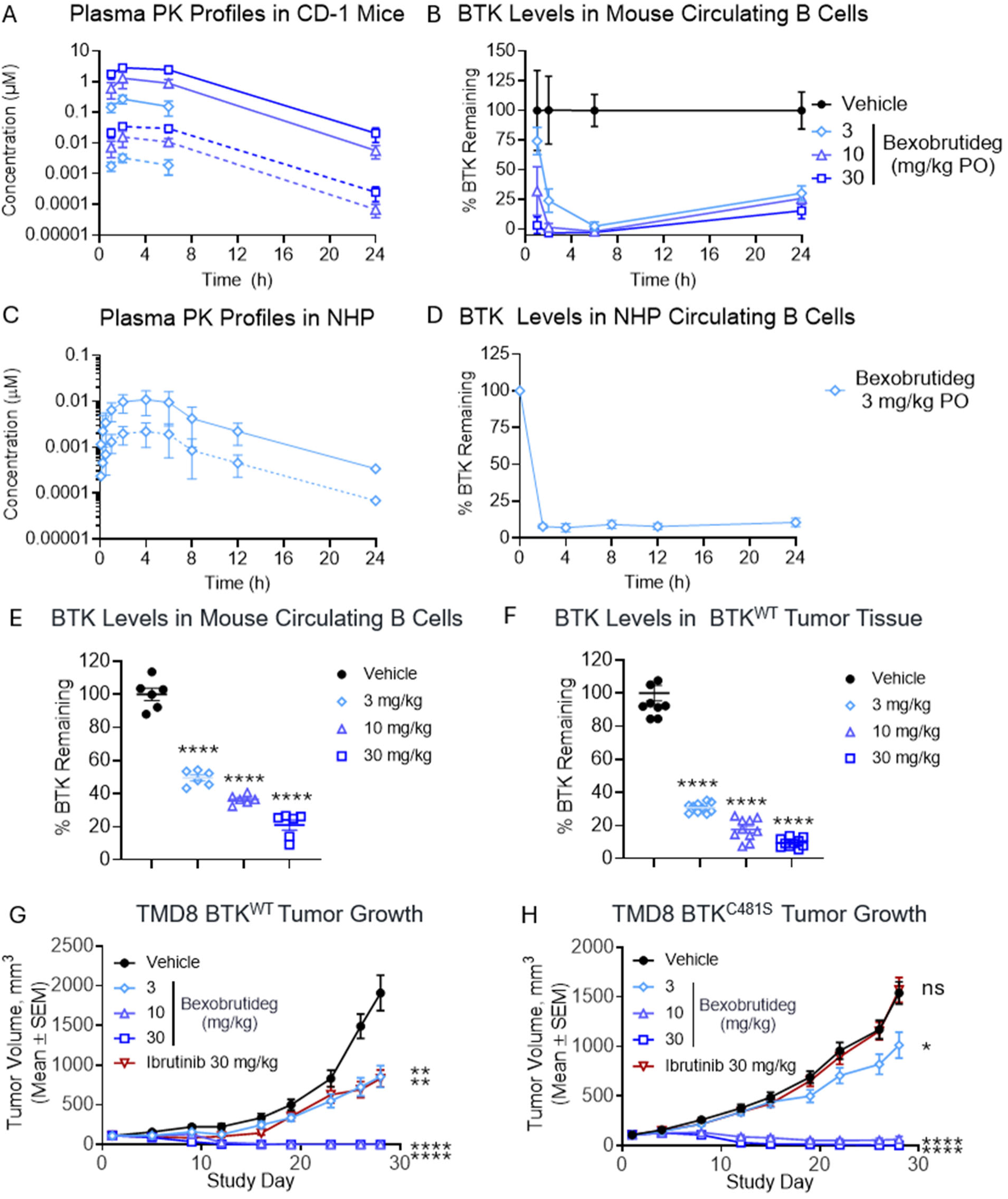
Pharmacokinetics, pharmacodynamics, and antitumor efficacy of bexobrutideg. (a) Plasma exposure and (b) circulating B cell BTK degradation in CD-1 mice after a single oral dose. (c-d) Plasma PK and B cell BTK degradation in NHP following a single oral dose. Solid lines, total plasma concentration; dotted lines, calculated free concentration (mean ± SD). (e-f) BTK degradation in circulating B cells in WT mice and in subcutaneous TMD8 tumor tissue in NSG mice after 5 days of QD oral bexobrutideg dosing (mean ± SEM). (g-h) Impact of bexobrutideg and ibrutinib treatment on tumor growth volume in (g) BTK^WT^ and (h) BTK^C481S^ TMD8 xenografts. One-way ANOVA with Dunnett’s test (ns, not significant; *P<0.05; **P<0.01, ***P<0.001, ****P<0.0001).

### Bexobrutideg is efficacious in wild-type and inhibitor-resistant TMD8 xenograft models

In a TMD8 xenograft model, oral administration of bexobrutideg degraded BTK in tumor tissue to a similar extent as in circulating B cells of non-tumor-bearing mice at matched doses (Fig. 4e-f). Once-daily dosing at 3, 10, and 30 mg/kg for 26 days produced 54%, 100%, and 100% tumor growth inhibition (TGI), versus 57% TGI for ibrutinib at 30 mg/kg (Fig. 4g). In BTK^C481S^ TMD8 tumors, bexobrutideg achieved dose-dependent growth inhibition (99% TGI at 10 mg/kg; 100% at 30 mg/kg), whereas ibrutinib was inactive, as expected (Fig. 4h). Degradation of BTK thus overcomes a resistance mechanism that limits the efficacy of covalent inhibitors, translating the modality’s biochemical resilience into antitumor efficacy.

### BTK degradation by bexobrutideg suppresses BCR, TLR/IL-1R, and FcR signaling in vitro and in vivo

Because BTK mediates BCR, TLR/IL-1R, and FcR signaling, a selective BTK degrader should suppress activation of B cells that mount autoantibody responses as well as the effector functions of those autoantibodies. The high catalytic activity of bexobrutideg led to exceptionally potent suppression of B cell activation upon stimulation with soluble anti-IgM, and the depth of suppression was similar between bexobrutideg and BTK inhibitors (Fig. 1b). Since B cells can respond to antigens presented in diverse physical formats, including surface-bound antigens, we quantified activation across formats and identified a scaffold function of BTK in BCR signaling. Bexobrutideg suppressed B cell activation by particulate anti-IgM more deeply than inhibitors, even at high concentrations of drug (Fig. 5a). Beyond B cells, bexobrutideg promoted potent degradation of BTK in primary human monocytes and differentiated human mast cells. Furthermore, bexobrutideg suppressed IL-1β- and IgG2-mediated activation of monocytes and FcεRI-mediated activation of mast cells (Fig. 5b-d; Fig. S3a-b). This activity translated into strong efficacy in two autoimmune disease models characterized by activation of these pathways: established collagen-induced arthritis (CIA; BCR + TLR/IL-1β + FcγR) and passive cutaneous anaphylaxis (PCA; FcεRI). In the CIA model, oral bexobrutideg resolved paw inflammation and, unlike BTK inhibitors, reduced normalized splenic plasma counts, consistent with degradation providing superior suppression of BCR-mediated plasma cell differentiation (Fig. 5e-f). In the PCA model, oral bexobrutideg administration depleted BTK in ear skin and suppressed both ear swelling and vascular permeability (Fig. 5g-i; Fig. S3c). By contrast, the BTK inhibitor remibrutinib showed trends towards reduction in dye extravasation and ear swelling, but this was not statistically significant. Bexobrutideg can therefore address the full suite of BTK-dependent activities that drive autoimmune and inflammatory disease: autoantibody production, TLR/IL-1R signaling, and Fcγr signaling, and FcεRI-mediated mast cell/ basophil degranulation.

**Figure 5.**
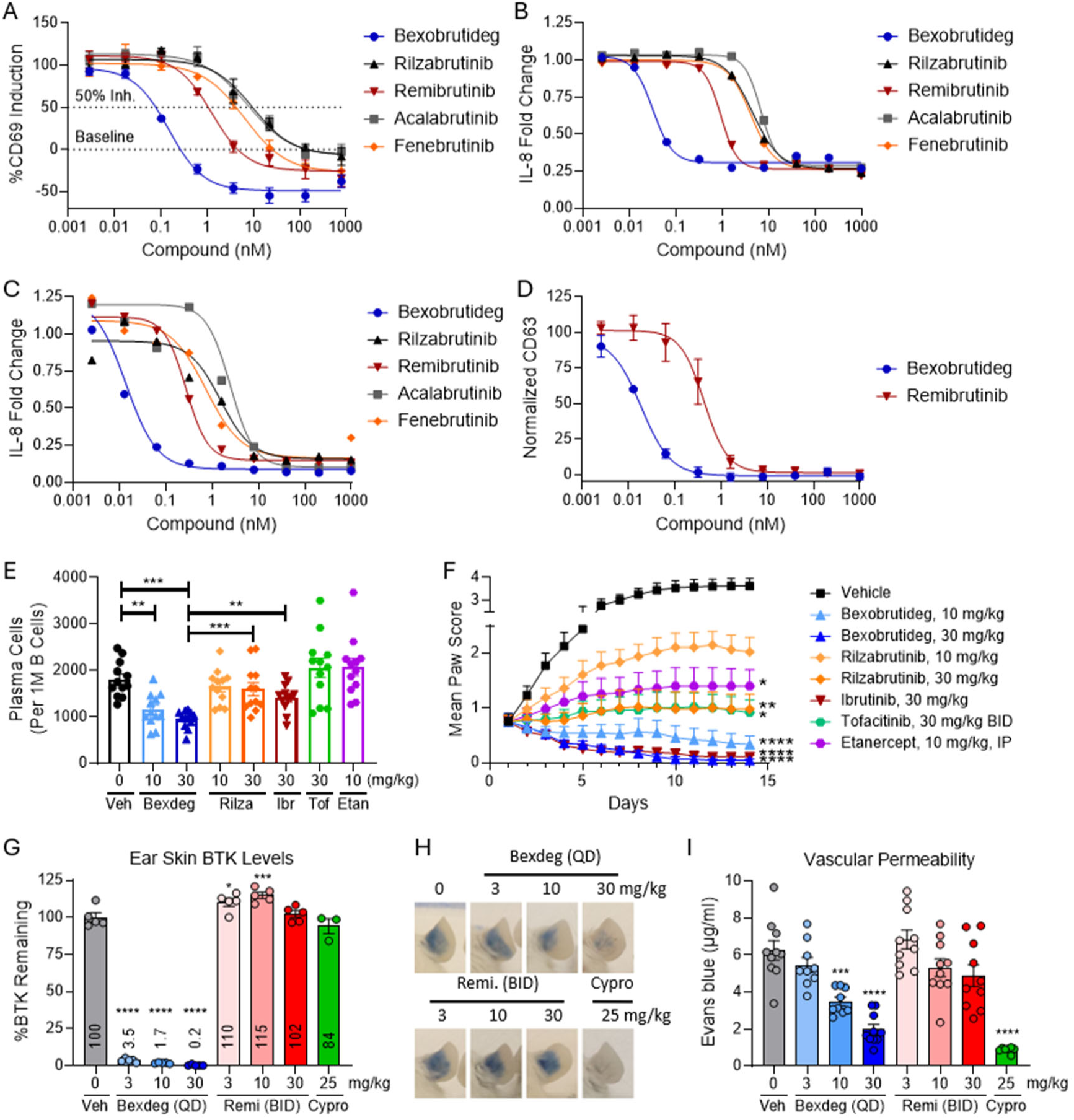
Bexobrutideg suppresses BCR-, TLR/IL-1R-, and FcR-driven effector responses in vitro and in vivo. (a) CD69 induction on human B cells upon bead-bound anti-IgM stimulation. (b-c) IL-8 secretion by human monocytes following stimulation with (b) soluble IL-1β or (c) plate-bound IgG2 (d) Degranulation of differentiated human mast cells pre-treated with compound and IgE for 24 hours and stimulated with anti-IgE; 100% and 0% correspond to CD63 expression DMSO-treated cells with and without anti-IgE stimulation (e-f) Normalized plasma cell counts and clinical score in established CIA model (mean ± SEM). (g-i) BTK degradation in ear skin and dye extravasation in PCA model (mean ± SEM). (*P<0.05; **P<0.01, ***P<0.001, ****P<0.0001). Abbreviations: Bexdeg (bexobrutideg), Rilza (rilzabrutinib), Ibr (ibrutinib), Tof (tofacitinib), Etan (etanercept), Remi (remibrutinib), Cypro (cyproheptadine).

### Preclinical safety

Bexobrutideg was negative in the Ames and in vitro micronucleus genotoxicity assays, did not potently inhibit hERG or five cytochrome P450 isoforms, and showed no time-dependent CYP3A4 inhibition (Supplementary Table 5). This profile, together with IND-enabling studies in mice and NHP, supported advancement of bexobrutideg into clinical development in patients with relapsed or refractory B-cell malignancies (NCT05131022) (Danilov et al., 2026).

## Discussion

Targeted protein degradation is a powerful therapeutic modality that can overcome many limitations of traditional small molecule inhibitors. Catalytic degradation can provide deep and sustained target coverage in disease-relevant tissues at sub-stoichiometric drug levels, eliminating both the enzymatic as well as scaffold activities of a given protein. Additionally, careful design of bifunctional degraders can also confer mutational resilience in the context of malignancy, as well as enhanced selectivity. By employing focused medicinal chemistry strategies to optimize potency, selectivity, and in vivo activity, we developed bexobrutideg, a degrader that exemplifies these differentiated properties with the potential to treat both oncogenic and inflammatory disease.

Bexobrutideg’s mechanism of action provides key advantages across therapeutic contexts. The degrader modality is well-suited to effectively target mutants that elicit kinase-independent scaffold signaling, which are poorly targeted by inhibitors and represent an area of growing unmet medical need. Next, since protein resynthesis, rather than drug clearance drives pathway rebound, the depth and breadth of target coverage achieved by bexobrutideg may enable deeper and more sustained efficacy. This is particularly advantageous in settings where a small amount of pathway activity is sufficient to promote tumor cell survival or inflammatory signaling. This is exemplified by the unique suppression of plasma cell generation by bexobrutideg, as well as the deep efficacy achieved compared to remibrutinib in the PCA model. In both of these cases, a small amount of residual BCR or FcεRI signaling is likely sufficient to drive disease processes, requiring an agent with exceptionally deep target coverage for full efficacy.

Drug-resistance mutations in BTK highlight the susceptibility of occupancy-driven inhibitors to changes in binding affinity. While inhibitors become ineffective with relatively small decreases in affinity, our data demonstrate that bexobrutideg does not. Degradation is event-driven and requires only transient target engagement, allowing bexobrutideg to tolerate >10-fold losses in binary binding affinity. As such, binding affinity inadequately models degrader activity, and the ultimate efficacy of a degrader depends more on its degradation efficiency, a property that can not only be measured, as we have shown in this study, but also a property that can be prospectively optimized.

Translating pharmacokinetic and pharmacodynamic measurements between in vitro and in vivo systems requires precise quantification of active drug in each system. In the case of degraders, in vitro estimates of unbound degrader concentrations can be significantly confounded by nonspecific loss of compound to plastic and protein binding, as well as by compound loss caused by hydrolytic ring opening, in the case of glutarimid-based molecules. When evaluating the catalytic rate of bexobrutideg in this study, these in vitro phenomena amounted to loss of 91-98% of nominal compound. Left uncorrected, these artifacts prevent accurate modeling of in vivo dose and activity relationships as well as in vivo PK/PD modeling. By measuring K_deg_, partitioning, free fractions, recovery, and absolute target abundance, we quantitatively established the catalytic efficiency of bexobrutideg (∼10,000 BTK copies per degrader per hour at 0.01-0.1 nM free media concentration) and offer this adapted framework as a standard for quantitative degrader pharmacology. This high catalytic activity enables deep and sustained pharmacodynamic effects at sub-stoichiometric levels of drug, a distinguishing feature of this category of drug.

Finally, given the right match between compound profile and target biology, an optimized degrader can demonstrate efficacy across therapeutic areas. By designing a highly selective molecule that spares IKZF1/3 degradation and does not display functional immunomodulatory activity, bexobrutideg possesses dual potential in CLL and autoimmunity, where immune activation needs to be suppressed and tolerability is paramount. In this manuscript, we demonstrated the ability of bexobrutideg to suppress BCR, TLR/IL-1R, and FcR signaling, and this translated to exceptional efficacy in autoimmune and inflammatory disease models driven by activation of these pathways in diverse cell types throughout the body. This work with bexobrutideg demonstrates that rigorous characterization of degraders, including selectivity, catalytic efficiency, and activity against resistance mutations, can guide deployment of the modality across both oncology and immunology indications.

## Methods

Full protocols, instrument parameters, crystallography methods, and synthetic procedures are provided in the Supporting Information.

### Chemistry

Synthesis of bexobrutideg and analogs, characterization (^1^H NMR, LC-MS), and stereochemical/interconversion analyses are provided in the Supporting Information.

### Quantification of BTK and IKZF1/3 degradation in vitro

During lead optimization, BTK degradation was screened in TMD8 cells by homogenous time-resolved fluorescence (HTRF) as previously described (Robbins et al., 2024). In human PBMCs, BTK and IKZF1/3 (Ikaros and Aiolos) degradation were quantified by flow cytometry, with 1000 nM compound treatment used for BTK background subtraction and isotype control used for IKZF1/3 background subtraction. %Remaining was calculated by normalizing background-subtracted sample MFI to DMSO-treated control:

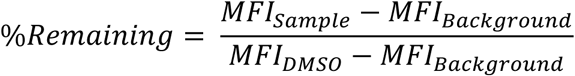

### Quantification of B cell activation

Human PBMCs were isolated from heparanized whole blood, then pre-treated with BTK degrader or inhibitor for 4 hours. Cells were then stimulated with soluble anti-IgM (30 µg/mL) or bead bound anti-IgM (10:1 bead:cell ratio) for 19 hours in the continued presence of compound. Bead-bound anti-IgM was prepared by incubating streptavidin-coated 1.5-1.9 µM polystyrene beads (Spherotech) with biotinylated goat anti-human IgM (4 µg/mL; Jackson ImmunoResearch), after which excess unbound anti-IgM was washed away. CD69 MFIs on B cells (CD3⁻CD20⁺HLA-DR⁺) was quantified by flow cytometry and then normalized using control samples. 100% induction (peak MFI) corresponds to samples stimulated with anti-IgM, and 0% induction (baseline MFI) corresponds to samples treated with media or uncoated bead controls. This is represented by the following formula:

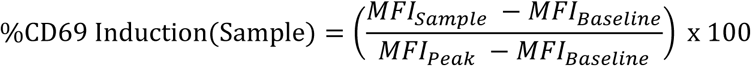

### Quantification of IL-2 secretion by activated T cells

Primary human T cells were isolated from Leukopaks (StemCell, Cat# 70500) by immunomagnetic negative selection (StemCell, Cat#17951), pre-treated with compound for one hour and then stimulated for 24 hours with plate bound anti-CD3 antibody (1 µg OKT3/well) and 5 µg/mL soluble anti-CD28 (clone 28.2) (Life Technologies) in the presence of compound. Total IL-2 in the cell-free supernatant was evaluated by ELISA (R&D, DY202).

### Determination of catalytic efficiency

Degradation kinetics were assessed in TMD8 cells by flow cytometry utilizing a reverse time-course format. Compound was pre-equilibrated as 10x stocks in media for at least 60 minutes prior to addition to cells. %BTK remaining was calculated as described above for PBMCs, and degradation was fit to the GraphPad Prism curve fitting equation “Dissociation-One phase exponential decay,” where K values from Prism are reported as K_deg_ in subsequent calculations:

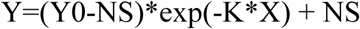

Compound recovery samples were generated in parallel with the kinetic degradation assay, with one set corresponding to the 24 hour timepoint (added first) and another set corresponding to the 0.5 hour timepoint (added last). At assay endpoint, the plate was spun down, and degrader concentration was quantified in the supernatant by LC-MS/MS. To calculate cell:media partitioning, TMD8 cells and media were mixed at a 1:1 v:v ratio and incubated with 1000 nM bexobrutideg or MT-802 for 24 hours. Cells were then spun down, and the cell pellet and supernatant were directly sampled for drug quantification by LC-MS/MS. The partitioning coefficient (K_p_) was calculated by dividing the concentration of drug in the cell pellet by the concentration of drug in the media.

The unbound fraction in TMD8 cell culture media (F_u,media_) was determined using a high-throughput equilibrium dialysis device (HTD 96B, HTDialysis LLC) with 12–14 kDa molecular weight cut-off membranes. The cellular unbound fraction (F_u,cell_) was determined according to the method described by Mateus et al. with minor modifications (Mateus et al., 2017). Cell homogenates were prepared at a density of 50 × 10^6^ cells/mL and incubated with test compound at a final concentration of 1 µM. Equilibrium dialysis was performed against PBS (pH 7.4) at 37 °C for 24 hours at 220 RPM using a high-throughput equilibrium dialysis device (HTD 96B, HTDialysis LLC) with 12–14 kDa molecular weight cut-off membranes.

Absolute intracellular BTK was measured by LC-MS/MS with AQUA stable-isotope-labeled peptides against a standard curve prepared in BTK-depleted matrix.

Catalytic efficiency was calculated using the following formula:

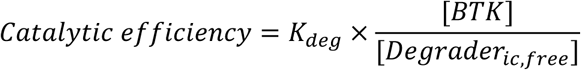

Where K_deg_ and [BTK] were calculated as described above, and [Degrader_ic,free_] is the free intracellular concentration of degrader, calculated as follows:

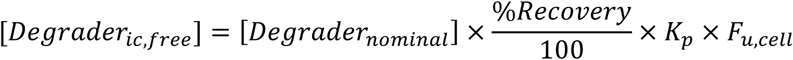

K_deg_ and catalytic efficiency values are plotted as a function of free degrader in media, calculated as follows:

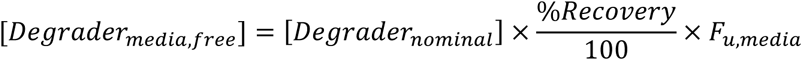

### Monocyte BTK degradation and activation assays

Human primary monocytes were isolated from LeukoPaks (StemCell) using the EasySep immunomagnetic negative selection kit (StemCell) according to manufacturer’s protocol. Monocytes were then pre-treated with BTK degrader or inhibitors for 4 hours followed by overnight stimulation with plate bound anti-IgG2 (10 µg/mL) or soluble IL-1β (10 ng/mL). BTK degradation after 4 hours of bexobrutideg treatment in monocytes (live, CD14^+^) was quantified by flow cytometry, with 1000 nM compound treatment used for BTK background subtraction. %Remaining was calculated by normalizing background-subtracted sample MFI to DMSO-treated control:

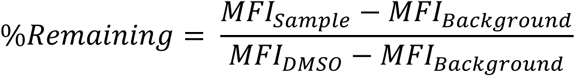

IL-8 secretion was quantified in the supernatants using ELISA (R&D Systems) according to manufacturer’s protocol. Samples were diluted at 1:6 or 1:8 in diluent buffer provided in the ELISA kit. IL-8 concentrations (pg/mL) were interpolated from a 4-parameter logistic ELISA standard curve and multiplied by the appropriate dilution factor. IL-8 fold-change was calculated as the ratio of the mean concentration in the treated samples to that in the matched control as follows:

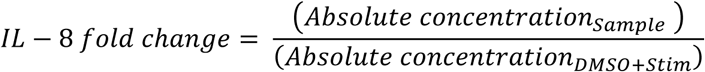

### Mast cell BTK degradation and degranulation assays

Human mast cells differentiated from CD34+ stem cells from two independent donors (Discovery Life Sciences) were treated with bexobrutideg for 24 hours, and BTK degradation was quantified by flow cytometry. Mast cells were also pre-treated for 24 hours with bexobrutideg or remibrutinib and loaded with human IgE (200 ng/mL; Sigma-Aldrich). Media was exchanged to remove unbound IgE, and cells were stimulated with anti-IgE (200 ng/mL; Beckman Coulter) for 90 minutes. Degranulation was quantified by flow cytometry using surface expression of CD63 as a marker of degranulation. DMSO-treated cells, without and with anti-IgE stimulation, were used to quantify baseline (0%) and peak (100%) CD63 induction, respectively, and 2 technical replicates per condition and donor were averaged. CD63 MFIs in compound-treated samples were normalized for each donor using the following equation:

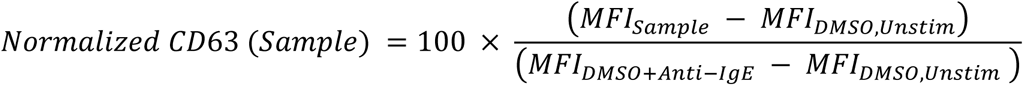

### In vivo pharmacology

Detailed methods for PK/PD, tumor efficacy studies, and the CIA and PCA mouse models are provided in the Supporting Information.

## Data availability

Crystallographic coordinates and structure factors are deposited in the Protein Data Bank under accession code 36QG. Other data supporting the findings are available in the Supporting Information.

## Acknowledgements

Crystallographic data were collected at the NE-CAT beamlines (NIGMS P30 GM124165) using beamtime awards (DOI: 10.46936/APS-182407/60010781) from the Advanced Photon Source under DOE contract DE-AC02-06CH11357. We thank Lori Dasargo for help with manuscript preparation and submission and John Lin for data QC.

## Author contributions

All authors contributed to the design, execution, or interpretation of the studies and to the manuscript, and approved the final version.

## Competing interests

All authors are or were employees of Nurix Therapeutics, Inc. and may hold company stock or stock options.

## LLM usage

LLMs were used during manuscript revision to improve clarity and flow. The authors have reviewed and edited all LLM outputs and attest to their accuracy.

## Notes

### Competing Interest Statement

The authors have declared no competing interest.

