## Supporting Information for "Bexobrutideg: A Selective, Catalytic Degrader of Bruton’s Tyrosine Kinase Overcomes Inhibitor Resistance and Suppresses Autoantibody-Mediated Disease"

### Supplementary Figures

Figure S1. SPR binding kinetics bexobrutideg binding to BTK .......... S3
Figure S2. Kinetics of compound recovery and ring opening .......... S4
Figure S3. BTK degradation in primary human cells and effects in mouse PCA model .......... S4

### Supplementary Tables

Supplementary Table 1. BTK degradation results for survey of CRBN binders .......... S5
Supplementary Table 2. IKZF1/IKZF3 degradation and CRBN binding affinity .......... S7
Supplementary Table 3. Mean ± error for data table in Fig. 2g .......... S7
Supplementary Table 4. Cross-species pharmacokinetics of bexobrutideg .......... S7
Supplementary Table 5. In vitro toxicology profile of bexobrutideg .......... S8
Supplementary Table 6. X-ray data collection and refinement statistics .......... S8

**Supplementary Methods .......... S9**

Crystallography .......... S10
Determination of plasma unbound fraction .......... S10
CRBN FRET probe displacement assay .......... S10
BTK FRET probe displacement assay .......... S11
SPR Methods .......... S11
TMD8 cell culture .......... S12
Generation of BTK mutant knock-in cell lines .......... S12
BTK degradation and viability assays in TMD8 cells .......... S13
Assessment of BTK degradation in human PBMCs by flow cytometry .......... S13

Assessment of IKZF1/3 degradation in human PBMCs by flow cytometry ........................ S15
Determination of catalytic efficiency........................................................................ S15
Quantification of compound recovery and ring opening in culture ..................................... S20
Quantification of B cell activation ............................................................................ S20
Proteomics assay in TMD8 cells.............................................................................. S21
In vivo studies ........................................................................................................ S22
Pharmacodynamic in vivo studies ........................................................................... S22
TMD8 and TMD8 BTK C481S mutant xenograft models .................................................. S23
Established collagen induced arthritis mouse model ........................................................ S24
Passive cutaneous anaphylaxis mouse model ................................................................. S26
Flow cytometry analysis of non-human primate samples.................................................... S26
In vitro interconversion.............................................................................................. S27
Synthetic Procedures................................................................................................. S29
1H NMR and LCMS for Assay Compounds ................................................................... S54
References............................................................................................................... S73

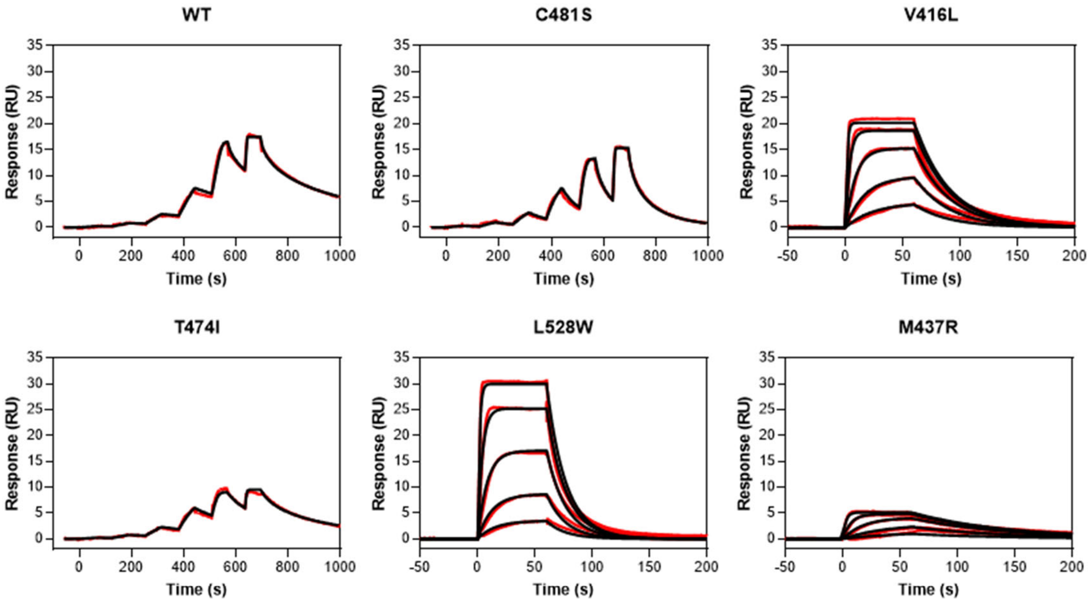


| | $K_D$ (nM) | $k_{on}$ ($10^6$ $M^{-1}s^{-1}$) | $k_{off}$ ($s^{-1}$) |
|---|---|---|---|
| **WT** | 1.8 ± 0.4 | 5.9 ± 3.7 | 0.011 ± 0.009 |
| **C481S** | 5.9 ± 1.9 | 2.4 ± 0.2 | 0.014 ± 0.004 |
| **V416L** | 23 ± 1.7 | 3.5 ± 2.3 | 0.077 ± 0.046 |
| **T474I** | 2.1 ± 0.9 | 1.6 ± 1.1 | 0.004 ± 0.003 |
| **L528W** | 55.3 ± 7.5 | 2.8 ± 1.8 | 0.14 ± 0.09 |
| **M437R** | 3.5 ± 1.5 | 6.1 ± 2.6 | 0.019 ± 0.009 |

**Figure S1**. SPR binding kinetics and representative traces of NX-5948 binding to full-length WT and mutant BTK proteins. Red lines represent raw data and overlaid black lines represent 1:1 binding model. Tabulated data is represented as mean ± SD of 3 or more independent measurements.

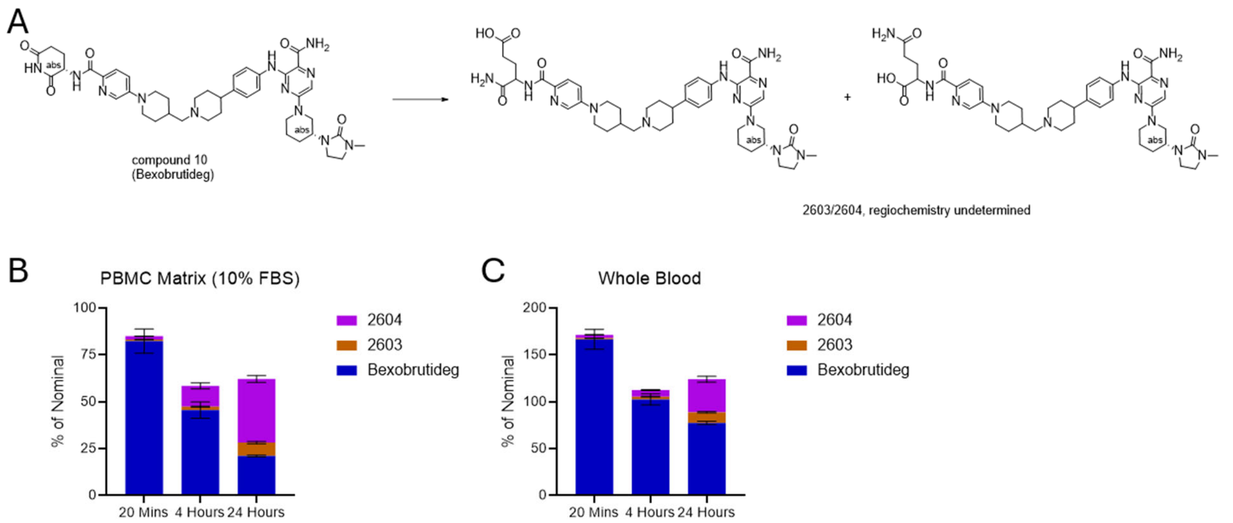


**Figure S2**: Kinetics of compound recovery and ring opening. (A) Bexobrutideg undergoes ring opening to form species 2603 and 2604, which are isomeric ring-opened products from addition of water across the glutarimide. Bexobrutideg was administered to (B) human PBMCs in RPMI-1640 + 10% FBS or (C) human whole blood at the designated timepoints before compound quantitation. At the assay endpoint, bexobrutideg, species 2603, and species 2604 were quantified and normalized to the nominal concentration of bexobrutideg.

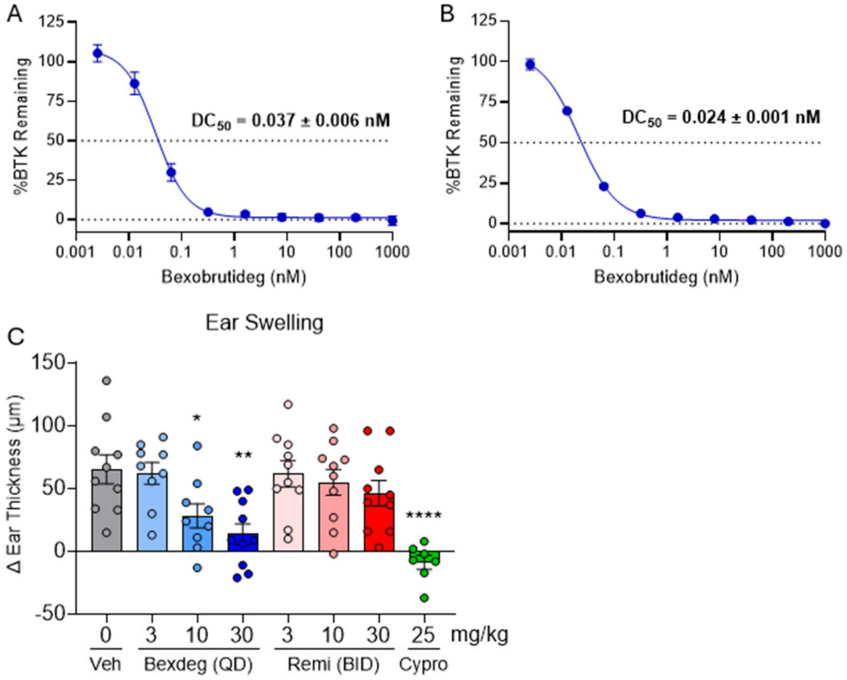


**Figure S3**: (A) BTK degradation in primary human monocytes following 4 h of incubation with bexobrutideg (n = 3 donors). (B) BTK degradation in differentiated human mast cells from 2 independent donors following 24 h of incubation with bexobrutideg. (C) Effect of bexobrutideg (Bexdeg) or remibrutinib (Remi) on ear swelling in the mouse PCA model. On days 1 and 2 of the study, BALB/cJ mice were administered either vehicle or bexobrutideg orally at 3, 10, 30 mg/kg once daily, remibrutinib orally at 3, 10, 30 mg/kg twice daily, or cyproheptadine (Cypro) intraperitoneally at 25 mg/kg once daily. Change in ear thickness was measured prior to and 1 hour after DNP-HSA challenge. Data are represented as mean ± SEM. Statistics were performed by one-way ANOVA with Dunnett's multiple comparisons test. *P < 0.05, **P < 0.01, ***P < 0.001, ****P < 0.0001.

**Supplementary Table 1.** BTK degradation results for a representative survey of CRBN binders. [a]Measured at 4 h by HTRF in TMD8 cells. [b]In vivo BTK degradation was assessed in BALB/c mouse splenocytes 24 h after a single 30 mg/kg PO dose using HTRF, except where noted as determined by flow cytometry. [c]Compound administered at 10 mg/kg PO. [d]Assessed by flow cytometry.

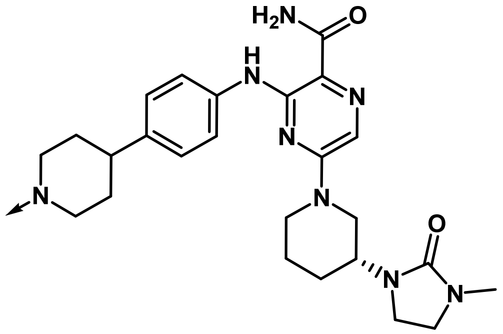


| Example | CRBN binder | BTK[a] $DC_{50}$ (nM) | BTK[a] $D_{max}$ (%) | Plasma conc @ 6 h (μM) 30mg/kg PO | *In vivo* % BTK remaining[b] |
|---|---|---|---|---|---|
| **1 (NRX-0492)** | 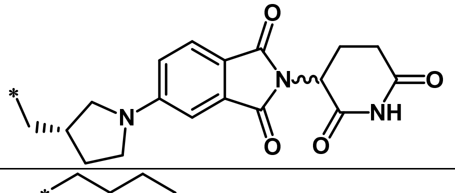 | 0.4 | 97 | 2.21 | 24 |
| **2** | 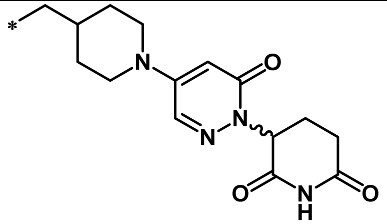 | 0.9 | 97 | 0.038 | 60 |

| | | | | | |
|---|---|---|---|---|---|
| **3** | 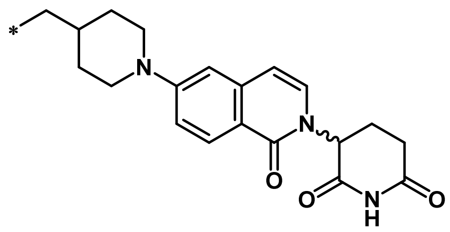 | 0.4 | 97 | 0.55 | 20 |
| **4** | 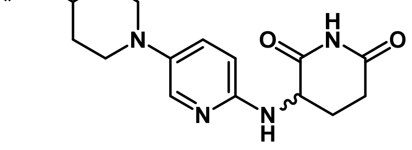 | 1.1 | 94 | 3.43 | 19 |
| **5** | 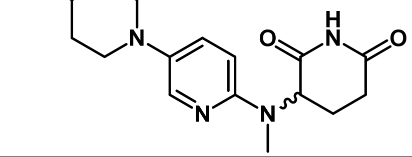 | 3.1 | 94 | - | - |
| **6** | 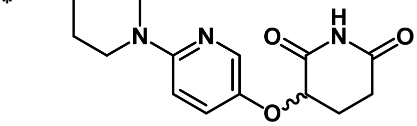 | 0.9 | 96 | 0.91 | 29 |
| **7** | 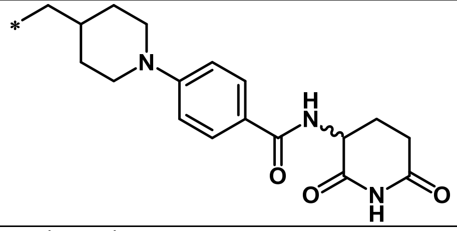 | 1.8 | 96 | - | - |
| **8** | 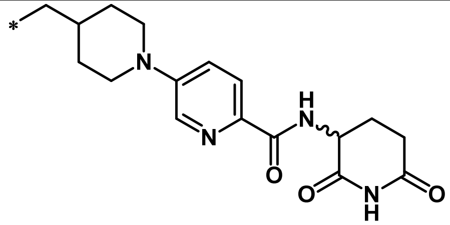 | 0.8 | 97 | 2.97 | 10 |
| **9** | 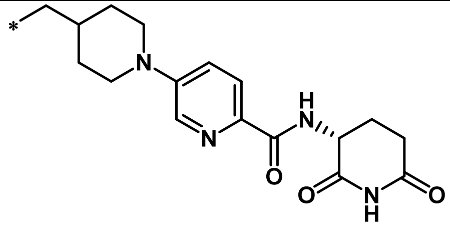 | 5.0 | 96 | 0.616[c] | 51[c, d] |

| | | | | | |
|---|---|---|---|---|---|
| **10** **Bexobrutideg** | 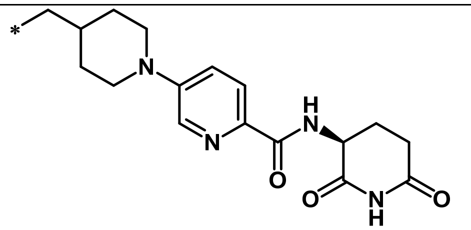 | 0.3 | 98 | 0.695[c] | 15[c, d] |

**Supplementary Table 2.** Ikaros (IKZF1) and Aiolos (IKZF3) degradation and CRBN binding affinity for a representative survey of CRBN binders. IKZF1 and IKZF3 degradation were determined by flow cytometry in human PBMCs, gated on T cells and B cells. CRBN binding affinity was calculated using a probe displacement assay. Chemical structures of each CRBN binder are provided in the synthesis section (Examples 1–10).

| | **T Cells** | | | | **B Cells** | | | | **Biochem.** |
|---|---|---|---|---|---|---|---|---|---|
| | **IKZF3** | | **IKZF1** | | **IKZF3** | | **IKZF1** | | **CRBN** |
| EX. | **$DC_{50}$ (µM)** | **$D_{max}$ (%)** | **$DC_{50}$ (µM)** | **$D_{max}$ (%)** | **$DC_{50}$ (µM)** | **$D_{max}$ (%)** | **$DC_{50}$ (µM)** | **$D_{max}$ (%)** | **$IC_{50}$ (µM)** |
| **1** | 0.004 | 88.25 | 0.017 | 65.13 | 0.008 | 78.69 | >1 | 50.58 | 0.008 |
| **2** | >1 | 18.02 | >1 | 16.84 | >1 | 18.32 | >1 | 11.08 | 0.037 |
| **3** | >1 | 28.47 | >1 | 19.29 | >1 | 25.08 | >1 | 11.89 | 0.007 |
| **4** | >1 | 11.01 | >1 | 12.64 | >1 | 12.89 | >1 | 8.46 | 0.191 |
| **5** | >1 | 14.42 | >1 | 19.59 | >1 | 13.79 | >1 | 12.87 | 0.274 |
| **6** | >1 | 7.88 | >1 | 12.98 | >1 | 8.77 | >1 | 8.02 | 0.901 |
| **7** | >1 | 28.37 | >1 | 22.84 | >1 | 26.38 | >1 | 18.43 | 0.563 |
| **9** | >1 | 19.41 | >1 | 16.50 | >1 | 17.75 | >1 | 9.28 | >1 uM |
| **10** | >1 | 37.90 | >1 | 23.89 | >1 | 30.83 | >1 | 19.42 | 0.0275 |

**Supplementary Table 3.** Mean ± error for data table in Fig. 2g.

| | **Pirtobrutinib** | | **Bexobrutideg** | | |
|---|---|---|---|---|---|
| | **$IC_{50}$ nM (Mean ± SD)** | **$GI_{50}$ nM (Mean ± SEM)** | **$IC_{50}$ nM (Mean ± SD)** | **$DC_{50}$ nM (Mean ± SEM)** | **$GI_{50}$ nM (Mean ± SEM)** |
| **WT** | 0.76 ± 0.26 | 21.1 ± 2.2 | 0.74 ± 0.15 | 0.04 ± 0 | 1.44 ± 0.40 |
| **C481S** | 0.83 ± 0.3 | 25.4 ± 2.4 | 1.61 ± 0.38 | 0.21 ± 0.02 | 2.43 ± 0.58 |
| **C481R** | 2.69 ± 0.67 | 92.9 ± 14.8 | 7.88 ± 0.37 | 0.60 ± 0.08 | 2.40 ± 0.38 |
| **V416L** | 137 ± 71 | 4133.7 ± 350.5 | 15 ± 6 | 0.13 ± 0.02 | 3.18 ± 0.72 |
| **T474I** | 13 ± 3 | 4964.3 ± 35.7 | 0.62 ± 0.14 | 0.06 ± 0.01 | 3.91 ± 0.78 |
| **L528W** | 1000 ± 0 | 5000 ± 0 | 19 ± 5 | 0.28 ± 0.06 | 8.53 ± 3.22 |
| **M437R** | 47 ± 24 | N/A | 1.06 ± 0.36 | N/A | N/A |

**Supplementary Table 4.** Cross-species pharmacokinetics of bexobrutideg. [a]Ultracentrifugation [b]Rapid equilibrium dialysis.

| Parameter | Mouse | Rat | Dog | NHP | Human |
|---|---|---|---|---|---|
| $CL_{obs}$ (mL/min/kg), 1 mg/kg IV | 10.6 | 31 | 68 | 39 | – |
| $V_{ss,obs}$ (L/kg), 1 mg/kg IV | 1.17 | 8.0 | 44.3 | 19.2 | – |
| PO Dose (mg/kg) | 10 | 10 | 5 | 10 | |
| AUC (h·µM), PO | 5.89 | 1.3 | 0.152 | 0.0736 | – |
| $C_{max}$ (µM), PO | 0.891 | 0.098 | 0.0068 | 0.0136 | – |
| Oral bioavailability (%F) | 29 | 16 | 9 | 2 | – |
| Plasma protein binding (% free) | 1.2[a] | 2.0[a] | 5.9[a] | 20.3[a] | 1.6[a] / 0.4[b] |

**Supplementary Table 5.** In vitro toxicology profile of bexobrutideg.

| In vitro preclinical safety assay | Result |
|---|---|
| Ames | Negative |
| In vitro micronucleus | Negative |
| hERG | $IC_{50}$ > 30 µM |
| CYP inhibition (CYP3A4, CYP2D6, CYP2C9, CYP2C19, CYP1A2) | $IC_{50}$ > 30 µM for all isoforms tested |
| CYP3A4 time-dependent inhibition | No time-dependent inhibition signal detected |

**Supplementary Table 6.** Data collection and refinement statistics for the BTK V416L mutant crystal structure bound to the bexobrutideg BTK binding element. The coordinates and data have been deposited in the RCSB as accession number 36QG. Statistics for the highest-resolution shell are shown in parentheses.

| | BTK V416L |
|---|---|
| **Data Collection** | |
| Wavelength [Å] | 0.97915 |
| Resolution range [Å] | 42.71-1.40 (1.44-1.40) |
| Space group | P 2 21 21 |
| Unit cell a,b,c [Å] | 37.99, 73.56, 104.93 |
| Total reflections | 515311 (38811) |
| Unique reflections | 58637 (4245) |

| | |
|---|---|
| Multiplicity | 8.8 (9.1) |
| Completeness (%) | 99.6 (99.6) |
| Mean I/sigma (I) | 17.62 (1.43) |
| Wilson B-factor [Å$^2$] | 21.24 |
| R-merge | 0.049 (0.627) |
| R-meas | 0.052 (0.667) |
| CC (1/2) | 99.9 (71.2) |
| **Model building and refinement** | |
| Reflections used in refinement | 58580 |
| Reflections used for R-free | 1999 |
| R-work | 0.1960 (0.3126) |
| R-free | 0.2123 (0.3108) |
| CC(work) | 0.746 |
| CC(free) | 0.766 |
| non-hydrogen atoms | 2385 |
| protein atoms | 2159 |
| NRX-ligand atoms | 43 |
| other ligands atoms | 5 |
| solvent atoms | 178 |
| Protein residues | 276 |
| **Validation** | |
| RMS(bonds) | 0.007 |
| RMS(angles) | 0.916 |
| Ramachandran favoured [%] | 98.47 |
| Ramachandran allowed [%] | 1.53 |
| Ramachandran outliers [%] | 0.00 |
| Rotamer outliers [%] | 0.44 |
| Clashscore | 5.81 |
| Average B-factor [Å$^2$] | 30.08 |
| protein [Å$^2$] | 29.58 |
| NRX-ligand [Å$^2$] | 38.54 |

| | |
|---|---|
| other ligands [Å$^2$] | 23.72 |
| solvent [Å$^2$] | 34.27 |

### Supplementary Methods

#### *Crystallography*

Crystallographic procedures were performed as published previously (Montoya et al., 2024), but the sample was prepared from V416L mutant BTK kinase domain preincubated at 10-molar excess with 3-{[4-(1-acetylpiperidin-4-yl)phenyl]amino}-5-[(3R)-3-(3-methyl-2-oxoimidazolidin-1-yl)piperidin-1-yl]pyrazine-2-carboxamide. The crystals were grown in 10% (w/v) PEG 8000, 20% (v/v) ethylene glycol, 0.05 M MES, 0.05 M imidazole, 0.03 M diethyleneglycol, 0.03 M triethyleneglycol, 0.03 M tetraethyleneglycol, 0.03 M pentaethyleneglycol, pH 6.5, at 4°C. See Supplementary Table 6 for data collection and refinement statistics.

#### *Determination of plasma unbound fraction*

The unbound fraction in plasma was determined by ultracentrifugation or rapid equilibrium dialysis (RED). For ultracentrifugation, plasma was spiked with test compound at a final concentration of 1 μM and incubated at 37 °C for 10 min to reach binding equilibrium. Spiked matrix was centrifuged at approximately 360,000 × g for 2.5 h at 37 °C (Beckman TLA100.4 rotor), after which supernatant was collected for quantification of unbound concentrations. Each condition was assessed in triplicate.

For RED, plasma was spiked with test compound at a final concentration of 1 μM and placed in a RED device with dialysis membranes of approximately 8 kDa molecular weight cut off (Thermo Scientific, IL, USA). Compounds were dialyzed against blank potassium phosphate buffer (pH 7.4) for 16 h at ambient temperature, after which concentrations were quantified in both chambers to determine the unbound fraction. Each condition was assessed in triplicate.

#### *CRBN FRET probe displacement assay*

Dose titrations of each compound were incubated in a 20 μl reaction volume with 10nM His-tagged CRBN/DDB1 proteins (Nurix), preincubated with 5nM terbium conjugated monoclonal antibody for capturing 6His-tagged proteins (CisBio) and EC40 of a Bodipy-FL labeled small molecule probe in a white 384-well microplate (Perkin Elmer Proxiplate). The final assay buffer was composed of 1x PBS, 0.01% Triton X-100, and 1mM DTT with 2% final DMSO. Following 2-hr incubation, plates were read for TR-FRET signal after excitation at 340nM and read at 520 and 620 nm using an EnVision plate reader (Perkin Elmer). Time-Resolved FRET signal was calculated using a 520/620 nm ratio. Percentage of relative probe binding was calculated compared to DMSO control and fit in ActivityBase (IDBS) to calculate IC50.

*BTK FRET probe displacement assay*

Dose titrations of each compound were incubated in a 20 µL reaction volume with biotinylated BTK (full length) proteins (Nurix), 1 nM WT, C481S, T474I, M437R, L528W, or 2 nM V416L, or 5 nM C481R (SignalChem), preincubated with 1 nM terbium conjugated monoclonal antibody for capturing 6His-tagged proteins (CisBio) and EC50 of a Bodipy-FL labeled small molecule probe in a white 384-well microplate (Perkin Elmer Proxiplate). The final assay buffer was composed of 50 mM HEPES pH7.5, 50mM NaCl, 0.01% Triton X-100, 0.01% BSA, and 1mM DTT with 2% final DMSO. Following 1-hr incubation, plates were read for TR-FRET signal after excitation at 340nM and read at 520 and 620 nm using an EnVision plate reader (Perkin Elmer). Time-Resolved FRET signal was calculated using a 520/620 nm ratio. Percentage of relative probe binding was calculated compared to DMSO control and fit in ActivityBase (IDBS) to calculate IC50.

*SPR Methods*

SPR data was collected using a Biacore S200 instrument (Cytiva Life Sciences). Measurements were performed on a Series-S CM5 sensor chip functionalized with NeutrAvidin (Invitrogen) via amine coupling. First, the sensor surface was activated by a 7-minute injection of 195.5 mM EDC/50 mM NHS. Next, 125 µg/mL NeutrAvidin in 50 mM sodium acetate buffer pH 4.4 was injected over the surface for 150 seconds. The surface was then deactivated with 0.5 M ethanolamine pH 8.0 for 3 minutes. This resulted in a final average NeutrAvidin density of >10,000 resonance units (RU) on each flow cell. All binding measurements were made in the assay buffer containing 50 mM HEPES pH 7.5, 150 mM NaCl, 2 mM MgCl2, 2 mM MnCl2, 5 mM DTT, 0.005% Tween-20, 0.1 mg/ml BSA and 5% DMSO at 25°C. C-terminal biotinylated Avi-tagged BTK (either WT or mutant) diluted to 10-20 µg/mL in assay buffer was captured on flow cells 2-4 of the NeutrAvidin surface to a density of 2000-3000 RU. Flow cell 1 served as the reference flow cell. All flow cells were then subjected to two sequential 30 second injections of 3 mM EZ_Link Amine PEG2 Biotin (Thermo Scientific) at 30 µL/min to block remaining biotin binding sites on the NeutrAvidin surface. Compounds were prepared in assay buffer and serially diluted 3-fold in a 6-point titration scheme. Samples were injected for 60 seconds at 50 µL/min in either multi-cycle or single-cycle mode and allowed varying dissociation times. The resulting data were DMSO-corrected and double referenced prior to analysis. All data was fitted to a kinetic binding model, with fit parameters derived from a 1:1 binding model using Biacore Evaluation software (Cytiva Life Sciences).

*TMD8 cell culture*

TMD8 cells were obtained from Tokyo Medical and Dental University and maintained in MEM alpha media supplemented with 10% heat-inactivated FBS (Tohda et al. 2006. Leukemia Research 30(11): 1385-1390).

*Generation of BTK mutant knock-in cell lines*

TMD8 cells harboring knock-in BTK-C481S, T474I, V416L, and L528W mutations were previously described (Montoya et al., 2024). To generate TMD8 cell lines expressing BTK-C481R mutations, ribonucleoprotein comprised of 20 pmol Cas9 (Thermo Fisher Scientific, A36499) + 120 pmol sgRNA and 100 pmol ssODN repair template were added to Nucleofector Solution SF and incubated for at least 10 minutes at room temperature. TMD8 cells were resuspended in the transfection mixture and nucleofected using the Lonza 4D Nucleofector (Solution SF, program EO-100), then recovered in complete medium + 0.25 µM NU-7441. After overnight incubation, cells were spun down and resuspended in full media. To isolate clones, single cells were plated in a 96-well plate by limiting dilution in 60% complete medium, 30% conditioned medium, and 10% additional FBS. Clones were validated by Sanger sequencing.

*Repair template sequence:*

5'-
AATCTTTCCCATGAGAAGCTGGTGCAGTTGTATGGCGTCTGCACCAAGCAGCGCCCC
ATCTTCATCATCACTGAGTACATGGCCAATGGCAGACTCCTGAACTACCTGAGGGAG
ATGCGCCACCGCTTCCAGACTCAGCAGCTGCTAGAGATGTGCAAGGATGTCTGTGAA
GCCATGGAATACCTGGAGTCAAAGCAGTT-3'

*sgRNA*: 5'- TCTCCCTCAGGTAGTTCAGG-3'

*Primers used to detect BTK C481R mutations:*

Forward: 5’-GATGGGCTCCAAATCCCTGCTT-3’

Reverse: 5’-TCATGGAGCTGAGGCTGGAGAT-3’

Validation of C481R mutation by Sanger sequencing of the endogenous BTK C481 region:

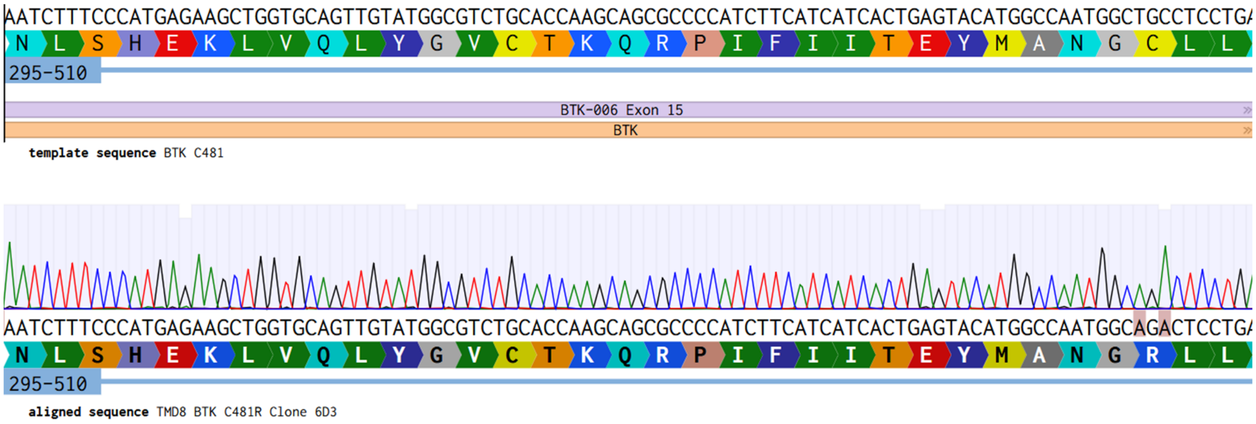


*BTK degradation and viability assays in TMD8 cells*

HTRF degradation assays were performed as previously described. (Robbins et al., 2024). Flow-cytometry based BTK degradation assays in WT and mutant TMD8 cells were performed as previously described (Montoya et al., 2024) except 1000 nM bexobrutideg was used for background subtraction instead of 1000 nM NX-2127. TMD8 viability assays were performed as described in Montoya et al.

*Assessment of BTK degradation in human peripheral blood mononuclear cells (PBMCs) by flow cytometry*

Human PBMCs were isolated from heparinized whole blood obtained from healthy adult donors under an IRB-approved protocol with informed consent (Sanguine BioSciences, Inc.). Blood was diluted 1:1 with PBS, layered onto Ficoll, and centrifuged at 1200×g for 30 min (brake off). The mononuclear cell interface was collected and washed with PBS. PBMCs were resuspended at $2.2\times10^6$ cells/mL in RPMI-1640 + 10% FBS and seeded at 200,000 cells/well in 96-well plates. Compounds were dispensed using a Tecan D300e at a final DMSO concentration of 0.1%, with dosing at 24 h and 4 h prior to endpoint.

At endpoint, cells were fixed in 1× Lyse/Fix Buffer (10 min, 37 °C) and transferred to fresh plates. Following washes with FACS buffer (PBS + 2% FBS + 2 mM EDTA) and 1× Foxp3 permeabilization buffer, cells were stained intracellularly with rabbit anti-BTK (1:100) in the presence of human Fc block (20 µg/mL) for 30 min at room temperature. After washing, a secondary stain was applied using PE-conjugated goat anti-rabbit IgG (1:500) alongside surface markers: BV421 anti-CD20 (1:100), AF488 anti-CD3 (1:100), PerCP-Cy5.5 anti-HLA-DR (1:300), and PE-Dazzle 594 anti-CD14 (1:200). Cells were resuspended in FACS buffer + 2% PFA and analyzed on an Attune NxT flow cytometer.

Flow cytometry data were analyzed using FlowJo, and MFI of BTK in B cells (CD3- CD20+ HLA-DR+) was calculated using the geometric mean of the height parameter. BTK MFI values were normalized using control samples treated with DMSO (peak staining) and 24 h of treatment with 1000 nM bexobrutideg (background staining). Normalized %BTK Remaining values were calculated using the following formula:

$$\%\text{BTK Remaining(Sample)} = \left(\frac{MFI_{Sample} - MFI_{1000\ nM\ bexobrutideg}}{MFI_{Sample} - MFI_{DMSO}}\right) \text{x } 100$$

Using GraphPad Prism, %BTK Remaining values were plotted as a function of compound concentration for each donor and timepoint. Values were fit to the Prism curve-fitting equation "[Inhibitor] vs. response – Variable slope (four parameters)," equation below:

$$\text{Y} = \text{Bottom} + \left(\frac{\text{Top} - \text{Bottom}}{1 + \left(\frac{\text{IC50}^{\text{HillSlope}}}{X^{\text{HillSlope}}}\right)}\right)$$

GraphPad Prism's interpolation function was used to determine the concentrations of bexobrutideg that promoted 50% and 90% degradation of BTK ($DC_{50}$ and $DC_{90}$). The equations for calculating these values are below:

$$DC50 = IC50\ x\ \left(\frac{Top - 50\%}{50\% - Bottom}\right)^{\frac{1}{abs(HillSlope)}}$$

$$DC90 = IC50\ x\ \left(\frac{Top - 10\%}{10\% - Bottom}\right)^{\frac{1}{abs(HillSlope)}}$$

$DC_{50}$ and $DC_{90}$ values were calculated for each individual donor and timepoint, and the mean and standard error of the mean were calculated for each parameter at each timepoint.

*Assessment of IKZF1/3 degradation in human PBMCs by flow cytometry*

Human peripheral blood mononuclear cells (PBMCs) were treated with 1000-0.00512 nM compound for 24 h at 37 °C. Cells were then stained with LIVE/DEAD Fixable Near-IR Dead cell stain kit (ThermoFisher, L10119) for 10 minutes and then fixed and permeabilized using eBioscience Foxp3/Transcription Factor Fixation/Permeabilization Kit (ThermoFisher, 00-5523-00). Cells were then stained with antibodies against IKZF1 (Ikaros) (Biolegend, 368414), IKZF3 (Aiolos) (Biolegend, 371106), CD20 (Biolegend, 302316), and CD3 (BD Pharmingen 552127). Additional samples were stained with Aiolos and Ikaros isotype control antibodies (Biolegend 400136, 400254) as staining controls. Flow cytometry was performed on an Attune NxT Acoustic Focusing Flow Cytometer (Thermo-Fisher A29004) and data were analyzed using FlowJo (v10.5.3) and GraphPad Prism. T cells (CD3+ CD20-) were gated, and the geometric mean fluorescence intensity (MFI) of Aiolos and Ikaros protein was calculated. Data were normalized and degradation parameters were calculated as described for BTK, except isotype control antibody was used for background subtraction.

*Determination of catalytic efficiency*

*Kinetic degradation assay*

TMD8 cells were seeded in 96-well plates at 54,000 cells/well in a volume of 90 µL/well. A 9-point 5-fold serial dilution of degrader (bexobrutideg or MT-802) was prepared in DMSO and then diluted with TMD8 media (alpha-MEM + 10% FBS) to prepare 10x stocks (10,000-0.0256 nM, 1.25% DMSO). 10x stocks were pre-equilibrated at 37°C for at least 60 minutes prior to adding 10 µL of 10x stock to the 24 h samples, diluting compound to 1x (1000-0.00256 nM, 0.125% DMSO). At this time, 6 additional technical replicates were treated with 1000 nM bexobrutideg to serve as background controls. 10x stocks were maintained overnight in a humidified incubator at 37°C and 5% $CO_2$, then 10 µL of 10x stock was added to cells 4, 3, 2, 1, and 0.5 h prior to endpoint. At endpoint, 100 µL of 2x Foxp3 fix/perm reagent (1:1 dilution of 4x concentrate and diluent) was added to cells, which were fixed at room temperature for one hour. Cells were stained with rabbit anti-BTK (1:100), washed twice with permeabilization buffer, then stained with goat anti-rabbit

IgG (1:500); both stains were prepared in permeabilization buffer with human Fc block (1:50). Samples were then analyzed on an Attune NxT Acoustic Focusing Flow Cytometer.

%BTK remaining values were calculated as described above for PBMCs, but without lineage marker gating. For each concentration of drug, %BTK remaining was plotted as a function of time and fit to the GraphPad Prism curve fitting equation "Dissociation- One phase exponential decay," equation below:

$$Y=(Y0-NS)*\exp(-K*X) + NS$$

K values ($K_{deg}$) for bexobrutideg x $BTK^{WT}$, bexobrutideg x $BTK^{L528W}$, and MT-802 x $BTK^{WT}$ were averaged from n = 4, 3, and 1 independent experiments, respectively. Given the weak degradation observed at the bottom three concentrations of drug (0.064, 0.0128, and 0.00256 nM nominal), these conditions are omitted from $K_{deg}$ and catalytic efficiency graphs.

*Quantification of compound recovery, free fraction, and partitioning*

Compound recovery samples were generated in parallel with the kinetic degradation assay, with one set corresponding to the 24 hour timepoint (added first) and another set corresponding to the 0.5 hour timepoint (added last). At assay endpoint, the plate was spun down, and compound concentrations were quantified in the supernatant by LC-MS/MS. Calibration standards and study samples underwent protein precipitation with acetonitrile containing internal standard prior to analysis on a Shimadzu Exion LC system coupled to a Sciex QTRAP 6500+ mass spectrometer. Compound concentrations were calculated from the standard curve generated using internal standard–to-analyte peak area ratios. The limit of quantitation of the assay was 0.62 nM. Drug was quantifiable in the 1000 nM, 200 nM, 40 nM, and 8 nM (nominal) samples for bexobrutideg and the 1000 nM, 200 nM, and 40 nM (nominal) samples for MT-802. The measured (empirical) concentration of drug was then divided by the intended (nominal) concentration of drug and multiplied by 100 to calculate %Recovery. Because %Recovery values did not differ substantially between the 24 and 0.5 hour samples and were consistent across concentrations, all data points were averaged to obtain a uniform %Recovery value for each compound: 8.74% for bexobrutideg and 1.99% for MT-802.

Artifacts from nonspecific plastic binding and dissociation confounded prior efforts to quantify cell:media partitioning at low cell densities. To mitigate this issue, TMD8 cells and media were mixed at a 1:1 v:v ratio and incubated with 1000 nM bexobrutideg or MT-802 for 24 h. Cells were then spun down, and the cell pellet and supernatant were directly sampled for drug quantification by LC-MS/MS. The partitioning coefficient ($K_p$) was calculated by dividing the concentration of drug in the cell pellet by the concentration of drug in the media. $K_p$ values were then averaged using the geometric mean, first by averaging technical replicates within a run, then by averaging between runs, with each run as a biological replicate (n = 5 and 3 runs for bexobrutideg and MT-802, respectively). Average $K_p$ values were similar for both degraders: 13.7 for bexobrutideg and 13.5 for MT-802.

The unbound fraction in TMD8 cell culture media ($F_{u,media}$) was determined using a high-throughput equilibrium dialysis device (HTD 96B, HTDialysis LLC, Gales Ferry, CT, USA) with 12–14 kDa molecular weight cut-off membranes. Media was spiked with test compound at a final concentration of 1 µM and dialyzed against phosphate-buffered saline (PBS, pH 7.4) at 37 °C for 24 h at 220 RPM. At the end of the incubation, samples were collected from both the media and buffer chambers for analysis. Each condition was assessed in triplicate in each experimental run.

The cellular unbound fraction ($F_{u,cell}$) was determined according to the method described by Mateus et al. with minor modifications (Mateus et al., 2017). Cell homogenates were prepared at a density of 50 × $10^6$ cells/mL and incubated with test compound at a final concentration of 1 µM. Equilibrium dialysis was performed against PBS (pH 7.4) at 37 °C for 24 h at 220 RPM using a high-throughput equilibrium dialysis device (HTD 96B, HTDialysis LLC, Gales Ferry, CT, USA) with 12–14 kDa molecular weight cut-off membranes. At the end of the incubation, samples were collected from both the homogenate and buffer chambers for analysis. Each condition was assessed in quadruplicate in each experimental run.

Bexobrutideg cell and media binding was quantified in two independent experimental runs, and MT-802 was assessed in a single run. The average fraction unbound values were as follows:

| | $F_{u,\ media}$ | $F_{u,\ cell}$ |
|---|---|---|
| Bexobrutideg | 0.4875 | 0.00875 |
| MT-802 | 0.734 | 0.0063 |

*Quantification of intracellular BTK protein*

Intracellular BTK concentrations in TMD8 cells were determined by LC-MS/MS using absolute quantification (AQUA) stable isotope labeled (SIL) peptides. TMD8 cells, treated with 1 µM bexobrutideg to degrade endogenous BTK, were used as blank matrix for standard curve preparation. Standard curve generation was accomplished by spiking unlabeled synthetic BTK peptide (Peptide sequence - FTNSETAEHIAQGLR) at 9 different concentrations to yield a calibration curve. The calculation for the curve was based on the peak area ratio relative to AQUA SIL internal standards. AQUA SIL peptides were synthesized, and purity was determined by Cell Signaling Technology (CST). 200K TMD8 cell pellets were lysed in 100 µL of 1% sodium deoxycholate (SDC) 50 mM triethylammonium bicarbonate (TEABC), pH 8.0 for 15 minutes at 90°C. Protein quantification was performed using a Pierce$^{TM}$ BCA Protein Assay Kit. Samples were processed on an Agilent AssayMAP Bravo liquid handling platform using the In-Solution Digestion Single Plate v2.0 protocol. 95 µL of cell lysate was reduced with 25 µL of dithiothreitol (DTT) for 30 minutes at 60°C, followed by alkylation with 25 µL of iodoacetamide for 30 minutes at 25°C. Protein digestion was performed by addition of 25 µL of Trypsin/LysC Mix, Mass Spec Grade (2 µg) and incubation for 720 minutes (12 h) at 37°C. Digestion was quenched by addition of 25 µL of 10% formic acid at 8°C. Resulting peptides were purified using AssayMAP RPS 25 µL cartridges per the Peptide Cleanup v4.0 protocol with a 75 µL elution volume.

Samples were injected on an Agilent 1290 Infinity Liquid Chromatography system connected to an Agilent 6495C QQQ mass spectrometer. 25 µL of sample was injected onto a Zorbax 300SB-C18 Narrow-Bore 2.1x150mm 5-Micron column at 45°C utilizing a flow rate of 0.2 mL/min. Mobile phase A was 0.1% formic acid in LC-MS grade water and mobile phase B was 0.1% formic acid in LC-MS grade acetonitrile. The gradient was a segmented linear gradient starting at 98% A, ramped to 90% A at 6 minutes, and 10% A at 7 minutes before equilibration with a total run time of 10 minutes. The mass spectrometer was operated in positive ion electrospray mode with dynamic multiple reaction monitoring (dMRM) for maximum sensitivity. MRM parameters:

| Analyte | Q1(m/z) | Q3 (m/z) | Average Dwell (msec) | CE (v) | Retention Time (min) |
|---|---|---|---|---|---|

| | | | | | |
|---|---|---|---|---|---|
| FTNSETAEHIAQGLR | 558.6129 +++ | [y8] 923.5057 | 16.34 | 24 | 6.3 |
| | | [y7] 794.4631 | | | |
| | | [y6] 657.4042 | | | |
| **F***TNSETAEHIAQGLR | 561.9553 +++ | [y8] 923.5057 | 16.34 | 24 | 6.3 |
| | | [y7] 794.4631 | | | |
| | | [y6] 657.4042 | | | |

Agilent Mass Hunter software (version 10.1.67) was used for LC-MS/MS instrument control and data acquisition. Skyline was used for data analysis. Peptide concentration was determined against a standard curve of internal standard versus compound peak area ratio (light to heavy) with 9 replicates.

*Calculation of catalytic efficiency*

Catalytic efficiency was calculated using the following formula:

$$Catalytic\ efficiency = K_{deg} \times \frac{[BTK]}{[Degrader_{ic,free}]}$$

Where $K_{deg}$ and [BTK] were calculated as described above, and $[Degrader_{ic,free}]$ is the free intracellular concentration of degrader, calculated as follows:

$$[Degrader_{ic,free}] = [Degrader_{nominal}] \times \frac{\%Recovery}{100} \times K_p \times F_{u,cell}$$

$K_{deg}$ and catalytic efficiency values are plotted as a function of free degrader in media, calculated as follows:

$$[Degrader_{media,free}] = [Degrader_{nominal}] \times \frac{\%Recovery}{100} \times F_{u,media}$$

*Quantification of compound recovery and ring opening in culture*

Given the degree of compound loss observed in the catalytic efficiency assessment, we took an orthogonal approach to quantify the kinetics and relative contribution of nonspecific binding and glutarimide ring opening (Figure S2). We quantified these processes in two different matrices with different protein content: human PBMCs in RPMI-1640 with 10% FBS (same protein content as TMD8 assay) and human whole blood. PBMCs were isolated as described above, and whole blood was collected in K2 EDTA tubes. Bexobrutideg was administered to PBMCs and whole blood in a reverse time course format using an automated Tecan D300e liquid at 1000, 200, and 40 nM, and samples were mixed thoroughly. Timepoints correspond to the amount of time that elapsed between compound administration and sampling of supernatant for LC-MS/MS quantitation of bexobrutideg and ring-opened species. Compound recovery did not differ across concentrations, which were averaged to produce a single recovery value for each analyte at each timepoint. Notably, bexobrutideg demonstrated uniformly lower recovery values in the PBMC matrix (Figure S2B) than in whole blood (Figure S2C), with recovery values of approximately 80% and 160%, respectively, at 20 minutes. Recovery further declined from 20 minutes to 4 h, but the sum of all three species (bexobrutideg + 2603 + 2604) remained stable from 4 to 24 h. While species 2603 and 2604 were detectable at low levels in the PBMC matrix at 4 h, these species outnumbered bexobrutideg at 24 h. By contrast, ring opening proceeded more slowly in the whole blood matrix, where bexobrutideg still outnumbered ring opened species at 24 h. Collectively, these results demonstrate that in tissue culture media (10% FBS), there is a rapid loss of bexobrutideg to nonspecific binding, followed by a more gradual loss of active drug due to ring opening.

*Quantification of B cell activation*

Human PBMCs were isolated from heparinized whole blood obtained from healthy adult donors under an IRB-approved protocol with informed consent (Sanguine BioSciences, Inc.). Blood was diluted 1:1 with PBS, layered onto Ficoll, and centrifuged at 1200×g for 30 min (brake off). The mononuclear cell interface was collected and washed three times with PBS, then resuspended at 2

$\times$ $10^6$ cells/mL in RPMI-1640 supplemented with 10% FBS. Cells were seeded at 200,000 cells/well in 96-well plates. Compounds were dispensed using a Tecan D300e liquid handler in an 8-point, 6-fold dose response (top concentration 800 nM; 0.1% final DMSO). After 4 h of compound pre-incubation, PBMCs were stimulated with anti-IgM (soluble or bead-bound) for 19 h in the continued presence of compound (compound concentrations reflect pre-stimulation values). Positive controls received DMSO + anti-IgM; negative controls received DMSO + media or blank beads.

Following stimulation, cells were stained with live/dead dye, then stained in FACS buffer (PBS + 2% FBS + 2 mM EDTA) with the following antibodies: BV421 anti-CD20 (1:100), AF488 anti-CD3 (1:100), PerCP-Cy5.5 anti-HLA-DR (1:300), PE-Cy7 anti-CD69 (1:100), AF647 anti-CD86 (1:100), and human Fc block (20 µg/mL). Fixed samples were acquired on an Attune NxT flow cytometer and analyzed in FlowJo v10.10.0. CD69 MFIs on B cells ($CD3^{-}CD20^{+}HLA\text{-}DR^{+}$) were normalized using control samples. 100% induction (peak MFI) corresponds to samples stimulated with anti-IgM, and 0% induction (baseline MFI) corresponds to samples treated with media or blank beads. This is represented by the following formula:

$$\%\text{CD69 Induction(Sample)} = \left(\frac{MFI_{Sample} - MFI_{Baseline}}{MFI_{Peak} - MFI_{Baseline}}\right) \text{x } 100$$

Using GraphPad Prism, %CD69 Induction values were plotted as a function of compound concentration for each donor. Values were fit to the Prism curve-fitting equation "[Inhibitor] vs. response – Variable slope (four parameters)," equation below:

$$\text{Y} = \text{Bottom} + \left(\frac{\text{Top} - \text{Bottom}}{1 + \left(\frac{\text{IC50}^{\text{HillSlope}}}{X^{\text{HillSlope}}}\right)}\right)$$

*Proteomics assay in TMD8 cells*

TMD8 cells were treated with DMSO or bexobrutideg at a concentration of 50 nM for 6h. Cells were tested for viability and washed with PBS prior to storage at -80° C. Proteomic sample preparation was performed with PreOmics iST 96x kit (PO.00027, PreOmics, USA), with cell lysis and protein digestion carried out using trypsin following the manufacturer's protocol. After sample processing, liquid chromatography tandem mass spectrometry (LC-MS/MS) data acquisition was conducted via the BoxCar method (Nature Methods volume 15, pages 440–448, 2018).

The acquired BoxCar data was processed in MaxQuant using the BoxCar mode with default settings against human protein database containing random sequences. High pH fractionated data-dependent acquisition (DDA) runs served as library files for BoxCar data analysis. MaxLFQ quantified proteins with a false discovery rate (FDR) cutoff of 1% were exported to the Perseus plugin. Subsequent to data filtration and imputation, the quantified proteins were uploaded into the Nurix-developed application for statistical analysis and visualization.

*In vivo studies*

All animal studies were conducted under approved Institutional Animal Care and Use Committee (IACUC) protocols.

*Pharmacodynamic in vivo studies*

For pharmacodynamic assessment, 8–9-week-old female BALB/cJ mice or CD-1 mice were orally dosed with compound or vehicle. Mice were bled via tail vein or cardiac puncture at the indicated time points. Red blood cells were lysed and remaining cells were stained with live/dead dye, then fixed, permeabilized using FoxP3/Transcription Factor Staining Buffer Set (Invitrogen) and stained with cell surface markers CD45 (Biolegend), TCR β chain (Biolegend), CD3 (Biolegend) and B220 (BD Biosciences). For BTK intracellular staining detection, cells were stained with anti-BTK (clone D3H5, Cell Signaling Technology Cat. 8547BC) primary and AF647-anti-Rabbit (Biolegend) secondary antibodies with mouse FC block (BD553142). Flow cytometry analysis was performed on an Attune NxT Acoustic Focusing Flow Cytometer (Thermo-Fisher A29004) and data were analyzed using FlowJo and GraphPad Prism. B cells (CD45+/B220+/TCR β-) and T cells (CD45+/B220-/TCR β+) BTK MFIs were measured and degradation was calculated using the following equation, with the BTK MFI in T cells used as an internal control for background subtraction:

$$\%BTK\ Remaining = 100 \times \frac{BTK\ MFI_{Sample,\ B\ Cell} - BTK\ MFI_{Sample,\ T\ Cell}}{BTK\ MFI_{Vehicle,\ B\ Cell} - BTK\ MFI_{Vehicle,\ T\ Cell}}$$

*TMD8 and TMD8 BTK$^{C481S}$ mutant xenograft models*

*In vivo Tumor Implantation*

7–8-week-old female CB.17 SCID mice (Fox Chase SCID®, CB17/Icr-Prkdc$^{scid}$/IcrIcoCrl, Charles River) were subcutaneously implanted into the mid-flank with 1 x $10^7$ cells of TMD8 cells resuspended in Hank's Balanced Salt Solution (HBSS) and Matrigel™ (Corning 354234) at a ratio of 1:1.

When tumors reached 60-150 $mm^3$, daily oral treatment of either bexobrutideg, ibrutinib, or vehicle was initiated. Body weight and tumor volumes were measured twice weekly. Tumors were measured using the formula of length x width$^2$ x 0.5. Tumor growth inhibition (% TGI) was calculated using the equation $[1 - (T - T_0/C - T_0)] \times 100$, where T and C represent the mean size of tumors in the treated (T) and control (C) groups, and $T_0$ refers to the tumor size at randomization. Statistical significance of tumor volumes was evaluated on the final measurement day using one-way ANOVA with Dunnett's multiple comparisons test (not significant (ns) $P > 0.05$, * $P \leq 0.05$, ** $P \leq 0.01$, *** $P \leq 0.001$, and **** $P \leq 0.0001$).

*BTK pharmacodynamics in TMD8 tumor tissues*

Tumor cell pellets were lysed in Pierce RIPA Lysis buffer (Fisher, PI89901) with 1x Halt Protease and Phosphatase inhibitor cocktail (ThermoFisher Scientific, 1861281). Lysates were incubated on ice for 30 minutes followed by centrifugation at 14,000 rpm for 10 minutes. The supernatants were collected for BCA (Bioinchoninic Acid) assay for determining total protein concentration measured at a maximum absorbance wavelength of 562 nm using SpectraMax iD3. Jess Simple Western was performed on the TMD8 tumor lysates following the manufacturer's instruction with a protein loading concentration of 2 µg (diluted in 0.1X sample buffer) and capillary cartridges used were derived from the 12–230 kDa separation module (ProteinSimple®, Bio-Techne). DTT (Dithiothreitol), Fluorescent 5× Master Mix, Biotinylated Ladder, and Luminol-Peroxide solution according to manufacturer instructions. 1 part 5× Fluorescent Master Mix was combined with 4 parts diluted lysate in a microcentrifuge tube. Samples were denatured with sample buffer + DTT according to Jess Simple Western guidelines (95 °C, 5 minutes). The rabbit anti-BTK monoclonal antibody (Cell Signaling Technology, 8547) was used at a 1:100 dilution in antibody diluent (Bio-Techne). The rabbit anti-β-Actin antibody (Cell Signaling Technology, 8457) was used at a 1:25 dilution in antibody diluent (Bio-Techne) as the loading control. Chemiluminescence was visualized using an HRP-conjugated anti-rabbit secondary antibody (ProteinSimple®, Bio-Techne). All samples were run on Jess Automated Western Blot System (Bio-Techne).

Data acquisition and analysis was performed with Compass for SW (v6.1.0) and GraphPad Prism (v10.4.1) software. Percent BTK remaining was calculated for compound-treated samples using the following equation, where $AUC_{Norm}$ is the AUC of BTK normalized to AUC of β-Actin in Compass software:

$$\%\text{BTK Remaining (Sample)} = \frac{(\text{AUC}_{Norm,Sample})}{(\text{AUC}_{Norm,Vehicle})} \times 100$$

Statistical significance of the percent BTK remaining between groups was evaluated by one-way ANOVA with Dunnett's multiple comparisons test (ns, not significant; *P>0.05; **P≤0.01, ***P≤0.001, ****P≤0.0001).

*Established collagen induced arthritis mouse model*

DBA/1 mice were injected intradermally with bovine type II collagen on study days 0 and 21 to induce arthritis. Daily mean clinical scores were calculated for each animal by averaging the daily values given for each paw using the following clinical scoring criteria:

| **Score** | | **Observation** |
|---|---|---|
| 0 | = | Normal |
| 1 | = | One hind or fore paw joint affected or minimal diffuse erythema and swelling |
| 2 | = | Two hind or fore paw joints affected or mild diffuse erythema and swelling |
| 3 | = | Three hind or fore paw joints affected or moderate diffuse erythema and swelling |
| 4 | = | Marked diffuse erythema and swelling, or 4-digit joints affected |
| 5 | = | Severe diffuse erythema and severe swelling entire paw, unable to flex digits |

Upon onset of arthritis between study days 26-28, the animals were randomized by clinical arthritis scores into groups, and treatment was initiated; the day of treatment initiation is defined as arthritis day 1 for graph of mean paw scores. Mice were treated on arthritis days 1 through 14 orally once daily (QD) with either vehicle, bexobrutideg, rilzabrutinib, or ibrutinib; orally twice daily (BID) with tofacitinib; or QD via intraperitoneal injection (IP) with etanercept. On arthritis day 15 animals were humanely euthanized and plasma cells in the spleen were enumerated by flow cytometry.

Spleens were dissociated through a 70 μm filter and red blood cells were removed by incubation with ACK lysing buffer. Washed cells were transferred to 96-well plates and stained with live/dead dye (diluted 1:1000 in PBS) for 10 minutes on ice, spun down, and washed once with FACS buffer (PBS + 2% FBS + 2 mM EDTA). Samples were fixed in 100 μL of 1x Foxp3 fixation and permeabilization solution for 1 hour at room temperature, then washed twice in 1x permeabilization buffer.

Intracellular stain was prepared in 1x permeabilization buffer with the following antibodies: Brilliant Violet 711 anti-mouse IgG3 (1:400), FITC anti-mouse IgA (1:400), PE anti-mouse IgE (1:400), PE-Dazzle 594 anti-mouse IgG1 (1:800), PE-Cy7 anti-mouse IgG2b (1:400), Alexa Fluor 647 anti-mouse IgM (1:400), Alexa Fluor 700 anti-mouse IgD (1:400), APC-Fire 750 anti mouse IgG2a (1:400), and mouse Fc block (1:50). Intracellular stain was applied to cells at 100 μL/well, and samples were incubated at room temperature for 30 minutes. Samples were spun down, washed once with 1x permeabilization buffer, and washed once with FACS buffer. Surface and lineage marker stain was prepared in Brilliant Stain Buffer with the following antibodies: Brilliant Violet 421 anti-mouse CD138 (1:100), Brilliant Violet 605 anti-mouse CD19 (1:100), Brilliant Violet 785 anti-mouse/human CD45R/B220 (1:200), and mouse Fc block (1:50). Surface and lineage stain was applied to cells at 100 uL/well, and samples were incubated at room temperature for 15 minutes. Samples were spun down, washed once with FACS buffer, and resuspended in FACS/2% PFA (1:1 dilution of 4% PFA with FACS buffer). Samples were subsequently analyzed on an Attune NxT Acoustic Focusing Flow Cytometer.

Flow cytometry data were analyzed using FlowJo. Lymphocytes were gated based on forward and side scatter, single cells were gated based on forward scatter height and forward scatter width, and live cells were gated as live/dead dye negative. Plasma cells (PCs) were further gated as CD138+ IgD-, and B cells were gated as B220+ CD19+. Plasma cell counts in each sample were normalized by dividing the total number of plasma cells by the total number of B cells and multiplying by $10^6$. Statistical analysis was performed in GraphPad Prism on all groups receiving anti-type II collagen immunization (groups 2-9). Two ordinary one-way ANOVA analyses were performed with Dunnett's multiple comparisons test: all groups compared to vehicle, and the 30 mg/kg ibrutinib and 30 mg/kg rilzabrutinib groups compared to 30 mg/kg bexobrutideg.

*Passive cutaneous anaphylaxis mouse model*

On day 1, BALB/cJ mice were injected with either PBS (left ear) or anti-dinitrophenyl (DNP) IgE (right ear, 100 ng). On day 2, animals were challenged with 10 mg/kg DNP-human serum albumin (DNP-HSA) and 50 mg/kg Evans blue dye intravenously. On days 1 and 2 treatment groups were administered either vehicle control or bexobrutideg orally once daily, remibrutinib orally twice daily, or the positive control cyproheptadine intraperitoneally once daily. Efficacy was evaluated one hour after antigen challenge on day 2 (at approximately 6 h post final treatment administration) by measuring change in ear thickness (Figure S3C) and vascular permeability via Evans blue dye extravasation in the anti-DNP IgE-sensitized ear. Evans blue was measured by extracting blue dye from the whole ear pinnae. Ears were isolated, cut into small pieces, dissolved in 500 µl DMF, and incubated at 37°C with shaking at 225 RPM. After overnight incubation, 200 µl of clarified DMF was used to measure the concentration of extracted dye at absorbance of 650 nm using a plate reader and normalizing to a standard curve. BTK levels were assessed from the PBS-control ear tissue by Jess Simple Western to calculate %BTK remaining as described for tumor tissues.

*Flow cytometry analysis of non-human primate samples*

Whole venous blood was collected into Cyto-Chex BCT tubes at pre-dose and 2, 4, 8, 12, and 24 h post oral administration of 3 mg/kg bexobrutideg. Fixed whole blood (100 µL) was plated in a 96-well round-bottom plate, washed with 1× PBS (500 × g, 5 min), red blood cells were lysed with ACK lysis buffer, and cells were washed twice with PBS. Surface staining was performed with PE anti-human CD3 and BV421 anti-human CD20 in the presence of mouse and human Fc receptor blocking reagents (20 µg/mL each), followed by two PBS washes. Intracellular BTK staining was performed using a Foxp3 fixation/permeabilization kit with an anti-BTK rabbit primary antibody and an Alexa Fluor 488–conjugated anti-rabbit IgG secondary antibody, with two permeabilization buffer washes between each step. Isotype controls were processed identically, omitting the primary antibody. Cells were fixed in 2% paraformaldehyde/ 1% FBS/1 mM EDTA in PBS and acquired on an Attune NxT Acoustic Focusing Flow Cytometer (Thermo Fisher, cat. no. A29004). Data were analyzed in FlowJo and GraphPad Prism. BTK geometric mean fluorescence intensity (MFI) was determined for B cells ($CD20^{+}CD3^{-}$) and isotype control was used for background subtraction. Percent BTK degradation was calculated for each post-dose sample using the following equation:

$$\%BTK\ Remaining = 100 \times \frac{BTK\ MFI_{Sample} - BTK\ MFI_{Isotype}}{BTK\ MFI_{Pre-dose} - BTK\ MFI_{Isotype}}$$

*In vitro interconversion*

The interconversion (racemization) may occur under physiological conditions for many cereblon (CRBN) harness ligands that contain a chiral glutarimide ring, such as thalidomide and lenalidomide (Mori et al., 2018; Amako et al., 2025). The potential for interconversion in bexobrutideg was assessed by in vitro using pooled cynomolgus monkey and human blood and hepatocytes. Compound **8** is a 1:1 mixture of diastereomers NX-5948 and **9**. Absolute stereochemistry of NX-5948 was determined as (S) at glutarimide stereogenic center. Initial physicochemical results (solution, pH adjustments) suggested that the diastereomers do not interconvert.

Supercritical fluid chromatography (SFC) was used to chromatographically resolve the diastereomers. This isocratic SFC method (details below) was then used to monitor the interconversion potential of each diastereomer in blood or hepatocytes.

SFC method:

Instrument: SFC-PICLABS ANALYTIC 10

Mobile Phase: 55% CO2, 45% (MeOH/Acetonitrile/NH4OH @ 80/20/0.1)

Column: AS-H

Flow rate: 6 grams / min

Detection: UV @ 290 nm

Column Temperature: 40°C

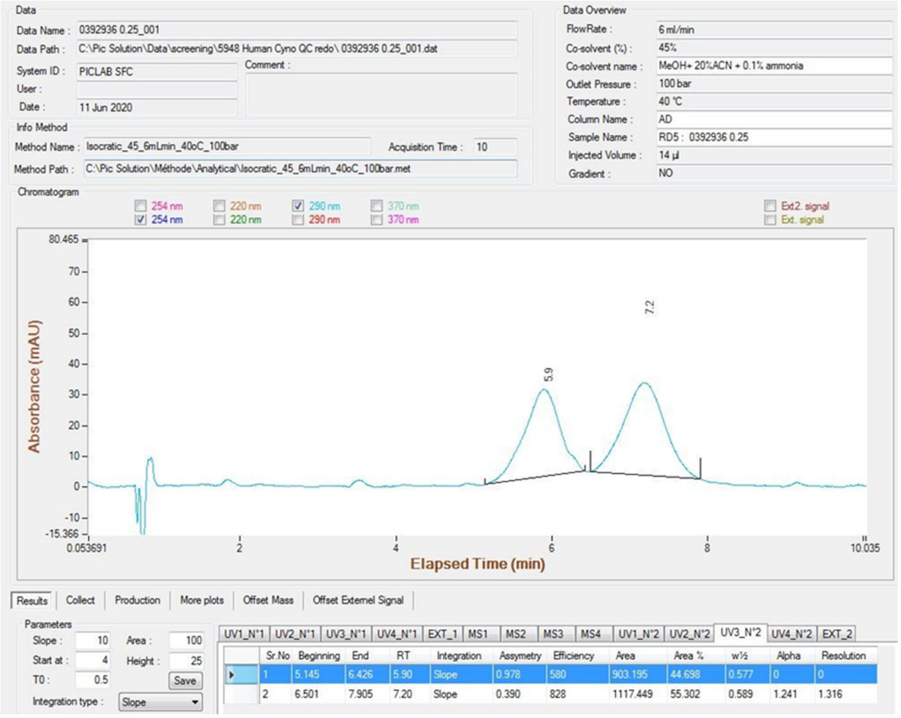


Peak retention times of 5.9 minutes and 7.2 minutes correspond to NX-5948 and **9**, respectively. The method was assessed for concentration linearity and robustness.

In Vitro Materials -

Fresh, pooled mixed gender, human and cynomolgus monkey blood (K2EDTA) was procured from BioIVT. Cryo-preserved pooled mixed gender human and cynomolgus monkey hepatocytes were procured from BioIVT. A 50 mg/mL stock solution of NX-5948 was made in DMSO.

Experimental –

Prewarmed blood and hepatocytes were spiked to 0.1mg/mL final

- 3mLs of blood per timepoint; 0h, 0.25h, 0.5h & 1h @ 37°C
- 2.5 Million hepatocytes (0.5mL) per timepoint; 0h, 0.5h, 1h & 2h @ 37°C
- Incubations were quenched with a 4X volume of acetonitrile + 0.1% formic acid
- Vortexed vigorously for 5 minutes, centrifuged and transferred supernatant
- Evaporated to dryness
- Reconstituted samples with 0.2mL of acetonitrile/water+0.1% formic acid (50/50)
- Inject for SFC-UV

Results:

After a 2h 37 °C incubation of NX-5948 in human hepatocytes, 74% of NX-5948 parent remained and no isomerization to give compound **9** was observed. After a 2h 37 °C incubation of NX-5948 in cynomolgus monkey hepatocytes, 79% of NX-5948 parent remained and no isomerization to give compound **9** was observed. After a 1h 37 °C incubation in human or cynomolgus monkey blood >98% of NX-5948 **9** remained.

Conclusions:

Human and cynomolgus monkey hepatocyte and blood incubations were carried out and analyzed using super critical fluid chromatography (SFC) to assess the possible interconversion of NX-5948 to compound **9**. There was no observed interconversion/racemization at any timepoint.

*Synthetic Procedures*

All reagents and solvents were purchased from commercial sources and used as received. Silica gel and reverse-phase flash column chromatography were conducted with Teledyne ISCO CombiFlash instruments and commercially available pre-packed columns. Reverse-phase preparative high-pressure liquid chromatography (HPLC) purification of final analogs was performed over C18 columns with UV and MS detection using a MeCN/water gradient and TFA modifier. All compounds tested were of ≥95% purity by UPLC-MS. For several compounds, $^1$H NMR integrations are reported as approximate owing to peak broadening, conformational exchange, or overlap with residual solvent signals inherent to the structural complexity of these molecules.

**Synthesis of BTK Binder**

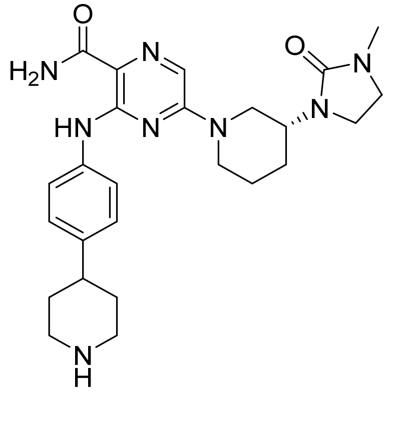

The synthesis of this compound was reported previously (Jia et al., 2015).

### Synthesis of Targeted BTK Degraders

**Example 1**: NRX-0492 (Robbins et al., 2024)

**Example 2**: 3-((4-(1-((1-(1-(2,6-dioxopiperidin-3-yl)-6-oxo-1,6-dihydropyridazin-4-yl)piperidin-4-yl)methyl)piperidin-4-yl)phenyl)amino)-5-((R)-3-(3-methyl-2-oxoimidazolidin-1-yl)piperidin-1-yl)pyrazine-2-carboxamide:

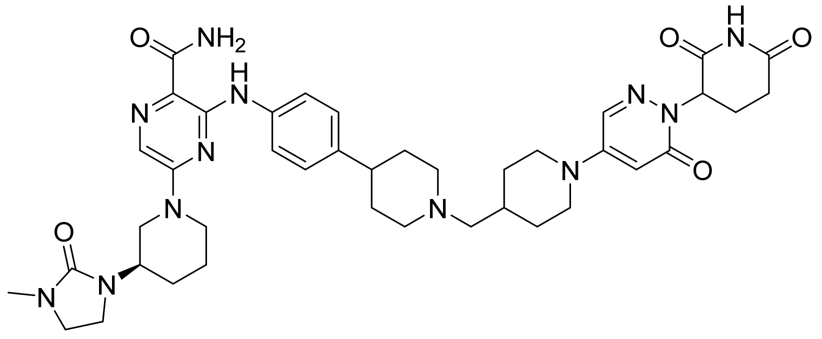

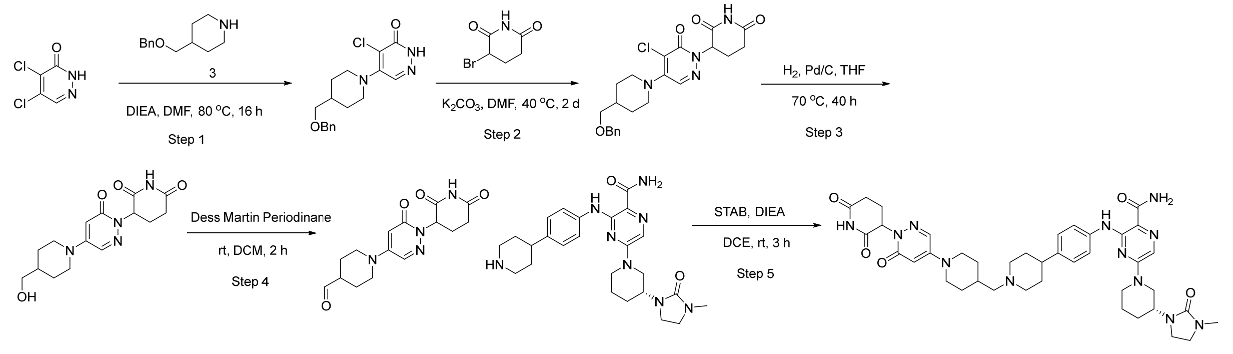


Step 1: 5-(4-(benzyloxymethyl)piperidin-1-yl)-4-chloropyridazin-3(2H)-one

To a solution of 4,5-dichloropyridazin-3(2H)-one (10.0 g, 60.6 mmol) in DMF (100 mL) were added 4-(benzyloxymethyl)piperidine (15.5 g, 75.8 mmol) and DIEA (23.5 g, 182 mmol). The resulting mixture was stirred at 80 ºC for 16 h under nitrogen before being concentrated under vacuum. The residue was triturated with ethyl acetate and dichloromethane to afford the desired product (12.0 g, crude) as a brown solid, which was used in the next step without further purification. MS (ESI) calculated for ($C_{17}H_{20}$ C l$N_3O_2$) $[M+H]^+$, 334.1; found 334.0.

Step 2: 3-(4-(4-(benzyloxymethyl)piperidin-1-yl) -5-chloro-6-oxopyridazin-1(6H)-yl)piperidine-2,6-dione

To a solution of 5-(4-(benzyloxymethyl)piperidin-1-yl)-4-chloropyridazin-3(2H)-one (10.5 g, 31.5 mmol) in DMF (100 mL) were added 3-bromopiperidine-2,6-dione (15.0 g, 78.3 mmol) and

$K_2CO_3$ (15.2 g, 110 mmol). The mixture was stirred at 40 ºC for 24 h under nitrogen. Another portion of 3-bromopiperidine-2,6-dione (7.5 g, 39.3 mmol) and $K_2CO_3$ (7.6 g, 55.1 mmol) was added. The resulting mixture was stirred at 40 ºC for another 24 h under nitrogen. The mixture was diluted with water and extracted with ethyl acetate. The combined organic layers were washed with brine, dried over anhydrous sodium sulfate, and concentrated under vacuum. The residue was purified by reverse phase flash column chromatography with 15~65% MeCN in $H_2O$ to afford the desired product (5.5 g, 39%) as a yellow solid. MS (ESI) calculated for ($C_{22}H_{25}C lN_4O_4$) $[M+H]^+$, 445.2; found 445.1.

Step 3: 3-(4-(4-(hydroxymethyl)piperidin-1-yl)-6-oxopyridazin-1(6H) -yl)piperidine-2,6-dione

To a solution of 3-(4-(4-(benzyloxymethyl)piperidin-1-yl)-5-chloro-6-oxopyridazin-1(6H)-yl) piperidine-2,6-dione (3.30 g, 7.41 mmol) in THF (80 mL) was added 10% Pd/C (dry, 600 mg) under nitrogen. The resulting suspension was stirred at 70 ºC for 40 h under 1 atm hydrogen. The mixture was cooled then filtered. The filtrate was concentrated under vacuum. The residue was purified by Prep-HPLC with the following conditions: [Column: XBridge Shield RP18 OBD Column, 30*150 mm, 5um; Mobile Phase A:Water(10 MMOL/L $NH_4HCO_3$), Mobile Phase B: ACN; Flow rate: 60 mL/min; Gradient: 7 B to 28 B in 7 min; 220 nm] to afford the desired product (485 mg, 20%) as a light yellow solid. MS (ESI) calculated for ($C_{15}H_{20}N_4O_4$) $[M+H]^+$, 321.1; found 321.3. $^1$H NMR (300 MHz, DMSO-*d6*) δ 10.92 (s, 1H), 8.01 (d, *J* = 2.7 Hz, 1H), 5.80 (d, *J* = 2.7 Hz, 1H), 5.56 (dd, *J* = 12.0, 5.4 Hz, 1H), 4.82 (br, 1H), 3.91 – 3.89 (m, 2H), 3.25 – 3.23 (m, 2H), 3.02 – 2.69 (m, 3H), 2.59 – 2.54 (m, 1H), 2.52 – 2.45 (m, 1H), 2.09 – 1.87 (m, 1H), 1.84 – 1.43 (m, 3H), 1.35 – 0.89 (m, 2H).

Step 4: 1-(1-(2,6-dioxopiperidin-3-yl)-6-oxo-1,6-dihydropyridazin-4-yl)piperidine-4-carbaldehyde

Dess Martin periodinane (191 mg, 0.450 mmol) was added to 3-{4-[4-(hydroxymethyl)piperidin-1-yl]-6-oxopyridazin-1-yl}piperidine-2,6-dione (120 mg, 0.375 mmol) in DCM (5 mL). The reaction mixture was stirred for 2 h, then filtered through Celite and concentrated onto silica gel. Purification by silica gel chromatography (0-100% ethyl acetate in hexane) provided the desired product (80 mg, 67%). This material was carried forward without further characterization.

Step 5: 3-((4-(1-((1-(1-((RS)-2,6-dioxopiperidin-3-yl)-6-oxo-1,6-dihydropyridazin-4-yl)piperidin-4-yl)methyl)piperidin-4-yl)phenyl)amino)-5-((R)-3-(3-methyl-2-oxoimidazolidin-1-yl)piperidin-1-yl)pyrazine-2-carboxamide

5-[(3R)-3-(3-methyl-2-oxoimidazolidin-1-yl)piperidin-1-yl]-3-{[4-(piperidin-4-yl)phenyl]amino}pyrazine-2-carboxamide (120 mg, 0.25 mmol), 1-[1-(2,6-dioxopiperidin-3-yl)-

6-oxopyridazin-4-yl]piperidine-4-carbaldehyde (1.2 equiv, 0.96 mg), and DIEA (3 equiv, 0.13 mL) in DCE (10 mL), stirred at rt for 15 minutes. Sonication helped the reaction mixture become homogeneous. Then, sodium triacetoxyborohydride (3 equiv, 0.16 g) was added and the reaction was stirred for 3h. The reaction was then partitioned between MeOH/DCM (10%) and water. The organic layer was separated, washed with brine, dried over magnesium sulfate, and concentrated under reduced pressure. The residue was purified by reverse phase HPLC (0-100% ACN in water) to provide the desired product (54 mg, 28%).

$^1$H NMR (500 MHz, DMSO) δ 11.29 (s, 1H), 10.97 (s, 1H), 9.04 (s, 1H), 8.08 (d, $J$ = 2.9 Hz, 1H), 7.79 (d, $J$ = 2.8 Hz, 1H), 7.68 (s, 1H), 7.60 – 7.54 (m, 2H), 7.36 (d, $J$ = 2.7 Hz, 1H), 7.21 – 7.09 (m, 2H), 5.91 (d, $J$ = 2.8 Hz, 1H), 5.59 (dd, $J$ = 12.1, 5.4 Hz, 1H), 4.37 – 4.27 (m, 2H), 3.98 (d, $J$ = 13.2 Hz, 2H), 3.67 – 3.58 (m, 2H), 3.39 – 3.21 (m, 5H), 3.06 (h, $J$ = 10.6, 9.2 Hz, 5H), 3.02 – 2.74 (m, 4H), 2.72 (d, $J$ = 2.0 Hz, 3H), 2.58 (dt, $J$ = 17.2, 4.1 Hz, 1H), 2.44 (qd, $J$ = 12.8, 4.5 Hz, 1H), 2.19 – 2.14 (m, 1H), 2.04 – 1.88 (m, 4H), 1.86 – 1.73 (m, 5H), 1.58 (dd, $J$ = 12.4, 3.7 Hz, 1H), 1.28 (d, $J$ = 11.6 Hz, 1H), 1.23 (d, $J$ = 12.1 Hz, 1H). MS (ESI) calculated for ($C_{40}H_{52}N_{12}O_5$) $[M+H]^+$, 781.9; found 781.7.

**Example 3**: 3-((4-(1-((1-(2-(2,6-dioxopiperidin-3-yl)-1-oxo-1,2-dihydroisoquinolin-6-yl)piperidin-4-yl)methyl)piperidin-4-yl)phenyl)amino)-5-((R)-3-(3-methyl-2-oxoimidazolidin-1-yl)piperidin-1-yl)pyrazine-2-carboxamide

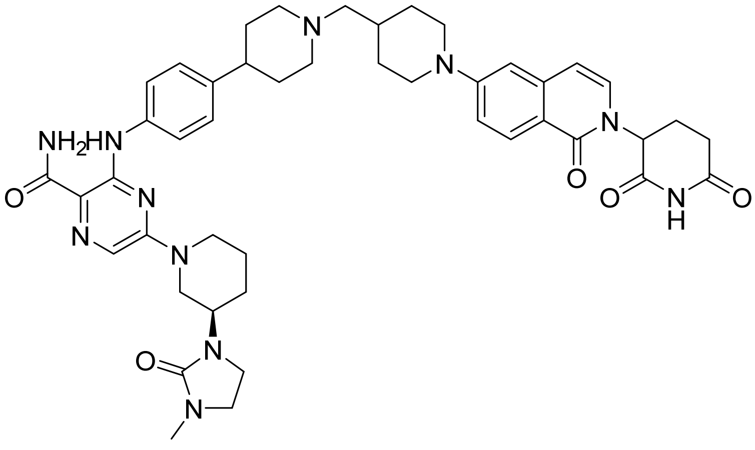

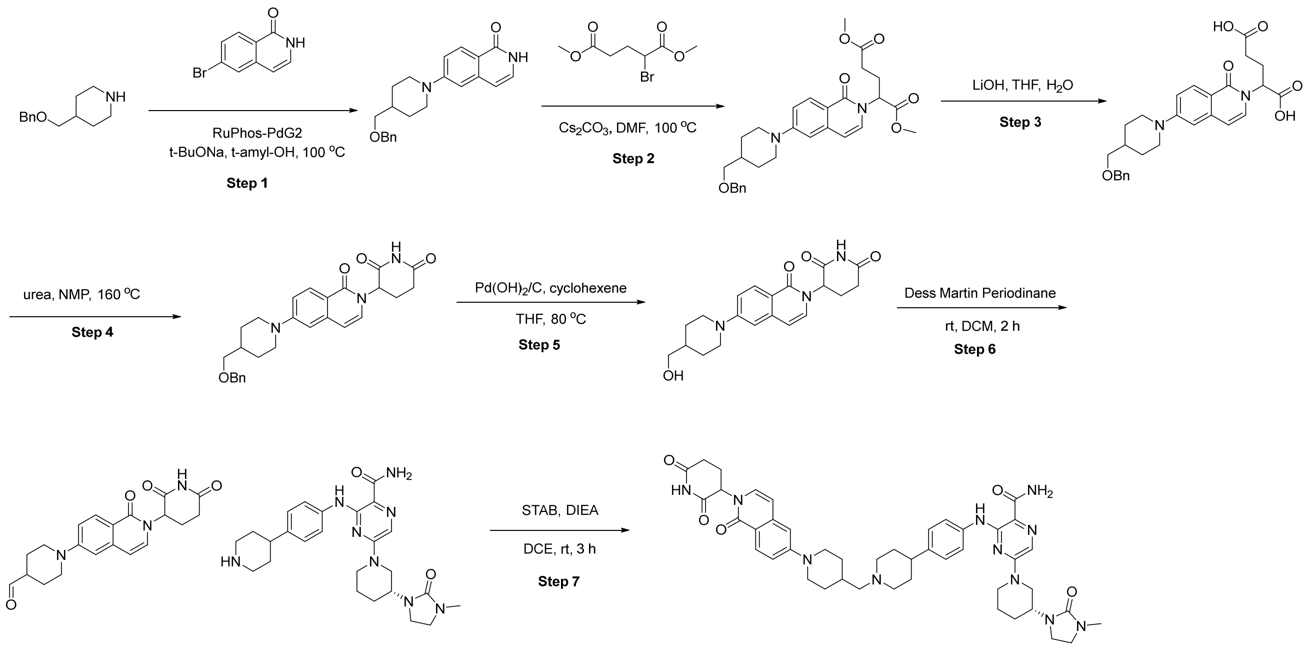

Step 1: 6-(4-(benzyloxymethyl)piperidin-1-yl)isoquinolin-1(2H)-one

To a degassed solution of 6-bromo-2H-isoquinolin-1-one (4.00 g, 17.9 mmol) in t-amyl alcohol (50 mL) were added 4-[(benzyloxy)methyl]piperidine (4.40 g, 21.4 mmol), t-BuONa (5.20 g, 53.9 mmol) and RuPhos-PdCl-2nd G (1.39 g, 1.78 mmol). The mixture was stirred at 100 ºC for 3 h under nitrogen atmosphere. The reaction mixture was quenched by the addition of saturated citric acid aqueous solution and extracted with ethyl acetate. The combined organic layers were washed with brine, dried over anhydrous $Na_2SO_4$ and filtered. The filtrate was concentrated under vacuum. The residue was purified by flash column chromatography with 0~10% methanol in dichloromethane to afford 6-(4-(benzyloxymethyl)piperidin-1-yl)isoquinolin-1(2H)-one (5.5 g, 88%) as a brown solid. MS (ESI) calculated for ($C_{22}H_{24}N_2O_2$) $[M+H]^+$, 349.2; found 349.2.

Step 2: dimethyl 2-(6-(4-(benzyloxymethyl)piperidin-1-yl)-1-oxoisoquinolin-2(1H)-yl)pentanedioate

To a solution of 6-(4-(benzyloxymethyl)piperidin-1-yl)isoquinolin-1(2H)-one (6.20 g, 17.8 mmol) in DMF (60 mL) were added dimethyl 2-bromopentanedioate (5.00 g, 20.9 mmol) and $Cs_2CO_3$ (17.4 g, 53.4 mmol). The resulting mixture was stirred at 100 ºC for 16 h under nitrogen atmosphere. The mixture was diluted with saturated citric acid aqueous solution and extracted with ethyl acetate. The combined organic layers were washed with brine, dried over anhydrous $Na_2SO_4$ and filtered. The filtrate was concentrated under vacuum to afford dimethyl 2-(6-(4-(benzyloxymethyl)piperidin-1-yl)-1-oxoisoquinolin-2(1H)-yl)pentanedioate (6 g, crude) as a brown oil, which was used in the next step without further purification. MS (ESI) calculated for ($C_{29}H_{34}N_2O_6$) $[M+H]^+$, 507.2; found 507.2.

Step-3: Synthesis of 2-(6-(4-(benzyloxymethyl)piperidin-1-yl)-1-oxoisoquinolin-2(1H)-yl)pentanedioic acid

To a solution of 1,5-dimethyl 2-(6-[4-[(benzyloxy)methyl]piperidin-1-yl]-1-oxoisoquinolin-2-yl)pentanedioate (20.0 g, 39.5 mmol) in MeOH (80 mL), THF (80 mL) and $H_2O$ (80 mL) was added LiOH (5.67 g, 237 mmol). The mixture was stirred at room temperature for 16 h. The organic solvents were removed under vacuum and the residue was diluted with water, extracted with ethyl acetate. The collected aqueous layer was acidified to pH 5~6 by citric acid and extracted with ethyl acetate. The combined organic layers were washed with brine, dried over anhydrous $Na_2SO_4$ and filtered. The filtrate was concentrated under vacuum to afford 2-(6-(4-(benzyloxymethyl)piperidin-1-yl)-1-oxoisoquinolin-2(1H)-yl)pentanedioic acid (15 g, crude) as a brown oil, which was used in the next step without further purification. MS (ESI) calculated for ($C_{27}H_{30}N_2O_6$) $[M+H]^+$, 479.2; found 479.0.

Step-4: Synthesis of 3-(6-(4-(benzyloxymethyl)piperidin-1-yl)-1-oxoisoquinolin-2(1H)-yl)piperidine-2,6-dione

To a solution of 2-(6-(4-(benzyloxymethyl)piperidin-1-yl)-1-oxoisoquinolin-2(1H)-yl)pentanedioic acid (1.60 g, 3.34 mmol) in NMP (15 mL) was added urea (2.00 g, 33.3 mmol). The mixture was stirred at 160 ºC for 4 h under nitrogen atmosphere. The resulting mixture was cooled to room temperature and diluted with water. The mixture was extracted with ethyl acetate. The combined organic layers were washed with brine, dried over anhydrous $Na_2SO_4$ and filtered. The filtrate was concentrated under vacuum. The residue was purified by reverse phase flash column chromatography with 5~65% acetonitrile in water to afford 3-(6-(4-(benzyloxymethyl)piperidin-1-yl)-1-oxoisoquinolin-2(1H)-yl)piperidine-2,6-dione (350 mg, 22%) as an off-white solid. MS (ESI) calculated for ($C_{27}H_{29}N_3O_4$) $[M+H]^+$, 460.2; found 460.2.

Step-5: Synthesis of 3-(6-(4-(hydroxymethyl)piperidin-1-yl)-1-oxoisoquinolin-2(1H)-yl)piperidine-2,6-dione

To a solution of 3-(6-(4-(benzyloxymethyl)piperidin-1-yl)-1-oxoisoquinolin-2(1H)-yl)piperidine-2,6-dione (1.80 g, 3.91 mmol) in THF (60 mL) were added $Pd(OH)_2/C$ (10%, 1.8 g) and cyclohexene (3.20 g, 39.0 mmol). The mixture was stirred at 80 ºC for 24 h under nitrogen atmosphere. When the reaction was completed, the solids were filtered, and the filtrate was concentrated under vacuum. The residue was purified by high-pressure reverse phase flash column chromotagraphy with the following conditions: [column, C18 silica gel; mobile phase, MeCN in water (0.1 % ammonium bicarbonate), 15% to 40% gradient in 30 min; detector, UV 254 nm] to afford 3-(6-(4-(hydroxymethyl)piperidin-1-yl)-1-oxoisoquinolin-2(1H)-yl)piperidine-2,6-dione (800 mg, 55%) as an off-white solid. 1H NMR (300 MHz, DMSO-d6) δ 10.72 (s, 1H), 7.94 (d, J = 9.0 Hz, 1H), 7.23 (d, J = 7.5 Hz, 1H), 7.14 (dd, J = 9.0, 2.4 Hz, 1H), 6.92 (d, J = 2.4 Hz, 1H), 6.44 (d, J = 7.5 Hz, 1H), 5.40 (s, 1H), 4.47 (t, J = 5.1 Hz, 1H), 3.97 – 3.94 (m, 2H), 3.27 – 3.24

(m, 2H), 2.85 – 2.78 (m, 3H), 2.58 – 2.52 (m, 2H), 1.98 – 1.94 (m, 1H), 1.81 – 1.67 (m, 2H), 1.65 – 1.59 (m, 1H), 1.26 – 1.12 (m, 2H). MS (ESI) calculated for ($C_{20}H_{23}N_3O_4$) $[M+H]^+$, 370.2; found, 370.3.

Step 6 **General oxidation of alcohol to aldehyde procedure:** Synthesis of 1-(2-(2,6-dioxopiperidin-3-yl)-1-oxo-1,2-dihydroisoquinolin-6-yl)piperidine-4-carbaldehyde

Dess martin periodinane (0.140 g, 0.330 mmol) was added to 3-{6-[4-(hydroxymethyl)piperidin-1-yl]-1-oxoisoquinolin-2-yl}piperidine-2,6-dione (100 mg, 0.270 mmol) in DCM (5 mL) then stirred for 2 h. The reaction was then filtered with Celite and concentrated onto silica gel. Silica gel chromatography (0-100% ethyl acetate in hexane) provided desired product. MS (ESI) calculated for ($C_{20}H_{21}N_3O_4$) $[M+H]^+$, 368.2; found, 368.3.

Step 7 **Reductive amination general procedure**: Synthesis of 3-((4-(1-((1-(2-((RS)-2,6-dioxopiperidin-3-yl)-1-oxo-1,2-dihydroisoquinolin-6-yl)piperidin-4-yl)methyl)piperidin-4-yl)phenyl)amino)-5-((R)-3-(3-methyl-2-oxoimidazolidin-1-yl)piperidin-1-yl)pyrazine-2-carboxamide

5-[(3R)-3-(3-methyl-2-oxoimidazolidin-1-yl)piperidin-1-yl]-3-{[4-(piperidin-4-yl)phenyl]amino}pyrazine-2-carboxamide (15 mg, 0.031 mmol) and 1-[2-(2,6-dioxopiperidin-3-yl)-1-oxoisoquinolin-6-yl]piperidine-4-carbaldehyde (23 mg, 0.063 mmol) were combined in DCE (0.1M) and DIEA (0.017 ml, 0.093 mmol), then stirred for 10 minutes. Then sodium triacetoxyborohydride (20 mg, 0.094 mmol) was added and the reaction was stirred for 3h. The reaction was then partitioned between methanol/DCM (10%) and water, and the organic layer was separated, dried over magnesium sulfate, concentrated, then purified by reverse phase HPLC to provide the desired product. $^1H$ NMR (500 MHz, DMSO-$d_6$) δ 11.21 (s, 1H), 10.99 (s, 1H), 7.97 (d, $J$ = 9.1 Hz, 1H), 7.76 (d, $J$ = 2.7 Hz, 1H), 7.67 (s, 1H), 7.51 (d, $J$ = 8.2 Hz, 2H), 7.34 (d, $J$ = 2.7 Hz, 1H), 7.26 (d, $J$ = 7.5 Hz, 1H), 7.23 (s, 1H), 7.18 (dd, $J$ = 8.8, 2.2 Hz, 3H), 6.96 (d, $J$ = 2.5 Hz, 1H), 6.47 (d, $J$ = 7.5 Hz, 1H), 5.44 (s, 1H), 4.36 (d, $J$ = 12.5 Hz, 1H), 4.29 (d, $J$ = 13.3 Hz, 1H), 3.97 (d, $J$ = 12.6 Hz, 2H), 3.63 (td, $J$ = 11.1, 9.0, 5.4 Hz, 1H), 3.33 – 3.23 (m, 2H), 3.00 (dt, $J$ = 37.8, 11.7 Hz, 3H), 2.92 – 2.80 (m, 3H), 2.73 (s, 3H), 2.62 (s, 1H), 2.62 – 2.56 (m, 1H), 2.29 (s, 0H), 2.05 – 1.97 (m, 1H), 1.86 – 1.72 (m, 7H), 1.70 (s, 4H), 1.62 – 1.52 (m, 1H), 1.19 (dd, $J$ = 20.9, 9.9 Hz, 2H). MS (ESI) calculated for $C_{45}H_{55}N_{11}O_5$ $[M+H]^+$, 830.1; found, 830.7.

**Example 4**: 3-((4-(1-((1-(6-((2,6-dioxopiperidin-3-yl)amino)pyridin-3-yl)piperidin-4-yl)methyl)piperidin-4-yl)phenyl)amino)-5-((R)-3-(3-methyl-2-oxoimidazolidin-1-yl)piperidin-1-yl)pyrazine-2-carboxamide

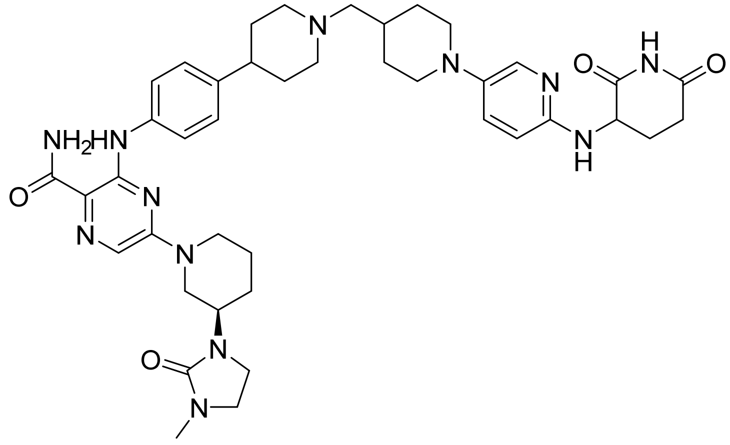

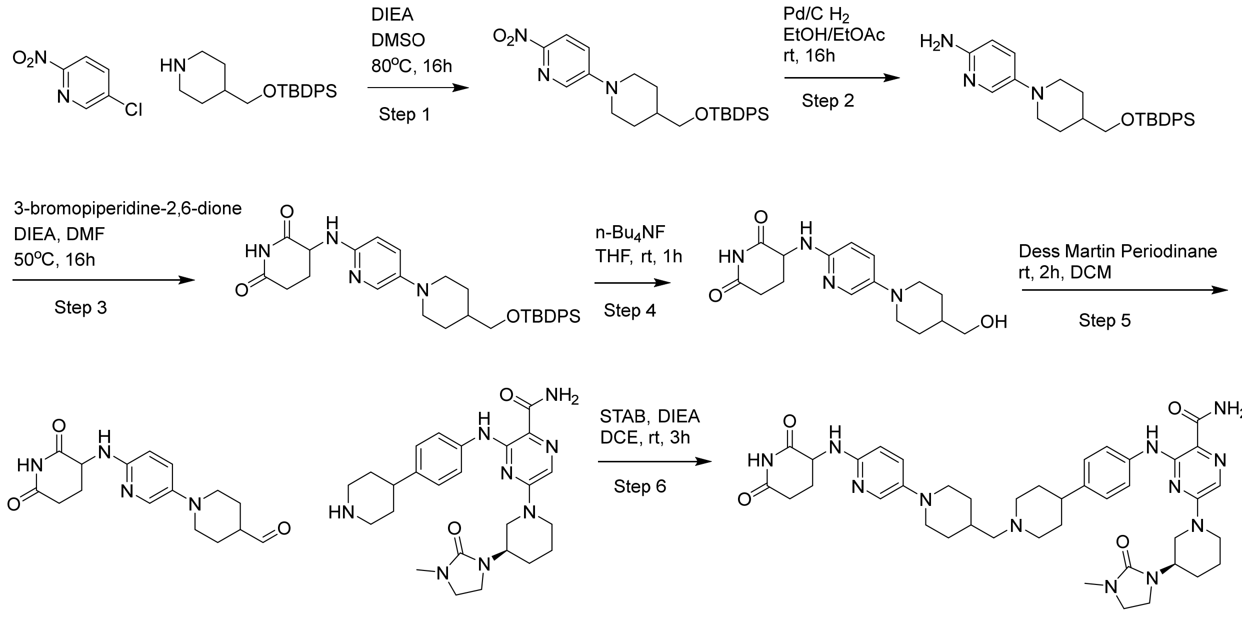

Step 1: 5-(4-(((tert-butyldiphenylsilyl)oxy)methyl)piperidin-1-yl)-2-nitropyridine

To a solution of *tert*-butyl-diphenyl-(4-piperidylmethoxy)silane (8.8 g, 25 mmol) and 5-chloro-2-nitro-pyridine (4.0 g, 25 mmol) in DMSO (20 mL) was added *i*-$Pr_2NEt$ (13 mL, 76 mmol). After stirring at 80 °C for 16 h, the reaction mixture was cooled to RT then diluted with DCM and water. The aqueous phase was extracted with DCM, washed with saturated aqueous NaCl, dried over $Na_2SO_4$, and concentrated under reduced pressure to provide the desired product (11 g, 92%). MS (ESI) calculated for $C_{27}H_{33}N_3OSi$ $[M+H]^+$, 476; found, 476.

Step 2: 5-(4-(((tert-butyldiphenylsilyl)oxy)methyl)piperidin-1-yl)pyridin-2-amine

A solution of *tert*-butyl-[[1-(6-nitro-3-pyridyl)-4-piperidyl]methoxy]-diphenyl-silane (12.0 g, 25.2 mmol) and 10% Pd/C (1.34 g) in EtOH (50 mL) and EtOAc (50 mL) was stirred under $H_2$ for 16 h. The reaction mixture was filtered through Celite and concentrated under reduced pressure. The residue was purified by flash chromatography ($SiO_2$) eluting with 0–10%

MeOH–DCM to provide the desired product (8.2 g, 73%) as a solid. This material was taken directly into the next step without further characterization.

Step 3: 3-((5-(4-(((tert-butyldiphenylsilyl)oxy)methyl)piperidin-1-yl)pyridin-2-yl)amino)piperidine-2,6-dione

To a solution of 5-[4-[[*tert*-butyl(diphenyl)silyl]oxymethyl]-1-piperidyl]pyridin-2-amine (8.2 g, 18.4 mmol) and 3-bromopiperidine-2,6-dione (4.24 g, 22.1 mmol) in DMF (20 mL) was added *i*-$Pr_2NEt$ (6.4 mL, 36.8 mmol). After stirring at 50 °C for 16 h, the reaction mixture was concentrated under reduced pressure. The residue was purified by flash chromatography ($SiO_2$, 0–50% MeOH–DCM) to provide the desired product (4.1 g, 40%). MS (ESI) calculated for $C_{32}H_{40}N_4O_3Si$ $[M+H]^+$, 557; found, 557.

Step 4: 3-((5-(4-(hydroxymethyl)piperidin-1-yl)pyridin-2-yl)amino)piperidine-2,6-dione

To a solution of 3-[[5-[4-[[*tert*-butyl(diphenyl)silyl]oxymethyl]-1-piperidyl]-2-pyridyl]amino]piperidine-2,6-dione (4.1 g, 7.4 mmol) in THF (20 mL) was added *n*-$Bu_4NF$ (11 mL, 11 mmol, 1 M in THF). After stirring for 1 h, the reaction mixture was concentrated under reduced pressure. Flash chromatography ($SiO_2$, 2–12% MeOH–DCM) followed by reverse phase chromatography (C18, 10-40% MeCN–$H_2O$/1% $HCO_2H$) provided the desired product (325 mg, 14%). $^1$H NMR (500 MHz, DMSO-$d_6$) δ 10.72 (s, 1H), 7.64 (d, $J$ = 3.0 Hz, 1H), 7.22 (dd, $J$ = 9.0, 3.0 Hz, 1H), 6.54 (d, $J$ = 9.0 Hz, 1H), 6.37 (d, $J$ = 7.6 Hz, 1H), 4.65 (ddd, $J$ = 12.5, 7.5, 5.1 Hz, 1H), 4.44 (t, $J$ = 5.4 Hz, 1H), 3.40 – 3.33 (m, 2H), 3.28 (t, $J$ = 5.8Hz, 2H), 2.74 (ddd, $J$ = 17.5, 13.1, 5.5 Hz, 1H, 2.57– 2.51 (m, 2H), 2.49 – 2.44 (m, 1H), 2.12 – 2.06 (m, 1H), 1.96 (qd, $J$ = 12.8, 4.5 Hz, 1H), 1.77 – 1.69 (m, 2H), 1.48 – 1.38 (m, 1H), 1.26 (dd, $J$ = 12.1 Hz, 1H), 1.21 (dd, $J$ = 12.1, 3.6 Hz, 1H). MS (ESI) calculated for $C_{16}H_{22}N_4O_3$ $[M+H]^+$, 319; found, 319.

Step 5: 1-(6-((2,6-dioxopiperidin-3-yl)amino)pyridin-3-yl)piperidine-4-carbaldehyde

Synthesized following the oxidation procedure described above starting from 3-({5-[4-(hydroxymethyl)piperidin-1-yl]pyridin-2-yl}amino)piperidine-2,6-dione (100 mg, 0.31 mmol) to afford the desired product (46 mg, 0 46%). MS (ESI) calculated for $C_{16}H_{20}N_4O_3$ $[M+H]^+$, 317; found, 317.

Step 6: 3-((4-(1-((1-(6-((2,6-dioxopiperidin-3-yl)amino)pyridin-3-yl)piperidin-4-yl)methyl)piperidin-4-yl)phenyl)amino)-5-((R)-3-(3-methyl-2-oxoimidazolidin-1-yl)piperidin-1-yl)pyrazine-2-carboxamide

Synthesized following the general procedure of reductive amination described above starting from *tert*-butyl 4-[4-({3-carbamoyl-6-[(3R)-3-(3-methyl-2-oxoimidazolidin-1-yl)piperidin-1-yl]pyrazin-2-yl}amino)phenyl]piperidine-1-carboxylate (41.3 mg, 0.0714 mmol) and 1-{6-[(2,6-dioxopiperidin-3-yl)amino]pyridin-3-yl}piperidine-4-carbaldehyde (22.6 mg,

0.0720 mmol) to afford the desired product (47.1 mg, 83%) $^1$H NMR (500 MHz, $CD_3CN$) δ 11.10 (s, 1H), 9.12 (s, 1H), 8.80 (s, 1H), 7.81 (d, J = 9.5 Hz, 1H), 7.59 (d, J = 8.0 Hz, 2H), 7.55 (s, 1H), 7.41 (s, 1H), 7.37 (s, 1H), 7.19 (d, J = 7.9 Hz, 2H), 6.96 (d, J = 9.7 Hz, 1H), 5.81 (s, 1H), 4.51 (dd, J = 12.3, 5.1 Hz, 1H), 4.39 (d, J = 12.9 Hz, 1H), 4.29 (d, J = 13.7 Hz, 1H), 3.71 – 3.62 (m, 3H), 3.52 (d, J = 11.9 Hz, 2H), 3.42 – 3.33 (m, 1H), 3.35 – 3.28 (m, 1H), 3.27 (dd, J = 10.5, 6.9 Hz, 2H), 3.06 (t, J = 11.7 Hz, 1H), 2.98 (dd, J = 15.5, 9.8 Hz, 4H), 2.82 (s, 3H), 2.75 (d, J = 1.7 Hz, 4H), 2.75 – 2.65 (m, 3H), 2.27 (dd, J = 13.0, 4.9 Hz, 1H), 2.19 – 2.10 (m, 1H), 2.10 – 2.03 (m, 3H), 1.97 – 1.89 (m, 3H), 1.88 (s, 1H), 1.80 (dd, J = 24.4, 12.9 Hz, 2H), 1.64 (d, J = 12.5 Hz, 1H), 1.43 (d, J = 12.7 Hz, 2H). MS (ESI) calculated for $C_{41}H_{54}N_{12}O_4$ $[M+H]^+$, 780; found, 780.

**Example 5**: Synthesis of 3-((4-(1-((1-(5-((2,6-dioxopiperidin-3-yl)(methyl)amino)pyridin-2-yl)piperidin-4-yl)methyl)piperidin-4-yl)phenyl)amino)-5-((R)-3-(3-methyl-2-oxoimidazolidin-1-yl)piperidin-1-yl)pyrazine-2-carboxamide:

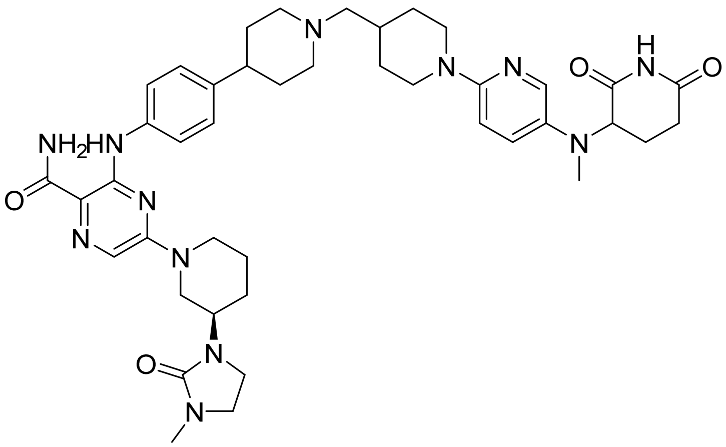

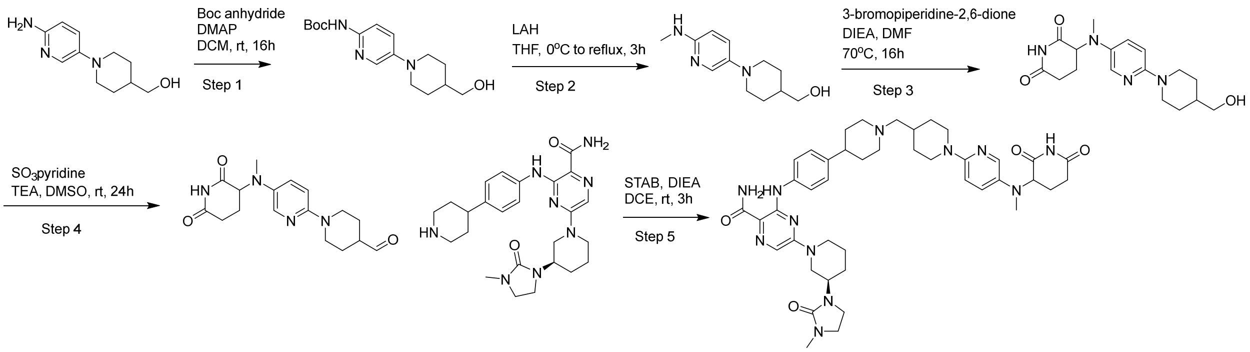


Step 1: Tert-butyl N-[6-[4-(hydroxymethyl)-1-piperidyl]-3-pyridyl]carbamate

To a mixture of [1-(5-amino-2-pyridyl)-4-piperidyl]methanol (5.0 g, 31.4 mmol) and DMAP (192 mg, 1.57 mmol) in anhydrous DCM (20 mL) at rt, was added a solution of *tert*-butoxycarbonyl tert-butyl carbonate (7.53 g, 34.5 mmol) in anhydrous DCM (10 mL) and the resulting mixture was stirred at rt for 16 h. The mixture was diluted with water (100 mL) and the layers were separated. The aqueous layer was extracted with DCM (3 x 20 mL and the combined organic layers were washed with brine (2 x 20 mL) then dried over anhydrous $Na_2SO_4$, filtered and concentrated under reduced pressure to afford title compound (5.3 g, 55%) as a solid, which was used in the next step without further purification. MS (ESI) calculated for $C_{16}H_{25}N_3O_3$ [M-tBu+H]$^+$ ,252.1; found, 252.1.

Step 2: [1-[5-(methylamino)-2-pyridyl]-4-piperidyl]methanol

To a solution of *tert*-butyl N-[6-[4-(hydroxymethyl)-1-piperidyl]-3-pyridyl]carbamate (2.2 g, 7.16 mmol) in anhydrous THF (20 mL) at 0 °C, was added $LiAlH_4$ (14.3 mL, 14.3 mmol, 2 M in THF) and the resulting suspension was warmed to rt then refluxed for 3 h. The mixture was cooled to rt, $Na_2SO_4.10H_2O$ was added and the mixture was stirred for 15 min. The suspension was filtered, and the volatiles were evaporated under reduced pressure. The material was purified by silica gel column chromatography (120 g) using a gradient of 5-15% MeOH in DCM to afford title compound (1.01 g, 62%) as a solid. MS (ESI) calculated for $C_{12}H_{19}N_3O$ [M+H]$^+$ 222.1; found, 222.1.

Step 3: 3-[[6-[4-(hydroxymethyl)-1-piperidyl]-3-pyridyl]amino]piperidine-2,6-dione

To a mixture of [1-[5-(methylamino)-2-pyridyl]-4-piperidyl]methanol (2.8 g, 12.7 mmol) and 3-bromopiperidine-2,6-dione (4.9 g, 25 mmol) in anhydrous DMF (20 mL) at rt, was added DIPEA (6.6 mL, 38.0 mmol) and the resulting mixture was heated at 70 °C for 16 h. The volatiles were evaporated under reduced pressure, and the material was purified by silica gel column chromatography (120 g) using a gradient of 3-12% MeOH in DCM. The material was further purified by reverse phase chromatography on C18 (120 g) using 10-30% MeCN and water (containing 1% formic acid) to afford desired product (1.1 g, 26%) as a gray solid. $^1$H NMR (500 MHz, DMSO-$d_6$) δ 10.73 (s, 1H), 7.79 (d, J = 3.1 Hz, 1H), 7.20 (dd, J = 9.2, 3.2 Hz, 1H), 6.74 (d, J = 9.1 Hz, 1H), 4.66 (dd, J = 12.6, 4.9 Hz, 1H), 4.44 (s, 1H), 4.08 (d, J = 12.8 Hz, 2H), 3.27 (d, J = 6.3 Hz, 2H), 2.84 – 2.74 (m, 1H), 2.70 (s, 3H), 2.66 – 2.57 (m, 2H), 2.57 – 2.52 (m, 1H), 2.31 – 2.19 (m, 1H), 1.91 – 1.83 (m, 1H), 1.75 – 1.65 (m, 2H), 1.58 – 1.48 (m, 1H), 1.20 – 1.08 (m, 2H). MS (ESI) calculated for $C_{16}H_{22}N_4O_3$ [M+H]$^+$ 319.2; found, 319.2.

Step 4: 1-(5-((2,6-dioxopiperidin-3-yl)(methyl)amino)pyridin-2-yl)piperidine-4-carbaldehyde

To a mixture of 3-({6-[4-(hydroxymethyl)piperidin-1-yl]pyridin-3-yl}(methyl)amino)piperidine-2,6-dione (100 mg, 0.30 mmol) and triethylamine (0.17 mL, 0.12 g, 1.20 mmol) in DMSO (1.50 mL) was added sulfur trioxide pyridine complex (96 mg, 0.60 mmol). After 24 h, the reaction was loaded directly onto a reverse phase ISCO 50 g C18 column which was eluted with 0 to 50% ACN/water gradient to provide 1-{5-[(2,6-dioxopiperidin-3-

yl)(methyl)amino]pyridin-2-yl}piperidine-4-carbaldehyde (0.0314 g, 31.6%). MS (ESI) calculated for $C_{16}H_{20}N_4O_3$ $[M+H]^+$ 317.2; found, 317.2.

Step 5: 3-((4-(1-((1-(5-((2,6-dioxopiperidin-3-yl)(methyl)amino)pyridin-2-yl)piperidin-4-yl)methyl)piperidin-4-yl)phenyl)amino)-5-((R)-3-(3-methyl-2-oxoimidazolidin-1-yl)piperidin-1-yl)pyrazine-2-carboxamide

Followed the general procedure of reductive amination described above starting from *tert*-butyl 4-[4-({3-carbamoyl-6-[(3R)-3-(3-methyl-2-oxoimidazolidin-1-yl)piperidin-1-yl]pyrazin-2-yl}amino)phenyl]piperidine-1-carboxylate (22 mg, 0.045 mmol) and 1-{5-[(2,6-dioxopiperidin-3-yl)(methyl)amino]pyridin-2-yl}piperidine-4-carbaldehyde (15 mg, 0.045mmol) to provide 3-((4-(1-((1-(5-((2,6-dioxopiperidin-3-yl)(methyl)amino)pyridin-2-yl)piperidin-4-yl)methyl)piperidin-4-yl)phenyl)amino)-5-((R)-3-(3-methyl-2-oxoimidazolidin-1-yl)piperidin-1-yl)pyrazine-2-carboxamide (27 mg, 75% yield).
$^1$H NMR (500 MHz, DMSO-$d_6$) δ 11.19 (s, 1H), 10.75 (s, 1H), 7.83 – 7.74 (m, 2H), 7.67 (s, 1H), 7.50 (d, $J$ = 8.2 Hz, 2H), 7.34 (d, $J$ = 2.9 Hz, 1H), 7.24 – 7.15 (m, 3H), 6.75 (d, $J$ = 9.2 Hz, 1H), 4.67 (dd, $J$ = 12.5, 4.8 Hz, 1H), 4.38 (d, $J$ = 12.3 Hz, 1H), 4.30 (d, $J$ = 13.4 Hz, 1H), 4.08 (d, $J$ = 12.5 Hz, 2H), 3.67 – 3.59 (m, 1H), 3.31 – 3.24 (m, 3H), 3.10 – 2.90 (m, 4H), 2.85 – 2.60 (m, 11H), 2.30 – 2.22 (m, 1H), 2.18 (d, $J$ = 6.9 Hz, 2H), 1.98 (t, $J$ = 11.4 Hz, 2H), 1.93 – 1.50 (m, 13H), 1.19 – 1.08 (m, 2H). MS (ESI) calculated for $C_{42}H_{56}N_{12}O_4$ $[M+H]^+$ 793.5; found, 793.7.

**Example 6**: 3-((4-(1-((1-(5-((2,6-dioxopiperidin-3-yl)oxy)pyridin-2-yl)piperidin-4-yl)methyl)piperidin-4-yl)phenyl)amino)-5-((R)-3-(3-methyl-2-oxoimidazolidin-1-yl)piperidin-1-yl)pyrazine-2-carboxamide

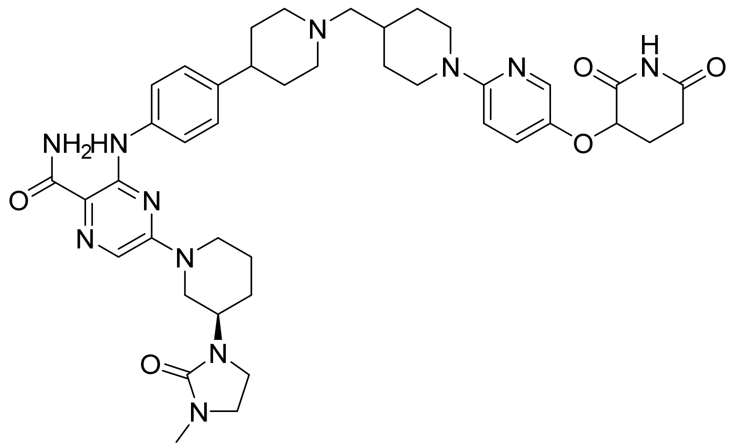

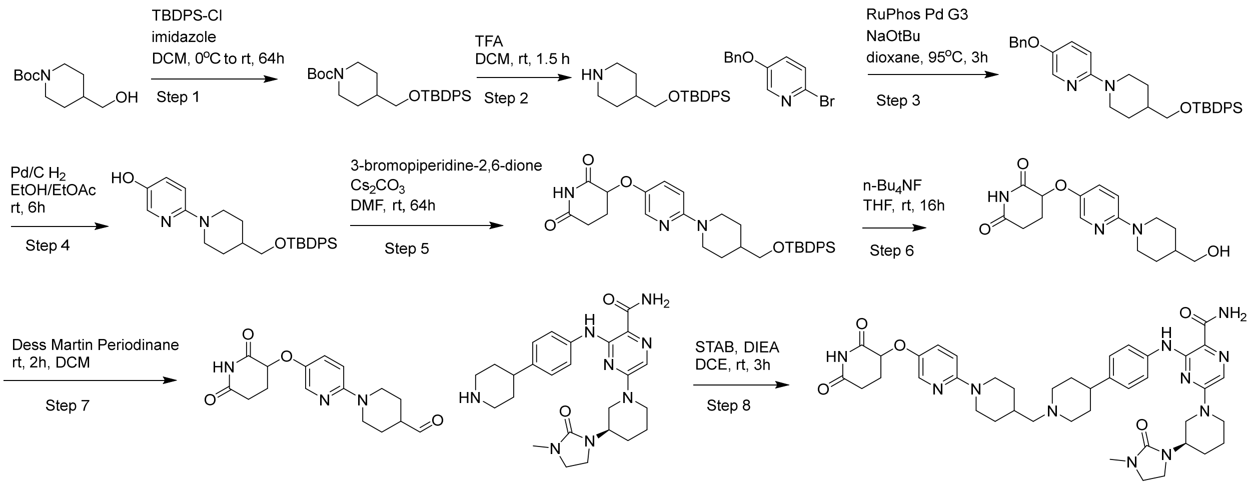

Step 1: tert-butyl 4-(((tert-butyldiphenylsilyl)oxy)methyl)piperidine-1-carboxylate

TBDPSCl (27.5 g, 100 mmol) was added dropwise to a solution of *tert*-butyl 4-(hydroxymethyl)piperidine-1-carboxylate (21.5 g, 100 mmol) and imidazole (8.17 g, 120 mmol) in DCM (250 mL) at 0 °C. After stirred at RT for 64 h, the reaction mixture was washed with $H_2O$, saturated aqueous NaCl, dried over $MgSO_4$, and concentrated under reduced pressure to provide the desired product (46.4 g, quant.). $^1H$ NMR (400 MHz, $CDCl_3$) δ 7.69 – 7.62 (m, 4H), 7.47 – 7.34 (m, 6H), 4.10 (s, 2H), 3.49 (d, $J$ = 5.8 Hz, 2H), 2.69 (t, $J$ = 12.9 Hz, 2H), 1.74 – 1.63 (m, 3H), 1.45 (s, 9H), 1.21 – 1.10 (m, 2H), 1.05 (s, 9H). MS (ESI) calculated for $C_{27}H_{39}NO_3Si$ $[M+H]^+$ 454; found, 454.

Step 2: 4-(((tert-butyldiphenylsilyl)oxy)methyl)piperidine

TFA (50 mL, 673 mmol) was added to a solution of *tert*-butyl 4-(((*tert*-butyldiphenylsilyl)oxy)methyl)piperidine-1-carboxylate (45.4 g, 100 mmol) in DCM (200 mL). After stirring for 1.5 h, the reaction mixture was concentrated under reduced pressure to provide the desired compound (34.5 g, 97.5 mmol, 98%). $^1H$ NMR (400 MHz, $CDCl_3$) δ 7.69 – 7.62 (m, 4H), 7.46 – 7.34 (m, 6H), 3.49 (d, $J$ = 6.1 Hz, 2H), 3.09 (dt, $J$ = 12.1, 3.0 Hz, 2H), 2.60 (td, $J$ = 12.2, 2.7 Hz, 2H), 1.78 – 1.59 (m, 3H), 1.22 – 1.11 (m, 2H), 1.05 (s, 9H). MS (ESI) calculated for $C_{22}H_{31}NOSi$ $[M+H]^+$ 354; found, 354.

Step 3: 5-(benzyloxy)-2-(4-(((tert-butyldiphenylsilyl)oxy)methyl)piperidin-1-yl)pyridine

A suspension of 4-(((*tert*-butyldiphenylsilyl)oxy)methyl)piperidine (7.85 g, 22.2 mmol), 5-benzyloxy-2-bromopyridine (4.89 g, 18.5 mmol), RuPhos Pd G3 (774 mg, 0.925 mmol) and NaO*t*Bu (5.33 g, 55.5 mmol) in dioxane (90 mL) was degassed and backfilled with $N_2$. After stirring at 95 °C for 3 h, the reaction mixture was cooled to RT and concentrated under reduced pressure. Flash chromatography ($SiO_2$, 0–10% EtOAc–hexanes) provided the desired product (9.70 g, 18.1 mmol, 81%). $^1$H NMR (400 MHz, $CDCl_3$) δ 7.98 (d, *J* = 3.1 Hz, 1H), 7.75 – 7.62 (m, 4H), 7.51 – 7.28 (m, 11H), 7.18 (dd, *J* = 9.1, 3.1 Hz, 1H), 6.63 (d, *J* = 9.1 Hz, 1H), 5.02 (s, 2H), 4.19 – 4.08 (m, 2H), 3.54 (d, *J* = 6.2 Hz, 2H), 2.75 (td, *J* = 12.6, 2.8 Hz, 2H), 1.88 – 1.79 (m, 2H), 1.79 – 1.69 (m, 1H), 1.33 (qd, *J* = 12.4, 4.1 Hz, 2H), 1.06 (s, 9H). MS (ESI) calculated for $C_{34}H_{40}N_2O_2Si$ $[M+H]^+$ 537; found, 537.

Step 4: 6-(4-(((tert-butyldiphenylsilyl)oxy)methyl)piperidin-1-yl)pyridin-3-ol

A solution of 5-(benzyloxy)-2-(4-(((*tert*-butyldiphenylsilyl)oxy)methyl)piperidin-1-yl)pyridine (9.7 g, 18.1 mmol) in EtOAc (45 mL) and EtOH (45 mL) was stirred under 1 atm $H_2$ in the presence of 10% Pd/C (1.92 g) for 6 h. The reaction mixture was filtered through Celite, and concentrated to provide the title desired product (7.84 g, 97%). MS (ESI) calculated for $C_{27}H_{34}N_2O_2Si$ $[M+H]^+$ 447; found, 447.

Step 5: 3-((6-(4-(((tert-butyldiphenylsilyl)oxy)methyl)piperidin-1-yl)pyridin-3-yl)oxy)piperidine-2,6-dione

A suspension of 6-(4-(((*tert*-butyldiphenylsilyl)oxy)methyl)piperidin-1-yl)pyridin-3-ol (7.84 g, 17.6 mmol), 3-bromopiperidine-2,6-dione (5.06 g, 26.3 mmol), and $Cs_2CO_3$ (11.4 g, 35.1 mmol) in DMF (70 mL) was stirred at RT for 64 h. The reaction mixture was diluted with EtOAc and $H_2O$. The aqueous layer was extracted with EtOAc, washed with $H_2O$, saturated aqueous NaCl, dried over $MgSO_4$, and concentrated under reduced pressure. Flash chromatography ($SiO_2$, 0–40% EtOAc–hexanes) provided the desired product (2.67 g, 27%). MS (ESI) calculated for $C_{32}H_{39}N_3O_4Si$ $[M+H]^+$ 558; found, 558.

Step 6: 3-((6-(4-(hydroxymethyl)piperidin-1-yl)pyridin-3-yl)oxy)piperidine-2,6-dione

*n*-$Bu_4NF$ (5.5 mL, 5.5 mmol, 1M in THF) was added to a solution of 3-((6-(4-(((*tert*-butyldiphenylsilyl)oxy)methyl)piperidin-1-yl)pyridin-3-yl)oxy)piperidine-2,6-dione (2.67 g, 4.79 mmol) in THF (25 mL). After stirring at RT for 16 h, the reaction mixture was concentrated under reduced pressure. Flash chromatography ($SiO_2$, 0–10% MeOH–DCM) provided the desired product (924 mg, 2.9 mmol, 61%). $^1$H NMR (400 MHz, DMSO-$d_6$) δ 10.90 (s, 1H), 7.91 (d, *J* = 3.1 Hz, 1H), 7.31 (dd, *J* = 9.2, 3.1 Hz, 1H), 6.78 (d, *J* = 9.2 Hz, 1H), 4.98 (dd, *J* = 10.6, 5.0 Hz, 1H), 4.44 (t, *J* = 5.4 Hz, 1H), 4.20 – 4.09 (m, 2H), 3.26 (t, *J* = 5.6 Hz, 2H), 2.76 – 2.55 (m, 4H), 2.26 – 2.15 (m, 1H), 2.15 – 2.03 (m, 1H), 1.75 – 1.64 (m, 2H), 1.63 – 1.49 (m, 1H), 1.15 (dd, *J* = 12.5, 3.8 Hz, 1H), 1.08 (dd, *J* = 11.8, 4.0 Hz, 1H). MS (ESI) calculated for $C_{16}H_{21}N_3O_4$ $[M+H]^+$ 320; found, 320.

Step 7: 1-(5-((2,6-dioxopiperidin-3-yl)oxy)pyridin-2-yl)piperidine-4-carbaldehyde

Synthesized following the procedure of oxidation described above starting from 3-({6-[4-(hydroxymethyl)piperidin-1-yl]pyridin-3-yl}oxy)piperidine-2,6-dione (200 mg, 0.63 mmol) to afford the desired product (192 mg, 0.61 mmol, 97%). MS (ESI) calculated for $C_{16}H_{19}N_3O_4$ $[M+H]^+$ 318; found, 318.

Step 8: 3-((4-(1-((1-(5-((2,6-dioxopiperidin-3-yl)oxy)pyridin-2-yl)piperidin-4-yl)methyl)piperidin-4-yl)phenyl)amino)-5-((R)-3-(3-methyl-2-oxoimidazolidin-1-yl)piperidin-1-yl)pyrazine-2-carboxamide

Synthesized following the general procedure of reductive amination described above starting from *tert*-butyl 4-[4-({3-carbamoyl-6-[(3R)-3-(3-methyl-2-oxoimidazolidin-1-yl)piperidin-1-yl]pyrazin-2-yl}amino)phenyl]piperidine-1-carboxylate (62.7 mg, 0.11 mmol) and 1-{5-[(2,6-dioxopiperidin-3-yl)oxy]pyridin-2-yl}piperidine-4-carbaldehyde (34.4 mg, 0.11 mmol) to afford the desired product (44 mg, 52%). $^1H$ NMR (500 MHz, DMSO-$d_6$) δ 11.28 (d, J = 7.6 Hz, 1H), 10.94 (s, 1H), 8.93 (s, 1H), 7.91 (d, J = 3.0 Hz, 1H), 7.78 (s, 1H), 7.68 (d, J = 2.2 Hz, 1H), 7.56 (d, J = 8.0 Hz, 2H), 7.46 (d, J = 9.2 Hz, 1H), 7.35 (s, 1H), 7.16 (dd, J = 13.4, 8.1 Hz, 2H), 6.96 (d, J = 9.3 Hz, 1H), 5.05 (dd, J = 10.7, 5.0 Hz, 1H), 4.30 (d, J = 20.5 Hz, 2H), 4.18 (d, J = 12.9 Hz, 2H), 3.62 (d, J = 11.5 Hz, 3H), 3.34 (tt, J = 15.5, 8.8 Hz, 2H), 3.26 (t, J = 7.8 Hz, 2H), 3.12 – 3.01 (m, 4H), 2.97 (t, J = 12.4 Hz, 1H), 2.85 (t, J = 12.4 Hz, 1H), 2.77 (d, J = 12.4 Hz, 1H), 2.71 (s, 3H), 2.65 (dt, J = 13.7, 5.3 Hz, 2H), 2.21 (dd, J = 11.6, 6.4 Hz, 1H), 2.17 – 2.08 (m, 2H), 2.03 – 1.87 (m, 4H), 1.82 (d, J = 12.4 Hz, 4H), 1.76 (s, 1H), 1.57 (d, J = 12.6 Hz, 1H), 1.26 (q, J = 12.3, 11.9 Hz, 2H). MS (ESI) calculated for $C_{41}H_{53}N_{11}O_5$ $[M+H]^+$ 781; found, 781.

**Example 7**: 3-((4-(1-((1-(4-((2,6-dioxopiperidin-3-yl)carbamoyl)phenyl)piperidin-4-yl)methyl)piperidin-4-yl)phenyl)amino)-5-((R)-3-(3-methyl-2-oxoimidazolidin-1-yl)piperidin-1-yl)pyrazine-2-carboxamide

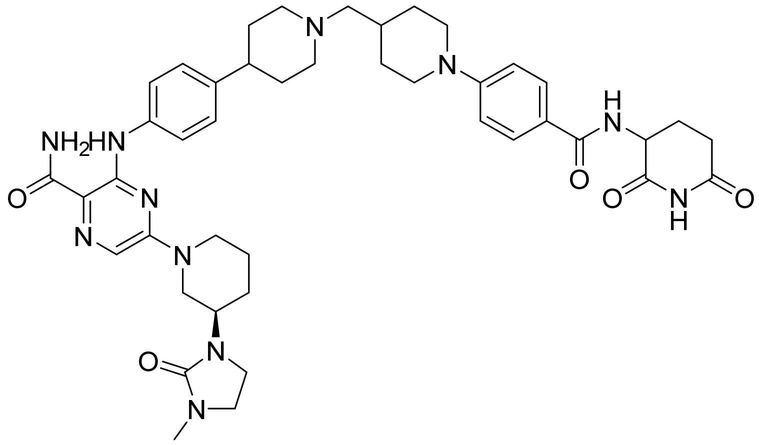

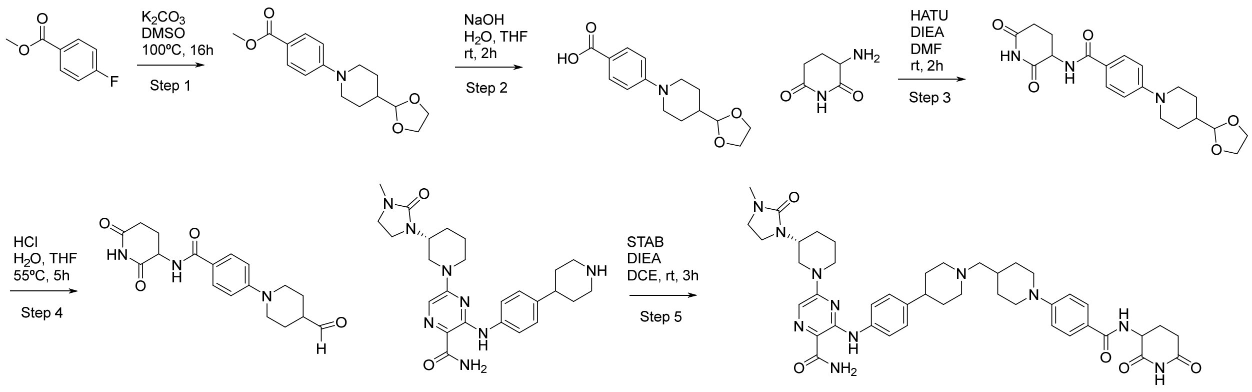


Step 1: methyl 4-[4-(1,3-dioxolan-2-yl)-1-piperidyl]benzoate

$K_2CO_3$ (7.04 g, 50.9 mmol) was added to a mixture of 4-(1,3-dioxolan-2-yl)piperidine (8.00 g, 50.9 mmol) and methyl 4-fluorobenzoate (7.85 g, 50.9 mmol) in anhydrous DMSO (20.0 mL) at 23 °C. The mixture was stirred for at 100 °C for 16 h and diluted with ice water (200 mL). The mixture was filtered, washing with $H_2O$ (100 mL), and the solid was dried to provide the title compound as a solid (13.8 g, 93%). MS (ESI) calculated for $C_{16}H_{21}NO_4$ $[M+H]^+$ 292.2; found, 292.2.

Step 2: 4-[4-(1,3-dioxolan-2-yl)-1-piperidyl]benzoic acid

NaOH (35.0 mL, 175 mmol, 5.00 M in $H_2O$) was added to a mixture of methyl 4-[4-(1,3-dioxolan-2-yl)-1-piperidyl]benzoate (10.0 g, 34.3 mmol) in $H_2O$/THF (200 mL, 1:1) at 23 °C. The mixture was stirred at 23 °C for 2 h and diluted with EtOAc (100 mL). The pH was adjusted to 4 with 1.00 M HCl. The organic layer was separated, and the aqueous layer was extracted with EtOAc (4 X 100 mL). The combined organic layers were washed with brine (200 mL), dried over $Na_2SO_4$, filtered and concentrated to provide the title compound as a solid (7.10 g, 75%). MS (ESI) calculated for $C_{15}H_{19}NO_4$ $[M+H]^+$ 278.1; found, 278.2.

Step 3: 4-[4-(1,3-dioxolan-2-yl)-1-piperidyl]-N-(2,6-dioxo-3-piperidyl)benzamide

DIPEA (3.58 mL, 20.6 mmol) was added to a mixture of 4-[4-(1,3-dioxolan-2-yl)-1-piperidyl]benzoic acid (2.50 g, 9.01 mmol), HATU (6.86 g, 18.0 mmol) and 3-aminopiperidine-2,6-dione;hydrochloride (1.48 g, 9.01 mmol) in anhydrous DMF (20 mL) at 23 °C. The mixture was stirred at 23 °C for 2 h and diluted with $H_2O$ (200 mL). The mixture was extracted with 2-propanol/$CHCl_3$ (4 X 100 mL, 10% 2-propanol in $CHCl_3$), and the combined organic layer was washed with brine (200 mL), dried ($Na_2SO_4$), filtered, and concentrated. The residue was triturated with diethyl ether (200 mL) to provide the title compound as a solid (1.60 g, 44%). MS (ESI) calculated for $C_{20}H_{25}N_3O_5$ $[M+H]^+$ 388.2; found, 388.3.

Step 4: *N*-(2,6-dioxo-3-piperidyl)-4-(4-formyl-1-piperidyl)benzamide

HCl (10.0 mL, 30.0 mmol, 3.00 M in $H_2O$) was added to a mixture of 4-[4-(1,3-dioxolan-2-yl)-1-piperidyl]-*N*-(2,6-dioxo-3-piperidyl)benzamide (1.60 g, 4.13 mmol) in THF (16.0 mL) and $H_2O$ (60.0 mL) at 22 °C. The mixture was stirred at 55 °C for 5 h and diluted with $NaHCO_3$ (1.91 g, 22.7 mmol) in $H_2O$ (200 mL) at 0 °C. The mixture was extracted with 10% 2-propanol/$CHCl_3$ (8 X 100 mL) and the combined organic layers were washed with brine (300 mL), dried ($Na_2SO_4$), filtered, and concentrated. The residue was triturated with diethyl ether (200 mL) to provide the title compound as a solid (277 mg, 20%). $^1$H NMR (500 MHz, DMSO-*d6*) δ 10.82 (s, 1H), 9.64 (s, 1H), 8.46 (d, J = 8.4 Hz, 1H), 7.75 (d, J = 9.0 Hz, 2H), 6.99 (d, J = 9.1 Hz, 2H), 4.78 – 4.70 (m, 1H), 3.81 – 3.74 (m, 2H), 3.03 – 2.92 (m, 2H), 2.84 – 2.72 (m, 1H), 2.62 – 2.52 (m, 2H), 2.17 – 2.05 (m, 1H), 2.01 – 1.86 (m, 3H), 1.62 – 1.50 (m, 2H). MS (ESI) calculated for $C_{18}H_{21}N_3O_4$ $[M+H]^+$ 344.2; found, 344.2.

Step 5: 3-((4-(1-((1-(4-((2,6-dioxopiperidin-3-yl)carbamoyl)phenyl)piperidin-4-yl)methyl)piperidin-4-yl)phenyl)amino)-5-((R)-3-(3-methyl-2-oxoimidazolidin-1-yl)piperidin-1-yl)pyrazine-2-carboxamide

Synthesized following the general procedure of reductive amination described above starting from (R)-5-(3-(3-methyl-2-oxoimidazolidin-1-yl)piperidin-1-yl)-3-((4-(piperidin-4-yl)phenyl)amino)pyrazine-2-carboxamide (30.0 mg, 0.0627 mmol) and N-(2,6-dioxopiperidin-3-yl)-4-(4-formylpiperidin-1-yl)benzamide (21.5 mg, 0.0627 mmol) to afford the desired product (12.7 mg, 25 %). $^1$H NMR (500 MHz, DMSO-*d6*) δ 11.12 (s, 1H), 10.75 (s, 1H), 8.36 (d, *J* = 8.3 Hz, 1H), 7.75 – 7.63 (m, 3H), 7.59 (s, 1H), 7.46 – 7.39 (m, 2H), 7.30 – 7.24 (m, 1H), 7.10 (d, *J* = 8.6 Hz, 2H), 6.92 – 6.85 (m, 2H), 4.67 (ddd, *J* = 12.9, 8.3, 5.4 Hz, 1H), 4.36 – 4.17 (m, 2H), 3.80 (d, *J* = 12.5 Hz, 2H), 3.63 – 3.49 (m, 1H), 3.21 – 3.16 (m, 2H), 3.03 – 2.83 (m, 4H), 2.78 – 2.61 (m, 6H), 2.39 – 2.33 (m, 1H), 2.26 – 1.82 (m, 8H), 1.82 – 1.63 (m, 8H), 1.63 – 1.44 (m, 4H), 1.18 – 1.05 (m, 2H). MS (ESI) calculated for $C_{43}H_{55}N_{11}O_5$ $[M+H]^+$ 806.4; found, 806.7.

**Synthesis of (R)-N-(2,6-dioxopiperidin-3-yl)-5-(4-formylpiperidin-1-yl)picolinamide and (S) -N-(2,6-dioxopiperidin-3-yl)-5-(4-formylpiperidin-1-yl)picolinamide**

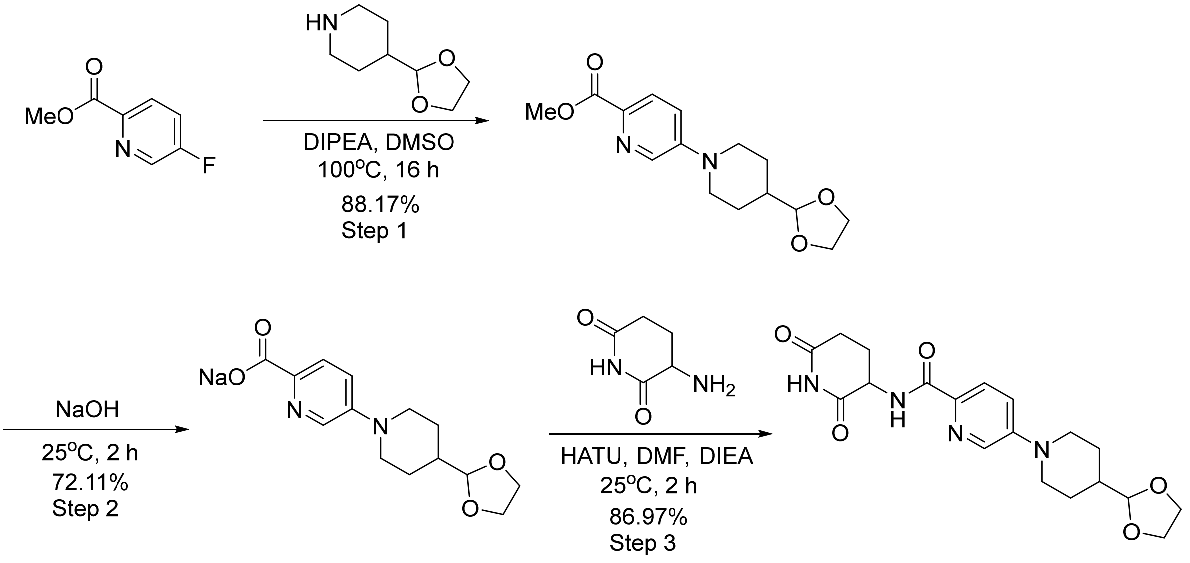


Step 1: methyl 5-(4-(1,3-dioxolan-2-yl)piperidin-1-yl)picolinate

Into a 2-L 3-necked round-bottom flask, was placed methyl 5-fluoropyridine-2-carboxylate (65.0 g, 419 mmol), 4-(1, 3-dioxolan-2-yl)piperidine (65.9 g, 419 mmol), DIPEA (108 g, 838 mmol), and DMSO (650 mL). The resulting solution was stirred for 16 h at 100ºC in an oil bath. The reaction was then quenched by the addition of water (2 L). The solids were collected by filtration to provide the desired product (108 g, 88%) as a yellow solid. MS (ESI) calculated for $C_{15}H_{20}N_2O_4$ $[M+H]^+$ 293; found, 293.

Step 2: 5-(4-(1,3-dioxolan-2-yl)piperidin-1-yl)picolinic acid

Into a 1-L 3-necked round-bottom flask was placed methyl 5-[4-(1,3-dioxolan-2-yl)piperidin-1-yl]pyridine-2- carboxylate (108 g, 369 mmol), caustic soda (18.5 g, 462 mmol), $H_2O$ (100 mL), and THF (500 mL). The resulting solution was stirred for 2 h at 25ºC in a water bath. The solids were collected by filtration to provide the desired product (80 g, 72%) as a brown solid. MS (ESI) calculated for $C_{14}H_{18}N_2O_4$ $[M+H]^+$ 279; found, 279.

Step 3: 5-(4-(1,3-dioxolan-2-yl)piperidin-1-yl)-N-(2,6-dioxopiperidin-3-yl)picolinamide

Into a 2-L 3-necked round-bottom flask, was placed 5-(4-(1,3-dioxolan-2-yl)piperidin-1-yl)picolinic acid (80.0 g, 266 mmol), 3-aminopiperidine-2, 6-dione (37.6 g, 293 mmol), DMF (800 mL), HATU (203 g, 533 mmol), DIEA (79.2 g, 613 mmol). The resulting solution was stirred for 2 h at 25ºC in a water bath. The reaction was then quenched by the addition of water (1 L). The resulting solution was extracted with DCM (3 x 1 L). The combined organic layers were washed with water (2 x 1 L), washed with brine (500 ml), and dried over $Na_2SO_4$ then concentrated under vacuum to provide the desired product (90 g, 87%) as a brown solid.

Step 4: Resolution of racemic mixture to enantiopure intermediates (R)- and (S)-5-(4-(1,3-dioxolan-2-yl)piperidin-1-yl)-N-(2,6-dioxopiperidin-3-yl)picolinamide

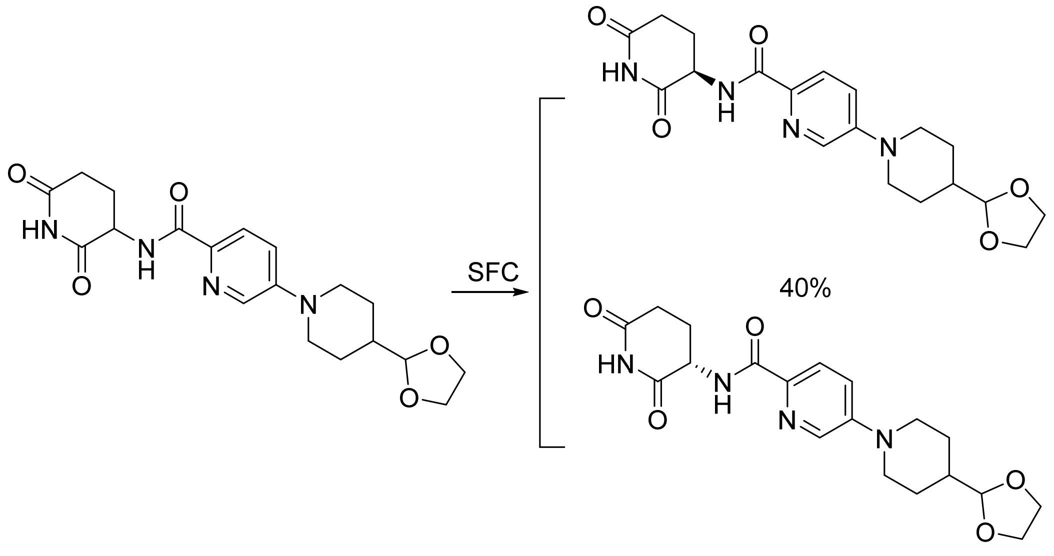


The crude product (90 g) was purified by Prep-SFC with the following conditions (NB-Prep SFC80-1): Column, CHIRAL ART Cellulose-SC, 2*25cm, 5um; mobile phase, $CO_2$ (50%) and ACN: THF=1:1(50%); Detector, UV:254 nm. This resulted in 36 g of 5-[4-(1,3-dioxolan-2-yl)piperidin-1-yl]-N-[(3R)-2,6- dioxopiperidin-3-yl]pyridine-2-carboxamide as a light yellow solid and 35.5 g of 5-[4-(1,3-dioxolan-2-yl) piperidin-1-yl] -N-[(3S)-2, 6-dioxopiperidin-3-yl]pyridine-2-carboxamide as a light yellow solid.

(R): MS (ESI) calculated for $C_{19}H_{24}N_4O_5$ $[M+H]^+$ 389; found, 389.

$^1$HNMR: (400 MHz, DMSO-*d6*): δ10.84 (s, 1H), 8.70 (d, 1H), 8.30 (d, 1H), 7.84 (d, 1H), 7.40 (dd, 1H), 4.74 (ddd, 1H), 4.61 (d, 1H), 3.98 (d, 2H), 3.93–3.72 (m, 4H), 2.84 (td, 3H), 2.82–2.72 (m, 1H), 2.56–2.51 (m, 1H), 2.18 (qd, 1H), 1.83–1.71 (m, 3H), 1.45–1.31 (m, 2H)

(S): MS (ESI) calculated for $C_{19}H_{24}N_4O_5$ $[M+H]^+$ 389; found, 389.

$^1$HNMR: (400 MHz, DMSO-*d6*): δ10.84 (s, 1H), 8.70 (d, 1H), 8.30 (d, 1H), 7.84 (d, 1H), 7.40 (dd, 1H), 4.74 (ddd, 1H), 4.61 (d, 1H), 3.98 (d, 2H), 3.93–3.72 (m, 4H), 2.84 (td, 3H), 2.83–2.72 (m, 1H), 2.54 (d, 1H), 2.18 (qd, 1H), 1.83–1.70 (m, 3H), 1.38 (td, 2H)

Step 5: Deprotection of aldehyde to provide (R) or (S)-N-(2,6-dioxopiperidin-3-yl)-5-(4-formylpiperidin-1-yl)picolinamide

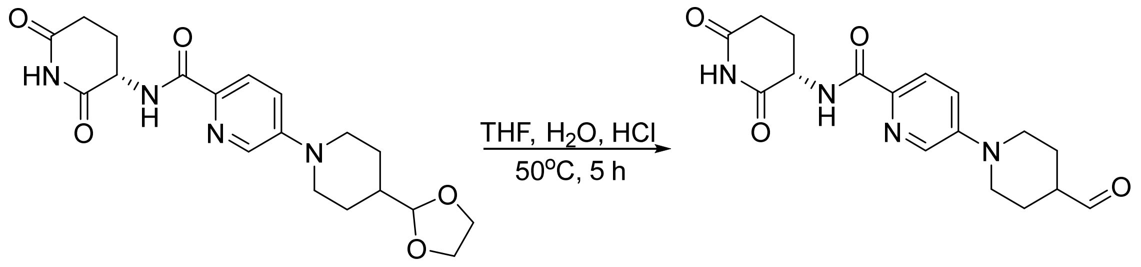

*The S-isomer is described in the main manuscript. The opposite enantiomer (R) was synthesized by an analogous procedure*

(R)-N-(2,6-dioxopiperidin-3-yl)-5-(4-formylpiperidin-1-yl)picolinamide

$^1$HNMR (400 MHz, DMSO-$d_6$): δ10.84 (s, 1H), 9.63 (s, 1H), 8.72 (d, 1H), 8.32 (d, 1H), 7.85 (d, 1H), 7.43 (dd, 1H), 4.74 (ddd, 1H), 3.84 (dt, 2H), 3.05 (ddd, 2H), 2.79 (ddd, 1H), 2.59 (ddt, 2H), 2.18 (qd, 1H), 2.10–1.92 (m, 3H), 1.58 (dtd, 2H). MS (ESI) calculated for $C_{17}H_{20}N_4O_4$ $[M+H]^+$ 345; found, 345.

**Example 8**: 3-((4-(1-((1-(6-((2,6-dioxopiperidin-3-yl)carbamoyl)pyridin-3-yl)piperidin-4-yl)methyl)piperidin-4-yl)phenyl)amino)-5-((R)-3-(3-methyl-2-oxoimidazolidin-1-yl)piperidin-1-yl)pyrazine-2-carboxamide

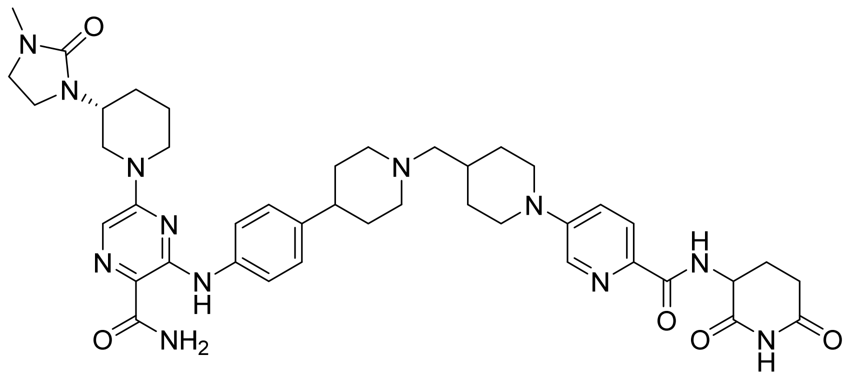

5-[(3R)-3-(3-methyl-2-oxoimidazolidin-1-yl)piperidin-1-yl]-3-{[4-(piperidin-4-yl)phenyl]amino}pyrazine-2-carboxamide (7.00 g, 14.6 mmol) was suspended in 400 mL of $CH_2Cl_2$ and N,N-diisopropylethylamine (5.10 mL, 3.78 g, 29.3 mmol) was added before adding N-(2,6-dioxopiperidin-3-yl)-5-(4-formylpiperidin-1-yl)pyridine-2-carboxamide (5.04 g, 14.6 mmol) in one portion at room temperature. The mixture was sonicated for 2 h owing to the poor solubility of the amine starting material. Sodium triacetoxyborohydride (3.87 g, 18.3 mmol) was added in one portion and the reaction mixture was stirred for 1 h before more aldehyde was added (400 mg) to convert all of the remaining amine to product. After 2 h of stirring the reaction mixture was added to a separatory funnel and $H_2O$ and $CH_2Cl_2$ were added (500 mL each) along with 10 mL MeOH. The organic phase was washed (2 x $H_2O$ 500 mL, 1 x $NaHCO_3$ 200 mL, 1 x brine 400 mL) before being dried ($Na_2SO_4$) filtered and concentrated. The crude solids (11 g) were suspended in ACN (450 mL) and the mixture was heated with stirring to 80 °C for 4 h before being cooled to 50 °C. The solids were removed by filtration to provide the desired product (8.38 g, 70%). $^1$H NMR (500 MHz, DMSO-*d*6) δ 11.29 (s, 1H), 10.85 (s, 1H), 8.91 (s, 1H), 8.70 (d, *J* = 8.2 Hz, 1H), 8.34 (s, 1H), 7.87 (d, *J* = 8.8 Hz, 1H), 7.78 (s, 1H), 7.68 (s, 1H), 7.57 (d, *J* = 8.0 Hz, 2H), 7.49 – 7.39 (m, 1H), 7.35 (s, 1H), 7.18 (d, *J* = 8.1 Hz, 2H), 4.84 – 4.65 (m, 1H), 4.40 – 4.20 (m, 2H), 4.00 (d, *J* = 12.7 Hz, 2H), 3.72 – 3.54 (m, 3H), 3.28 – 3.21 (m, 4H), 3.06 (d, *J* = 9.7 Hz, 5H), 2.93 (q, *J* = 13.3, 12.4 Hz, 3H), 2.78 (d, *J* = 12.3 Hz, 2H), 2.71 (s, 3H), 2.17 (dd, *J* = 21.1, 9.0 Hz, 2H), 2.08 – 1.96 (m, 3H), 1.96 – 1.70 (m, 6H), 1.67 – 1.47 (m, 2H), 1.44 – 1.24 (m, 2H). MS (ESI) calculated for $C_{42}H_{54}N_{12}O_5$ $[M+H]^+$ 807.7; found, 807.7.

**Example 9**: 3-((4-(1-((1-(6-(((R)-2,6-dioxopiperidin-3-yl)carbamoyl)pyridin-3-yl)piperidin-4-yl)methyl)piperidin-4-yl)phenyl)amino)-5-((R)-3-(3-methyl-2-oxoimidazolidin-1-yl)piperidin-1-yl)pyrazine-2-carboxamide

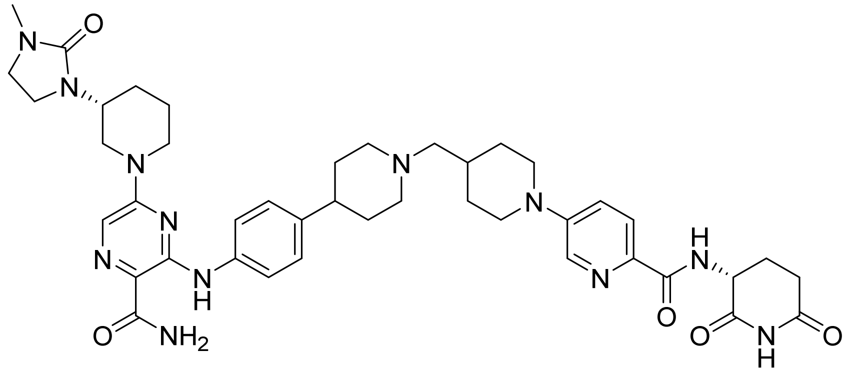

Synthesized following general reductive amination procedure as described above, with (R)-N-(2,6-dioxopiperidin-3-yl)-5-(4-formylpiperidin-1-yl)picolinamide to provide desired product (24 mg, 39% yield). $^1$H NMR (500 MHz, DMSO-*d*6) δ 11.30 (s, 1H), 10.86 (s, 1H), 8.71 (d, J = 8.1 Hz, 1H), 8.36 (s, 1H), 7.88 (d, J = 8.7 Hz, 1H), 7.79 (s, 1H), 7.69 (s, 1H), 7.58 (d, J = 7.9 Hz, 2H), 7.46 (d, J = 8.8 Hz, 1H), 7.36 (s, 1H), 7.19 (d, J = 8.2 Hz, 2H), 4.75 (ddd, J = 13.1, 8.2, 5.4 Hz, 1H), 4.42 – 4.25 (m, 2H), 4.01 (d, J = 12.9 Hz, 2H), 3.65 (dd, J = 13.8, 7.9 Hz, 3H), 3.27 (t, J = 8.3 Hz, 3H), 3.17 – 2.87 (m, 6H), 2.86 – 2.70 (m, 5H), 2.27 – 2.10 (m, 2H), 2.11 – 1.69 (m, 12H), 1.68 – 1.47 (m, 2H), 1.30 (d, J = 52.1 Hz, 3H). MS (ESI) calculated for $C_{42}H_{54}N_{12}O_5$ $[M+H]^+$ 807.4; found, 807.9.

**Example 10 (bexobrutideg, NX-5948)**: 3-{[4-(1-{[1-(6-{[(3S)-2,6-dioxopiperidin-3-yl]carbamoyl}pyridin-3-yl)piperidin-4-yl]methyl}piperidin-4-yl)phenyl]amino}-5-[(3R)-3-(3-methyl-2-oxoimidazolidin-1-yl)piperidin-1-yl]pyrazine-2-carboxamide

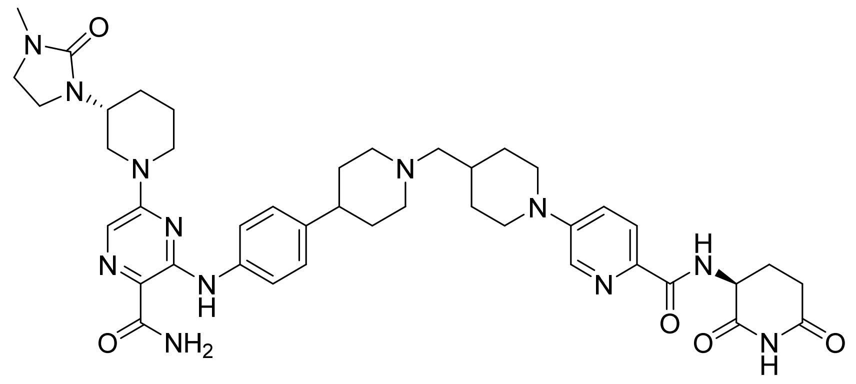

5-[(3R)-3-(3-methyl-2-oxoimidazolidin-1-yl)piperidin-1-yl]-3-{[4-(piperidin-4-yl)phenyl]amino}pyrazine-2-carboxamide (5.00 g, 10.4 mmol) and N-[(3S)-2,6-dioxopiperidin-3-yl]-5-(4-formylpiperidin-1-yl)pyridine-2-carboxamide (3.60g, 10.4 mmol) in DMSO (100 mL) and $CH_2Cl_2$ (400 mL), acetic acid (0.030 mL, 0.52 mmol) was added and stirred for 30 min. Sodium triacetoxyborohydride (4.43g, 20.9 mmol) was added in one portion and the reaction mixture was stirred for 2 h resulting in clarification of the solution. The reaction mixture was poured into a separatory funnel with $CH_2Cl_2$ (300 mL), saturated aq. $NaHCO_3$ (200 mL), $H_2O$ (200 mL), and MeOH (50 mL). The organic layer was collected. The aqueous layer was extracted with 200 mL $CH_2Cl_2$ /50 mL MeOH. The combined organic layers were washed with brine (300 mL), then washed three times with saturated aq. $NaHCO_3$ (300 mL), and washed with brine (400 mL). The organic solution was dried over anhydrous $Na_2SO_4$, filtered, and concentrated to afford a pale yellow solid (8.2 g). The crude material was triturated with ACN (200 mL) at reflux for 3 h then cooled to 40 ºC and filtered to provide desired product (5.91g, 69%). 1H NMR (500 MHz, DMSO-$d_6$) δ 11.20 (s, 1H), 10.86 (s, 1H), 8.71 (d, J = 8.2 Hz, 1H), 8.31 (d, J = 2.9 Hz, 1H), 7.85 (d, J = 8.8 Hz, 1H), 7.76 (d, J = 2.9 Hz, 1H), 7.66 (s, 1H), 7.50 (d, J = 8.6 Hz, 2H), 7.40 (dd, J = 9.0, 2.9 Hz, 1H), 7.34 (d, J = 2.9 Hz, 1H), 7.17 (d, J = 8.6 Hz, 2H), 4.75 (ddd, J = 12.5, 8.2, 5.3 Hz, 1H), 4.37 (d, J = 12.5 Hz, 1H), 4.29 (d, J = 13.3 Hz, 1H), 3.94 (d, J = 13.0 Hz, 2H), 3.62 (ddt, J = 11.0, 8.2, 4.1 Hz, 1H), 3.40 – 3.23 (m, 2H), 3.30 – 3.24 (m, 2H), 3.02 (t, J = 11.7 Hz, 1H), 2.98 (s, 1H), 2.95 – 2.91 (m, 2H), 2.86 (t, J = 12.9 Hz, 2H), 2.80 (ddd, J = 17.2, 13.6, 5.5 Hz, 1H), 2.73 (s, 3H), 2.57 – 2.52 (m, 1H), 2.42 (tt, J = 12.0, 3.8 Hz, 1H), 2.25 – 2.14 (m, 3H), 2.07 – 2.00 (m, 1H), 1.97 (td, J = 11.7, 2.6 Hz, 2H), 1.87 – 1.69 (m, 8H), 1.67 – 1.58 (m, 2H), 1.60 – 1.51 (m, 1H), 1.19 (q, J = 11.9 Hz, 2H). $^{13}C$ NMR (500 MHZ, DMSO-$d_6$) **δ** 173.0, 172.5, 169.2, 164.1, 160.3, 153.5, 150.5, 148.4, 140.0, 138.1, 137.3, 135.1, 127.0, 122.7, 120.6, 119.8, 118.2, 114.2, 64.1, 54.3, 49.3, 48.6, 46.9, 46.7, 45.0, 44.4, 41.3, 38.5, 33.4, 32.7, 31.1, 31.0, 29.6, 27.7, 24.2, 24.0. LCMS: MS (ESI) calculated for $C_{42}H_{54}N_{12}O_5$ $[M+H]^+$, 807.4; found, 807.9

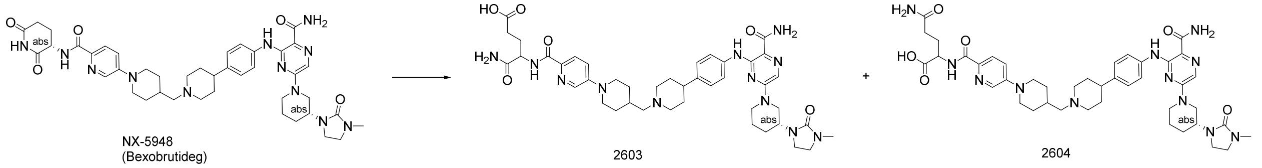

NX-5948 (Bexobrutideg) 2603 2604

two regioisomers after hydrolysis

NX-5948 (21.0 g, 26.01 mmol) was suspended in a mixture of EtOH (70 mL) and THF (350 mL), and 2 M aqueous NaOH (13.0 mL, 26.01 mmol) was added. The resulting suspension was stirred at room temperature and monitored by HPLC until complete consumption of starting material was observed. The reaction mixture remained as a suspension throughout. The pH of the mixture was adjusted from 10.5 to 6.8 by addition of 2 M aqueous HCl, and the suspension was maintained after pH adjustment. The solid was collected by filtration, washed with EtOH (210 mL), and dried under vacuum at 40 °C to afford a mixture of 2603 and 2604 as a solid (18.5 g, 85% yield).

Above crude product (1.16 grams) was purified by Prep-SFC with the following conditions: Column: Princeton SFC-Diol-HL 60A, 15×3 cm, 5μm; Mobile phase A: CO2; Mobile phase B: 0.1% ammonium hydroxide in 40% ethanol and 60% acetonitrile (v/v); Isocratic 73% B over 5 minutes; Flow rate: 110 mL/minute; Temperature: 45 C; Back Pressure: 100 bar; Detection: UV 254,; Injection volume: 1,400 μL; Instrument: SFC-PICLAB PREP 200. Resulting in Peak 1 (2603 - 144 mg, 97.5% purity), Peak 2 (2604 -518 mg, 97.35 purity) both were light yellow free flowing powder post lyophilization.

Compound 2603 LCMS: MS (ESI) calculated for $C_{42}H_{56}N_{12}O_6$ $[M+H]^+$, 825.3; found, 825.3

Compound 2604 LCMS: MS (ESI) calculated for $C_{42}H_{56}N_{12}O_6$ $[M+H]^+$, 825.3; found, 825.3

*<u>$^1$H NMR and LCMS for Assay Compounds</u>*

Example 2 LCMS

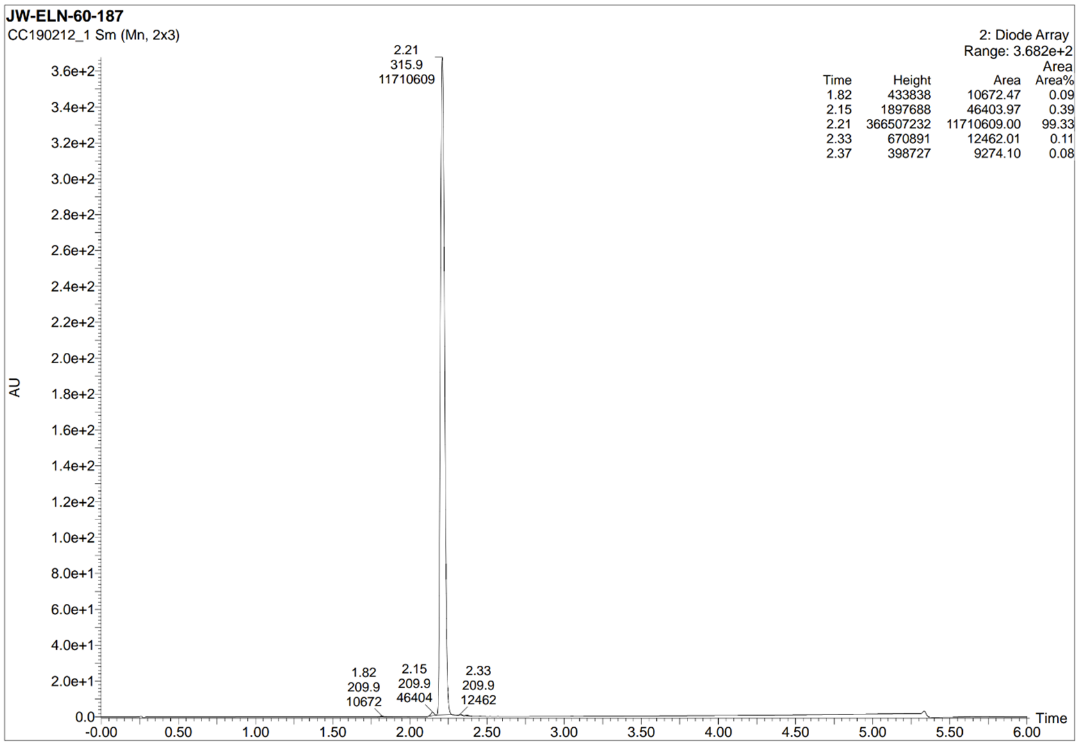


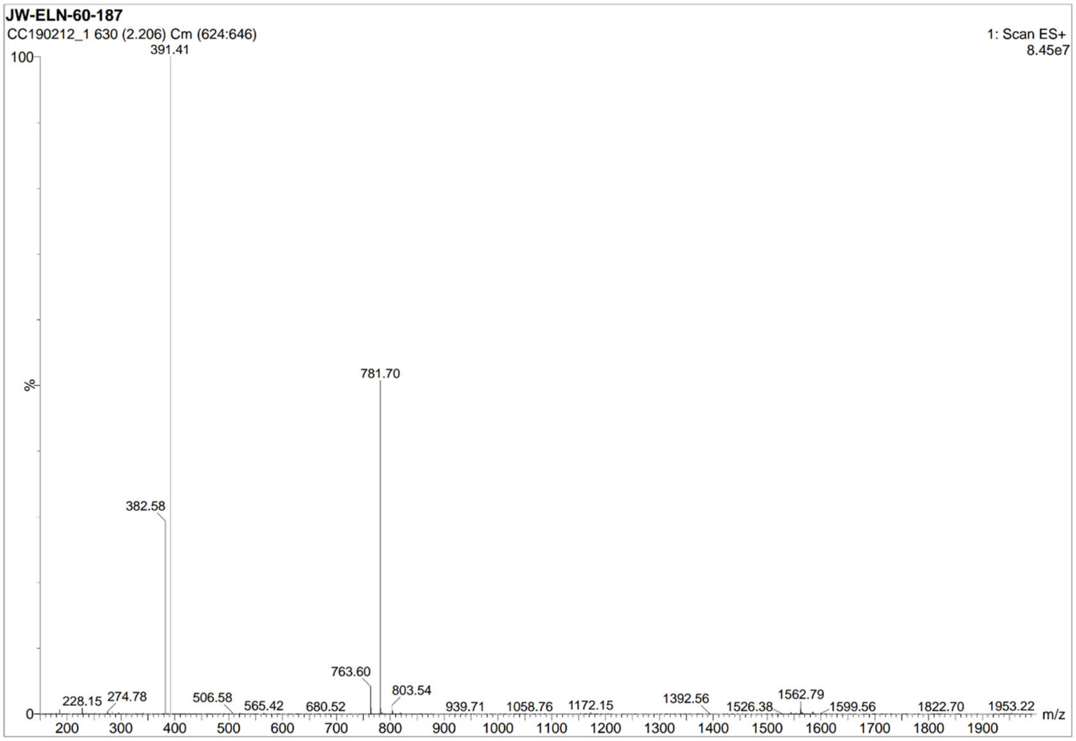


Example 2 $^{1}H$ NMR

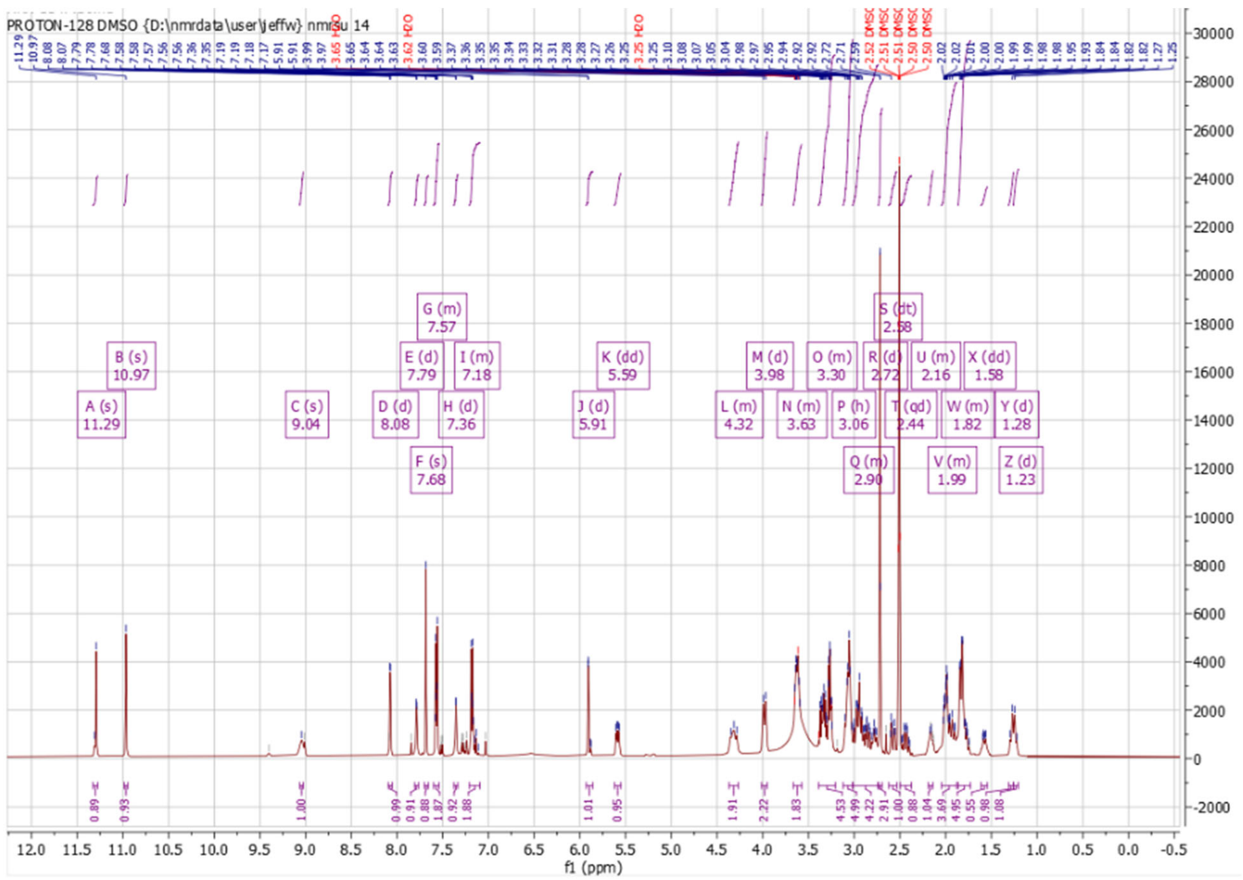
PROTON-128 DMSO {D:\nmrdata\user\jeffw} nmrsu 14
A (s) 11.29
B (s) 10.97
C (s) 9.04
D (d) 8.08
E (d) 7.79
F (s) 7.68
G (m) 7.57
H (d) 7.36
I (m) 7.18
J (d) 5.91
K (dd) 5.59
L (m) 4.32
M (d) 3.98
N (m) 3.63
O (m) 3.30
P (h) 3.06
Q (m) 2.90
R (d) 2.72
S (dt) 2.58
T (qd) 2.44
U (m) 2.16
V (m) 1.99
W (m) 1.82
X (dd) 1.58
Y (d) 1.28
Z (d) 1.23
f1 (ppm)

Example 3 LCMS

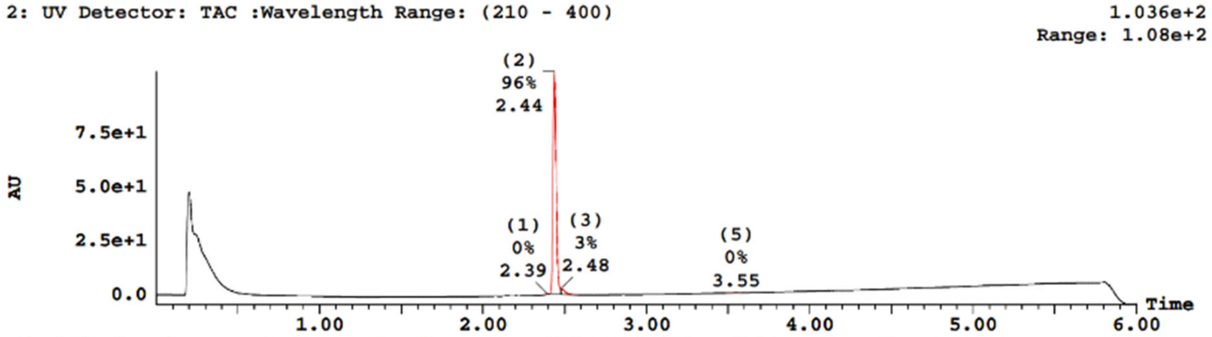


| Peak Number | Compound | Time | AreaAbs | Area %Total | Width | Height | Mass Found |
|---|---|---|---|---|---|---|---|
| 1 | | 2.39 | 6e+003 | 0.25 | 0 | 3e+005 | Not Found |
| 2 | | 2.44 | 2e+006 | 96.08 | 0 | 1e+008 | Not Found |
| 3 | | 2.48 | 7e+004 | 2.97 | 0 | 3e+006 | Not Found |
| 4 | | 2.53 | 6e+003 | 0.25 | 0 | 3e+005 | Not Found |
| 5 | | 3.55 | 1e+004 | 0.45 | 0 | 3e+005 | Not Found |

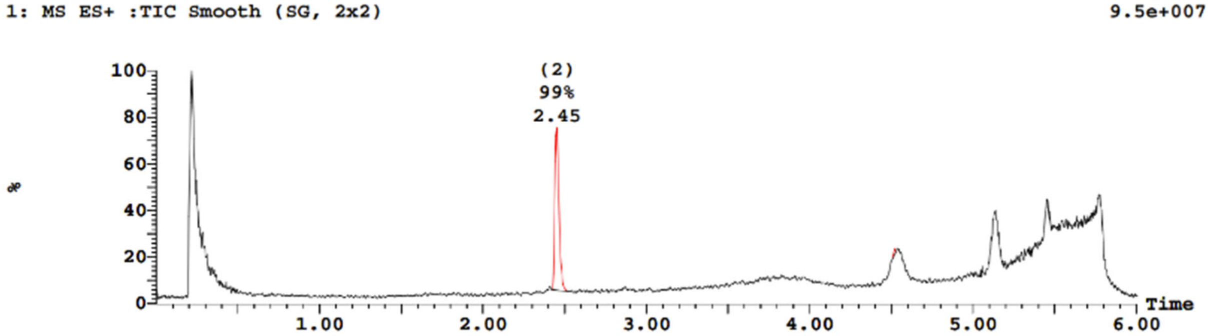


| Peak Number | Time | AreaAbs | Area %Total | Mass Found |
|---|---|---|---|---|
| 2 | 2.45 | 2e+006 | 98.52 | Not Found |
| 6 | 4.52 | 3e+004 | 1.48 | Not Found |

| Peak ID | Time | Mass Found |
|---|---|---|
| 1 | 2.39 | Not Found |

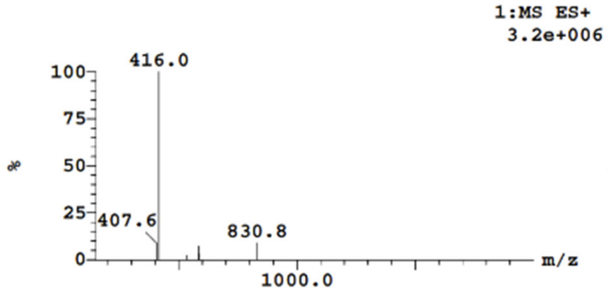


| Peak ID | Time | Mass Found |
|---|---|---|
| 2 | 2.45 | Not Found |

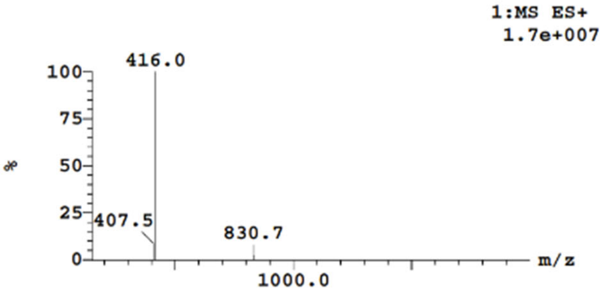


Example 3 $^{1}H$ NMR

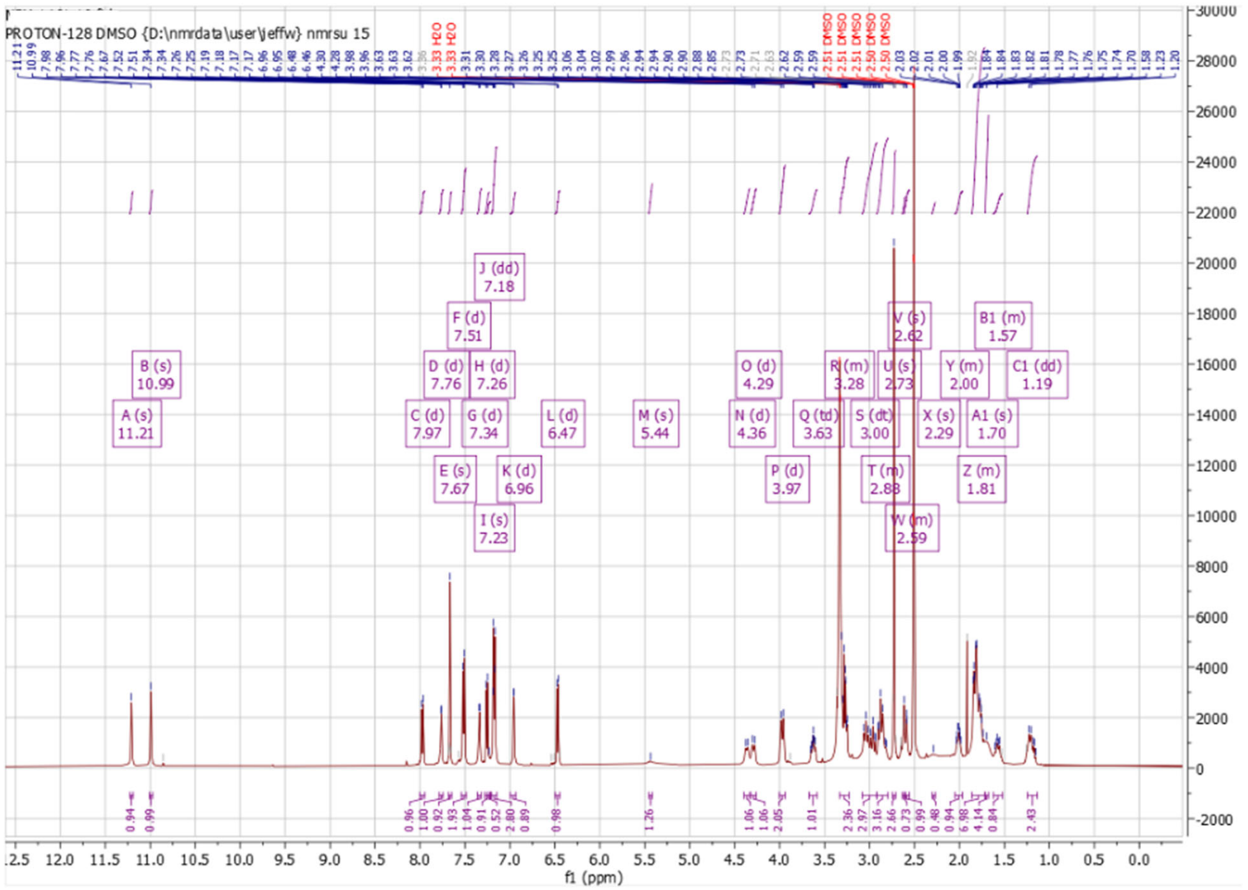


Example 4 LCMS

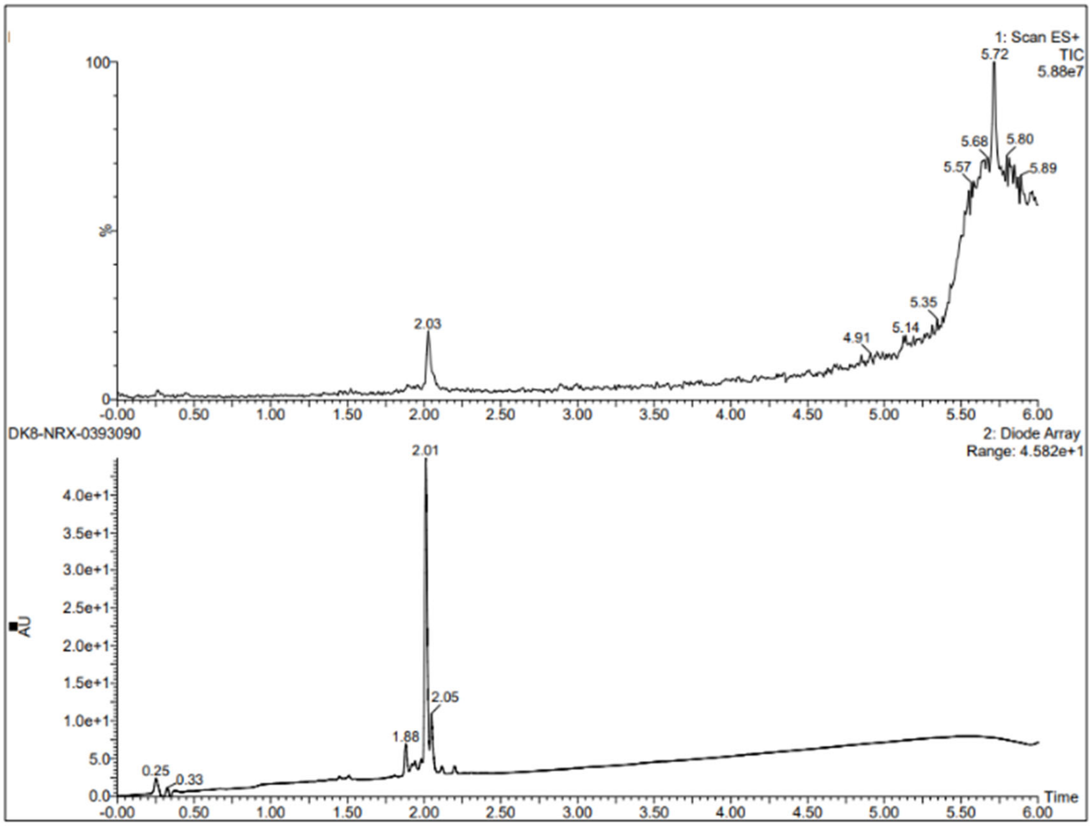
1: Scan ES+
TIC
5.88e7
5.72
5.68
5.80
5.57
5.89
5.35
5.14
4.91
2.03
DK8-NRX-0393090
2: Diode Array
Range: 4.582e+1
2.01
2.05
1.88
0.25
0.33
AU
Time

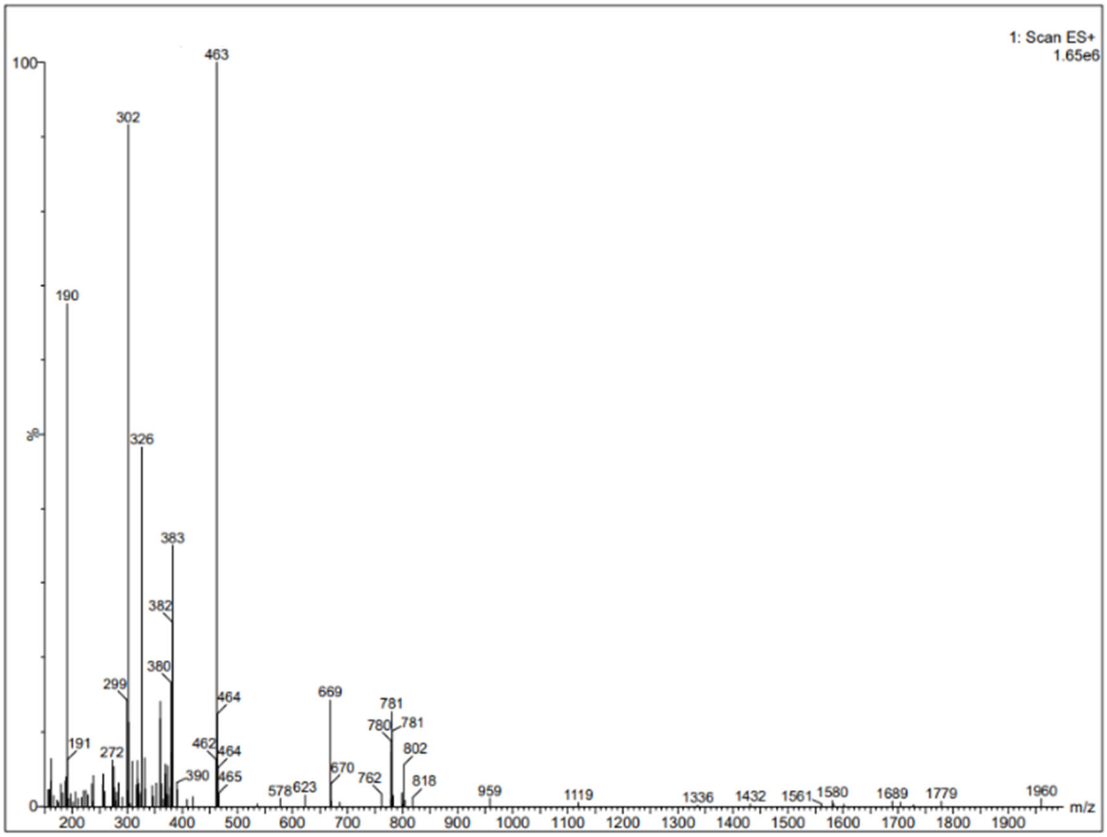
1: Scan ES+
1.65e6
463
302
190
326
383
382
380
299
464
669
781
780
802
191
272
462
390
465
578
623
670
762
818
959
1119
1336
1432
1561
1580
1689
1779
1960
m/z

Example 4 $^{1}$H NMR

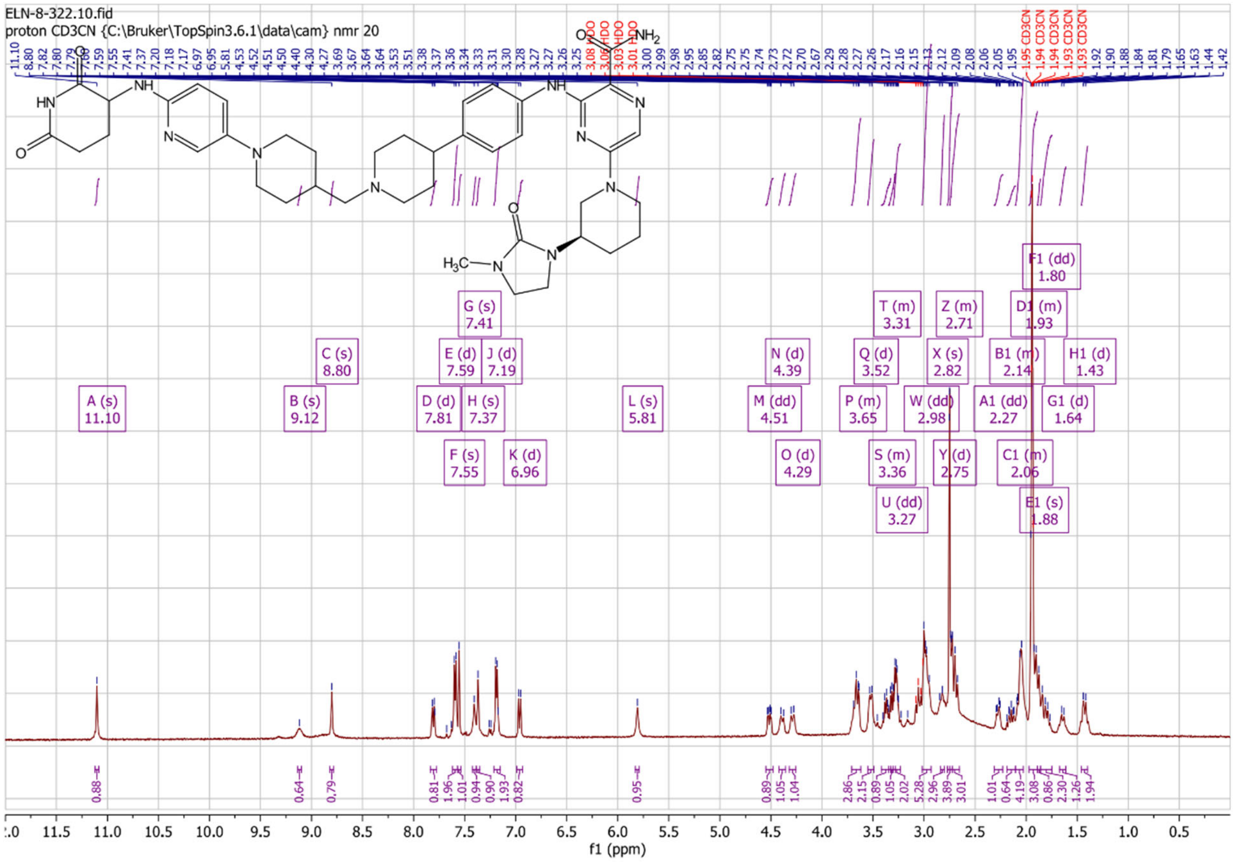

Example 5 LCMS

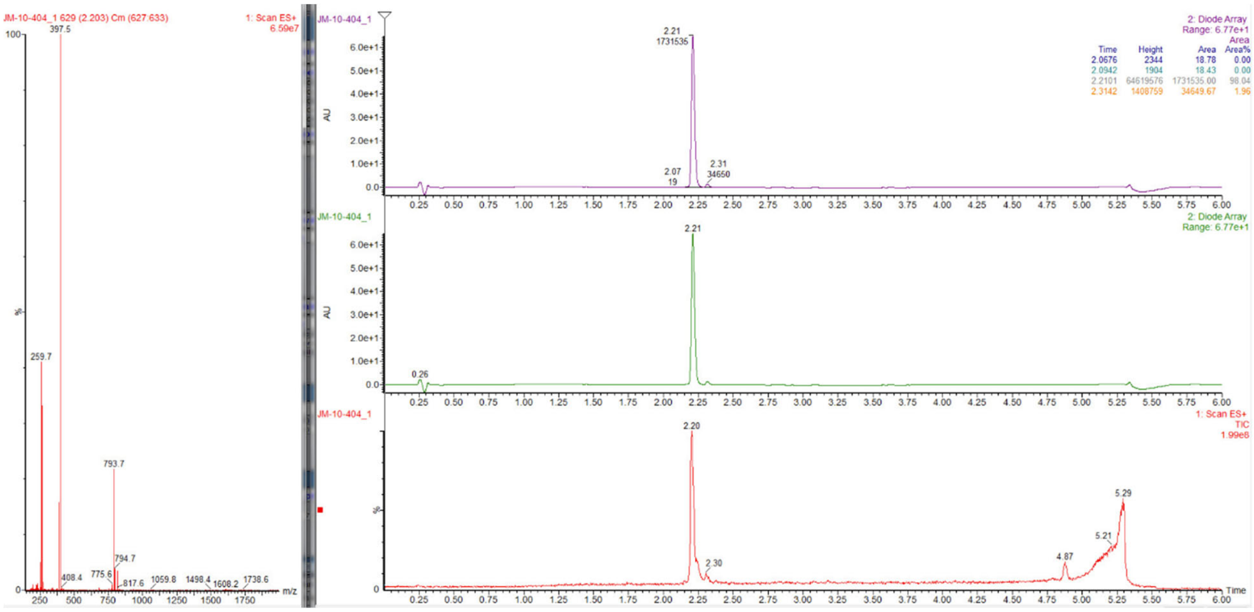


Example 5 ¹H NMR

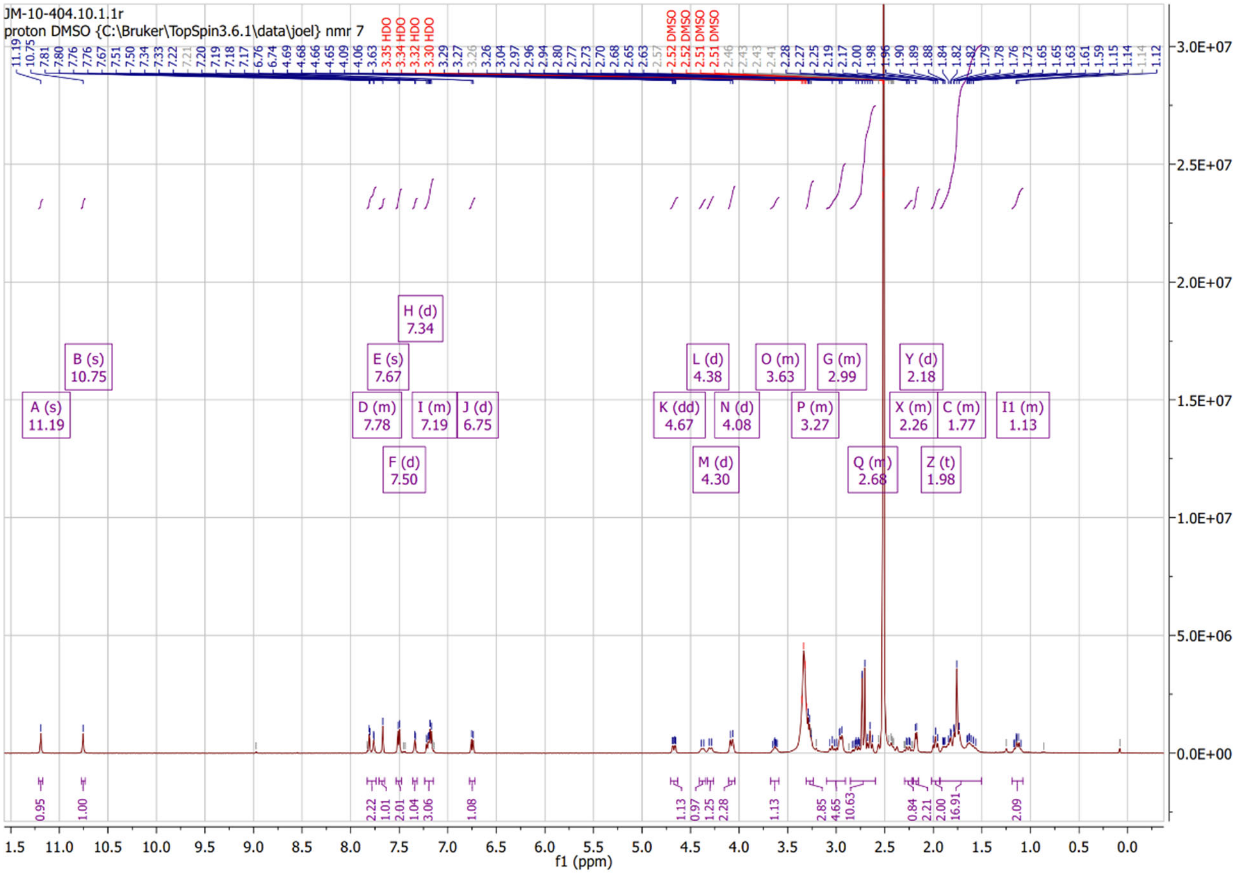


Example 6 LCMS

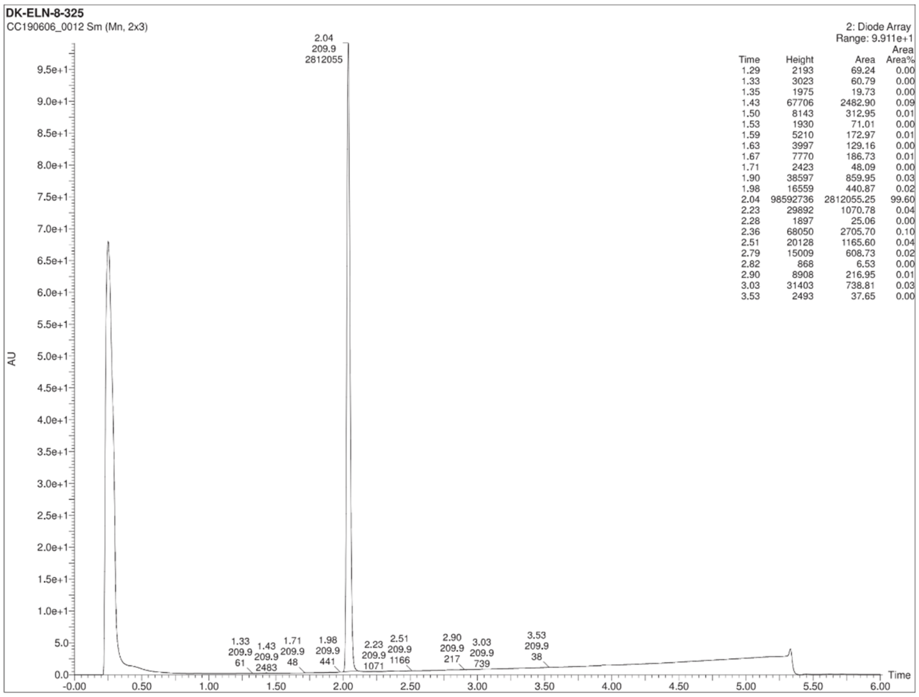


| Time | Height | Area | Area Area% |
|---|---|---|---|
| 1.29 | 2193 | 69.24 | 0.00 |
| 1.33 | 3023 | 60.79 | 0.00 |
| 1.35 | 1975 | 19.73 | 0.00 |
| 1.43 | 67706 | 2482.90 | 0.09 |
| 1.50 | 8143 | 312.95 | 0.01 |
| 1.53 | 1930 | 71.01 | 0.00 |
| 1.59 | 5210 | 172.97 | 0.01 |
| 1.63 | 3997 | 129.16 | 0.00 |
| 1.67 | 7770 | 186.73 | 0.01 |
| 1.71 | 2423 | 48.09 | 0.00 |
| 1.90 | 38597 | 859.95 | 0.03 |
| 1.98 | 16559 | 440.87 | 0.02 |
| 2.04 | 98592736 | 2812055.25 | 99.60 |
| 2.23 | 29892 | 1070.78 | 0.04 |
| 2.28 | 1897 | 25.06 | 0.00 |
| 2.36 | 68050 | 2705.70 | 0.10 |
| 2.51 | 20128 | 1165.60 | 0.04 |
| 2.79 | 15009 | 608.73 | 0.02 |
| 2.82 | 868 | 6.53 | 0.00 |
| 2.90 | 8908 | 216.95 | 0.01 |
| 3.03 | 31403 | 738.81 | 0.03 |
| 3.53 | 2493 | 37.65 | 0.00 |

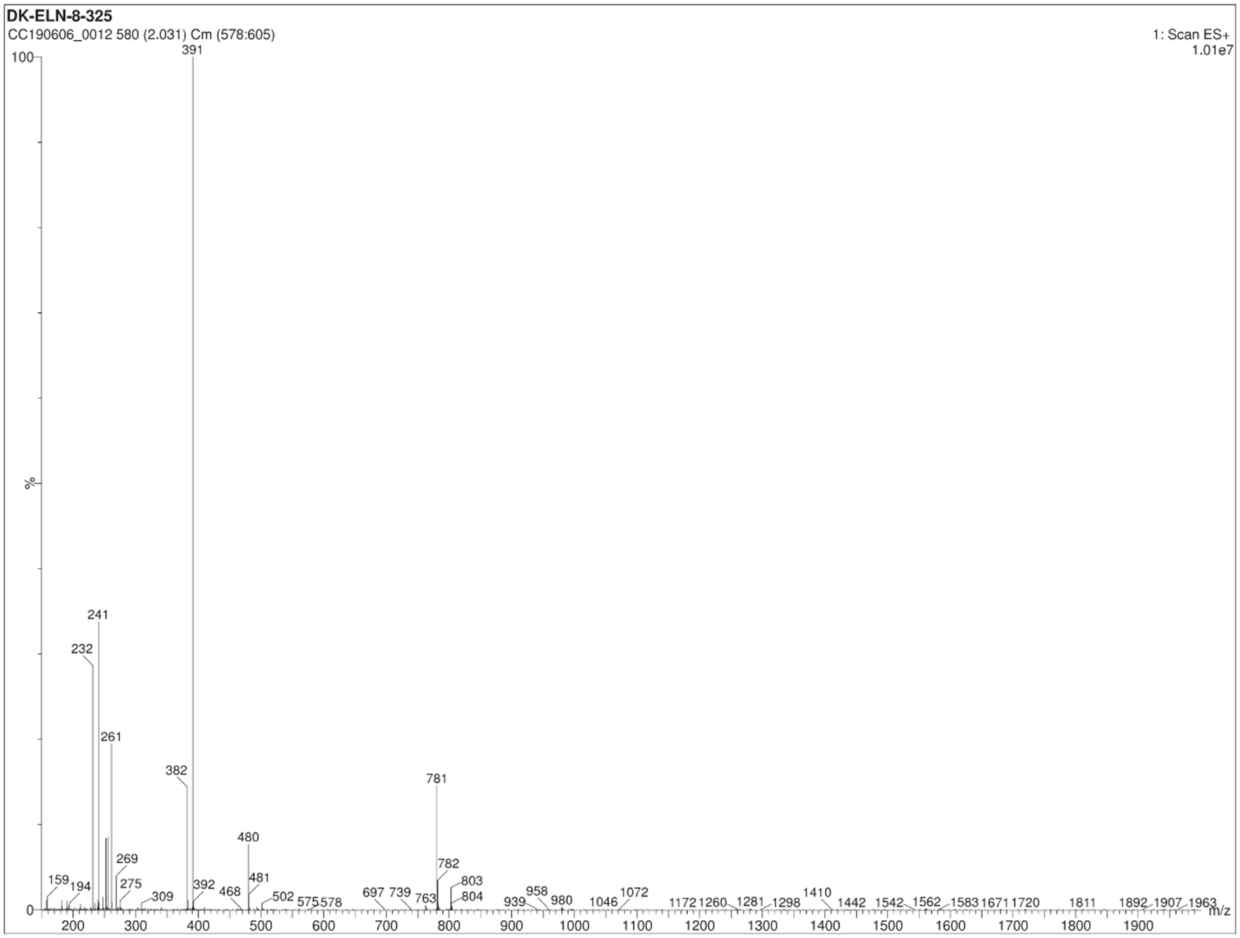

Example 6 $^{1}$H NMR

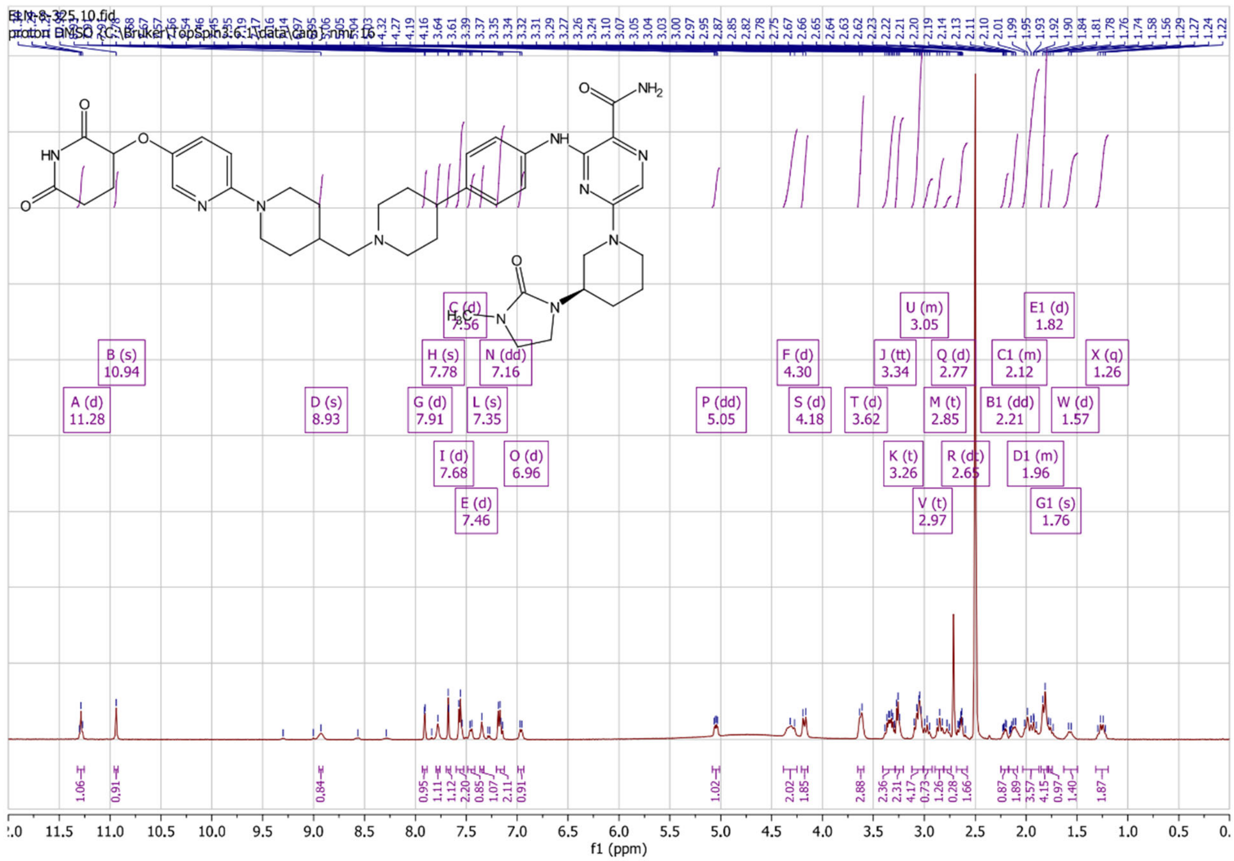

Example 7 LCMS

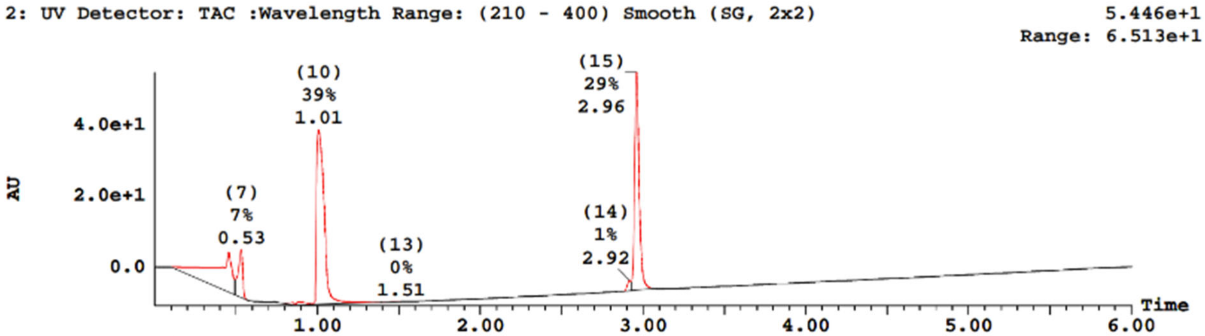


| Peak Number | Compound | Time | AreaAbs | Area %Total | Width | Height | Mass Found |
|---|---|---|---|---|---|---|---|
| 4 | | 0.45 | 2e+006 | 22.54 | 0 | 1e+007 | Not Found |
| 7 | | 0.53 | 4e+005 | 6.55 | 0 | 1e+007 | Not Found |
| 8 | | 0.85 | 3e+004 | 0.40 | 0 | 6e+005 | Not Found |
| 9 | | 0.88 | 5e+004 | 0.78 | 0 | 8e+005 | Not Found |
| 10 | | 1.01 | 3e+006 | 39.43 | 0 | 5e+007 | Not Found |
| 13 | | 1.51 | 3e+003 | 0.04 | 0 | 1e+004 | Not Found |
| 14 | | 2.92 | 7e+004 | 1.06 | 0 | 3e+006 | Not Found |
| 15 | | 2.96 | 2e+006 | 29.19 | 0 | 6e+007 | Not Found |

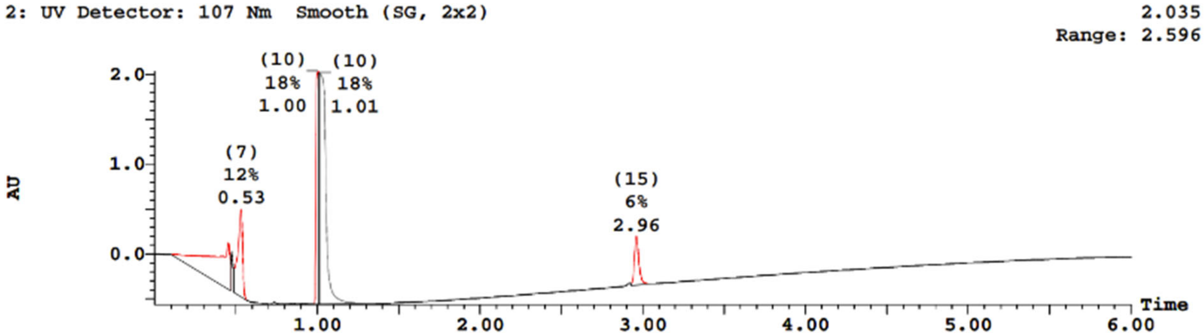


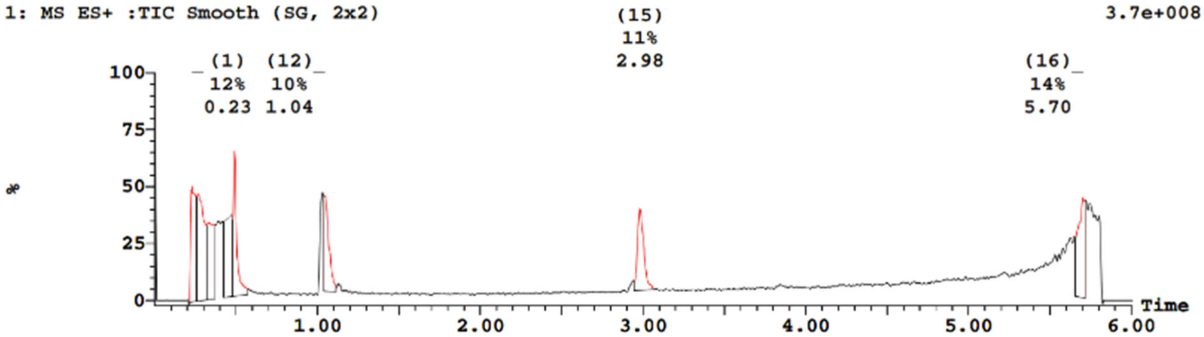

| Peak ID | Time | Mass Found |
| --- | --- | --- |
| 10 | 1.01 | Not Found |

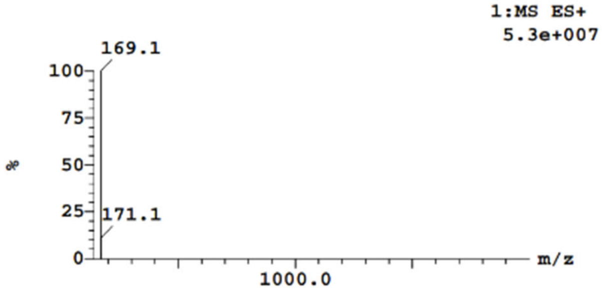


| Peak ID | Time | Mass Found |
| --- | --- | --- |
| 12 | 1.04 | Not Found |

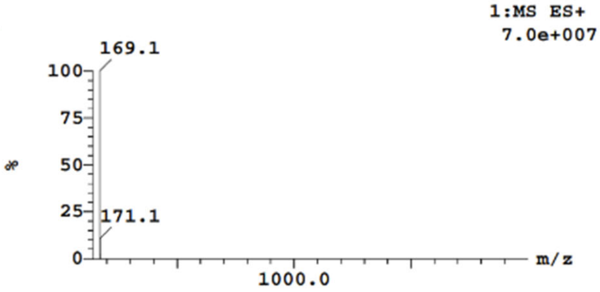


| Peak ID | Time | Mass Found |
| --- | --- | --- |
| 13 | 1.51 | Not Found |

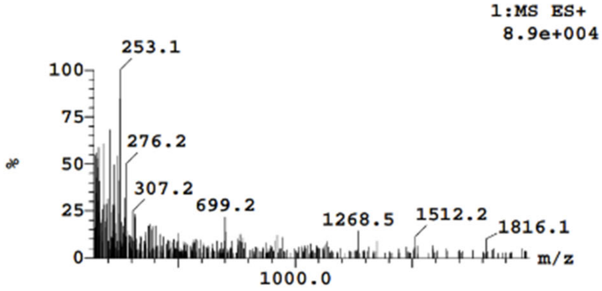


| Peak ID | Time | Mass Found |
| --- | --- | --- |
| 14 | 2.92 | Not Found |

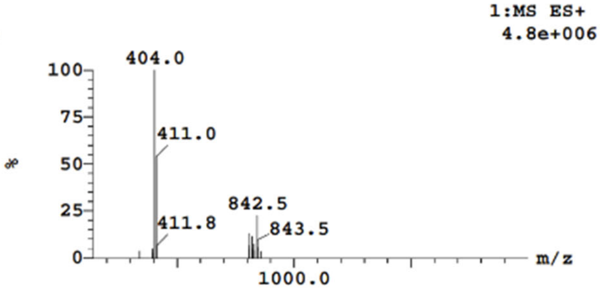


| Peak ID | Time | Mass Found |
| --- | --- | --- |
| 15 | 2.98 | Not Found |

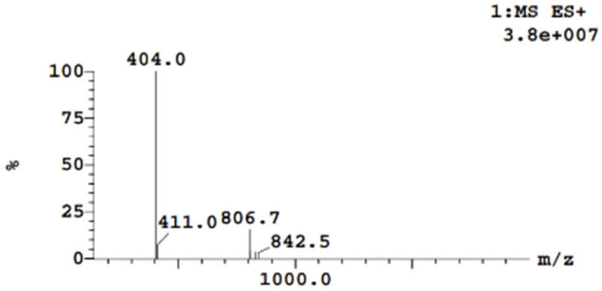


| Peak ID | Time | Mass Found |
| --- | --- | --- |
| 16 | 5.70 | Not Found |

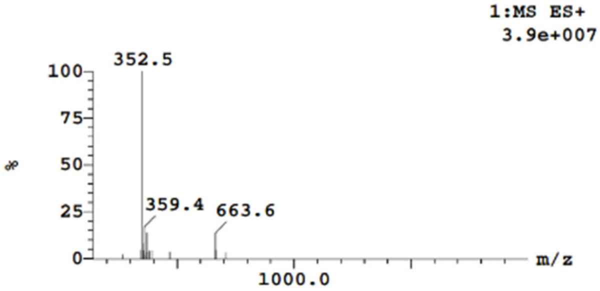

Example 7 $^{1}$H NMR

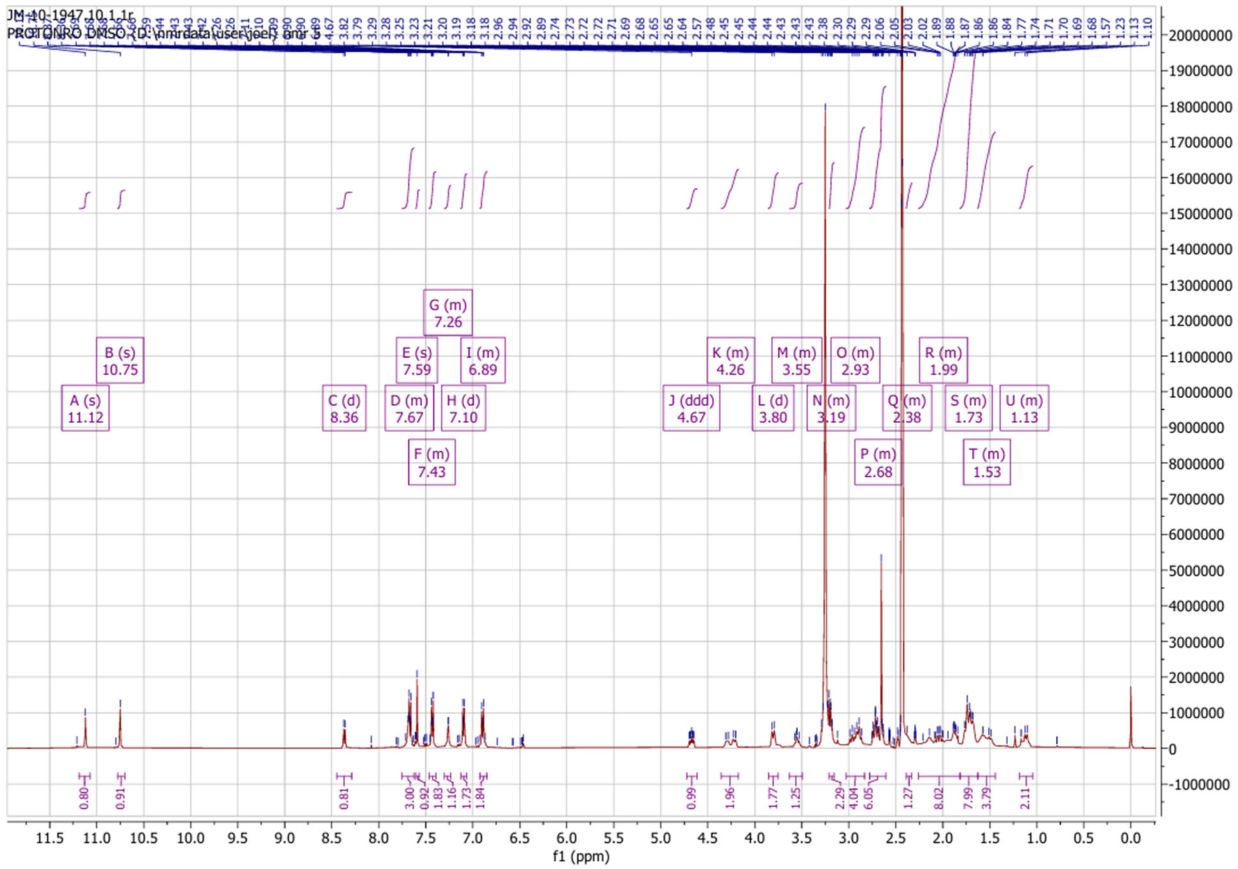

Example 8 LCMS

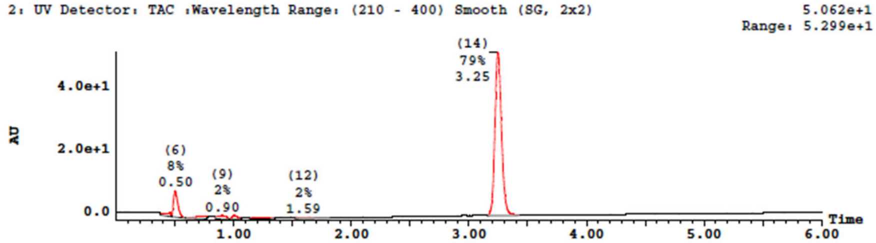


| Peak Number | Compound | Time | AreaAbs | Area %Total | Width | Height | Mass Found |
|---|---|---|---|---|---|---|---|
| 5 | | 0.46 | 8e+004 | 1.83 | 0 | 1e+006 | Not Found |
| 6 | | 0.50 | 4e+005 | 8.28 | 0 | 8e+006 | Not Found |
| 8 | | 0.73 | 8e+004 | 1.92 | 0 | 7e+005 | Not Found |
| 9 | | 0.90 | 1e+005 | 2.41 | 0 | 1e+006 | Not Found |
| 10 | | 1.01 | 8e+004 | 1.80 | 0 | 1e+006 | Not Found |
| 11 | | 1.16 | 1e+005 | 2.47 | 0 | 6e+005 | Not Found |
| 12 | | 1.59 | 1e+005 | 2.23 | 0 | 4e+005 | Not Found |
| 14 | | 3.25 | 3e+006 | 79.06 | 0 | 5e+007 | Not Found |

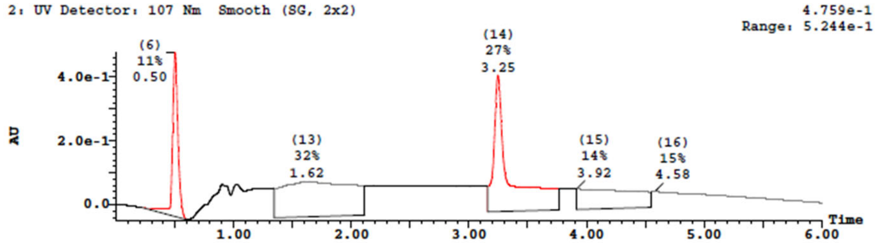


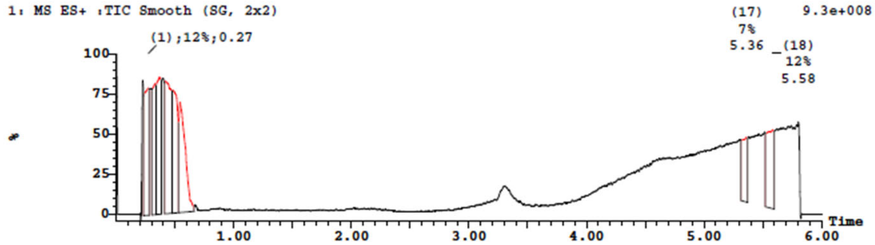


| Peak ID | Time | Mass Found |
|---|---|---|
| 14 | 3.25 | Not Found |

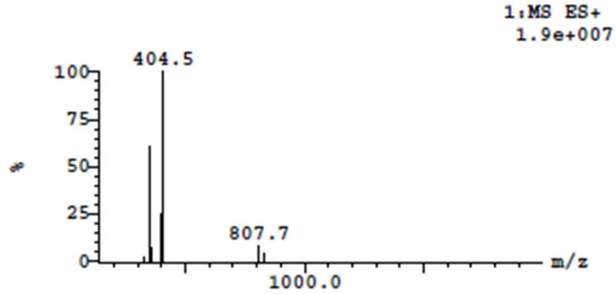


| Peak ID | Time | Mass Found |
|---|---|---|
| 17 | 5.36 | Not Found |

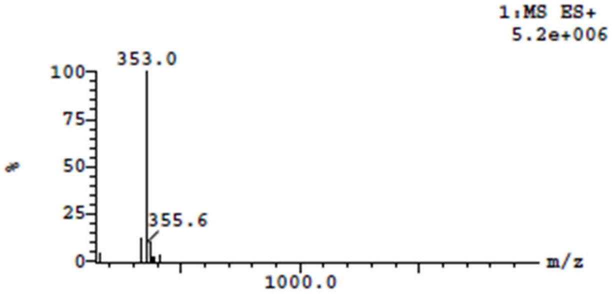

Example 8 $^{1}$H NMR

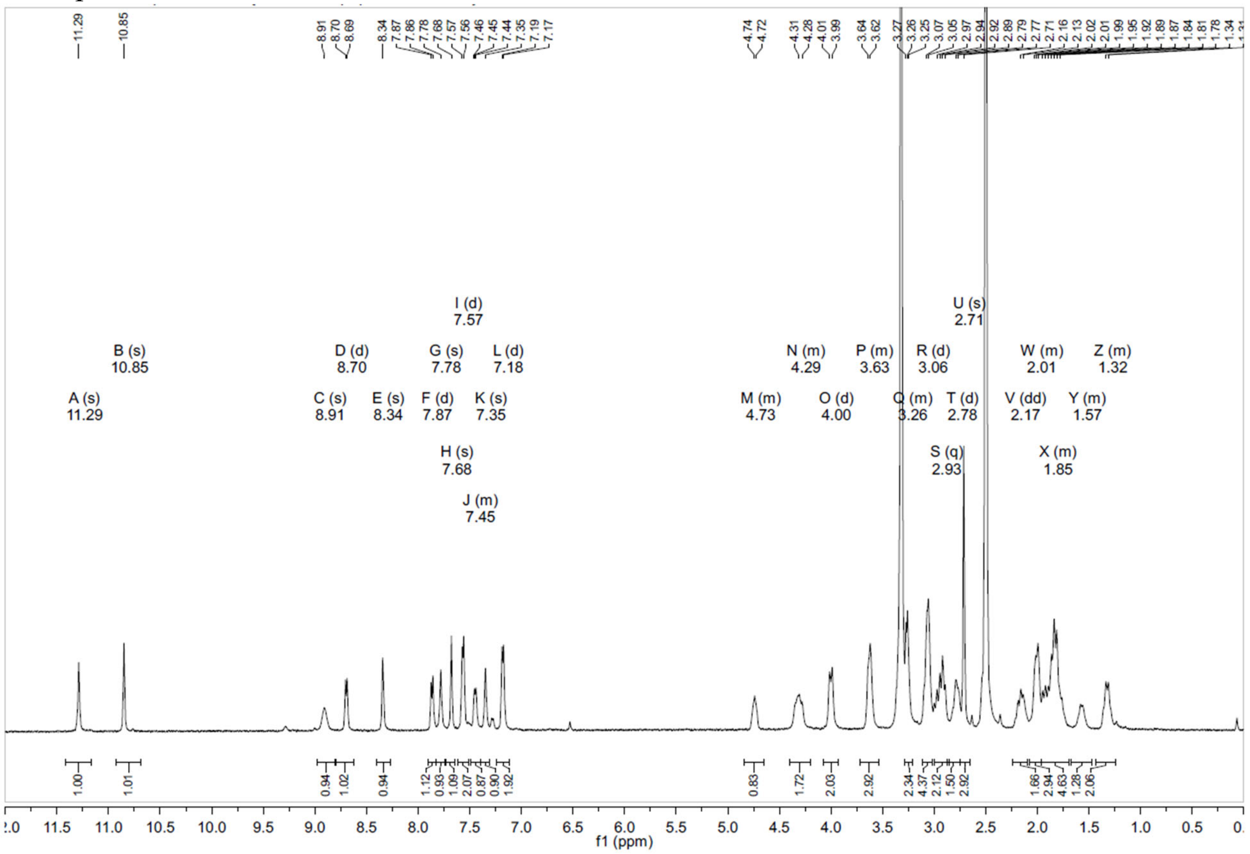

Example 9 LCMS

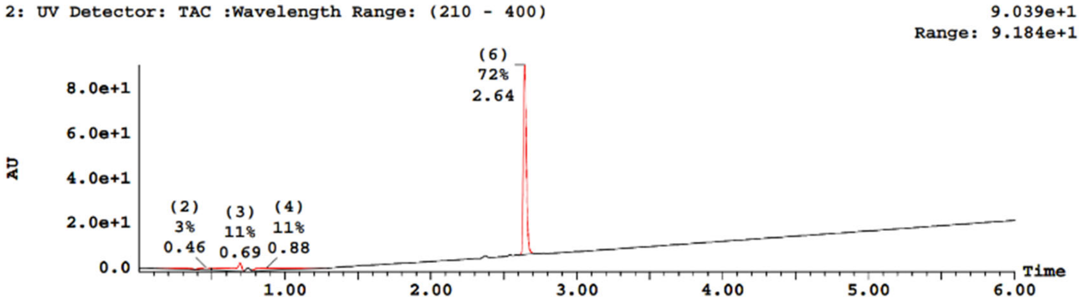


| Peak Number | Compound | Time | AreaAbs | Area %Total | Width | Height | Mass Found |
|---|---|---|---|---|---|---|---|
| 1 | | 0.17 | 9e+004 | 3.37 | 0 | 3e+005 | Not Found |
| 2 | | 0.46 | 9e+004 | 3.16 | 0 | 1e+006 | Not Found |
| 3 | | 0.69 | 3e+005 | 10.63 | 0 | 4e+006 | Not Found |
| 4 | | 0.88 | 3e+005 | 10.64 | 1 | 1e+006 | Not Found |
| 6 | | 2.64 | 2e+006 | 72.20 | 0 | 8e+007 | Not Found |

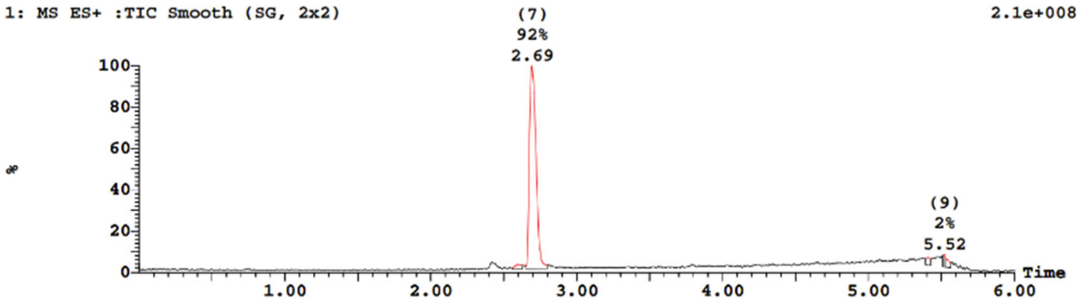


| Peak Number | Time | AreaAbs | Area %Total | Mass Found |
|---|---|---|---|---|
| 5 | 2.59 | 2e+005 | 2.01 | Not Found |
| 7 | 2.69 | 1e+007 | 92.15 | Not Found |
| 8 | 5.41 | 3e+005 | 2.14 | Not Found |
| 9 | 5.52 | 2e+005 | 1.64 | Not Found |
| 10 | 5.53 | 2e+005 | 2.06 | Not Found |

| Peak ID | Time | Mass Found |
|---|---|---|
| 3 | 0.69 | Not Found |

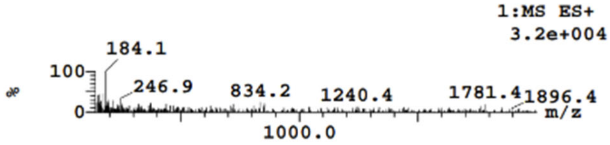


| Peak ID | Time | Mass Found |
|---|---|---|
| 4 | 0.88 | Not Found |

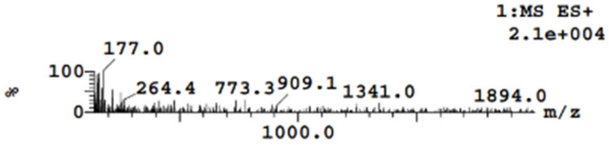


| Peak ID | Time | Mass Found |
|---|---|---|
| 5 | 2.59 | Not Found |

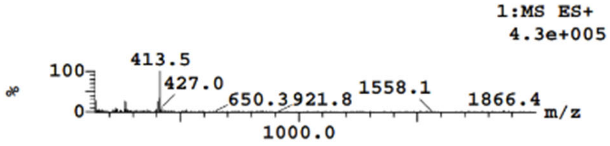


| Peak ID | Time | Mass Found |
|---|---|---|
| 6 | 2.64 | Not Found |

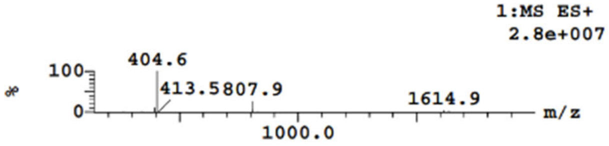


| Peak ID | Time | Mass Found |
|---|---|---|
| 7 | 2.69 | Not Found |

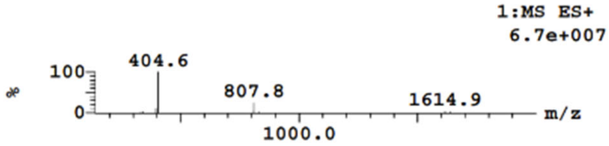


| Peak ID | Time | Mass Found |
|---|---|---|
| 8 | 5.41 | Not Found |

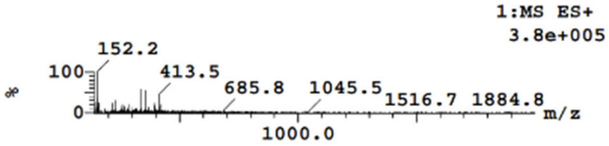

Example 9 $^{1}$H NMR

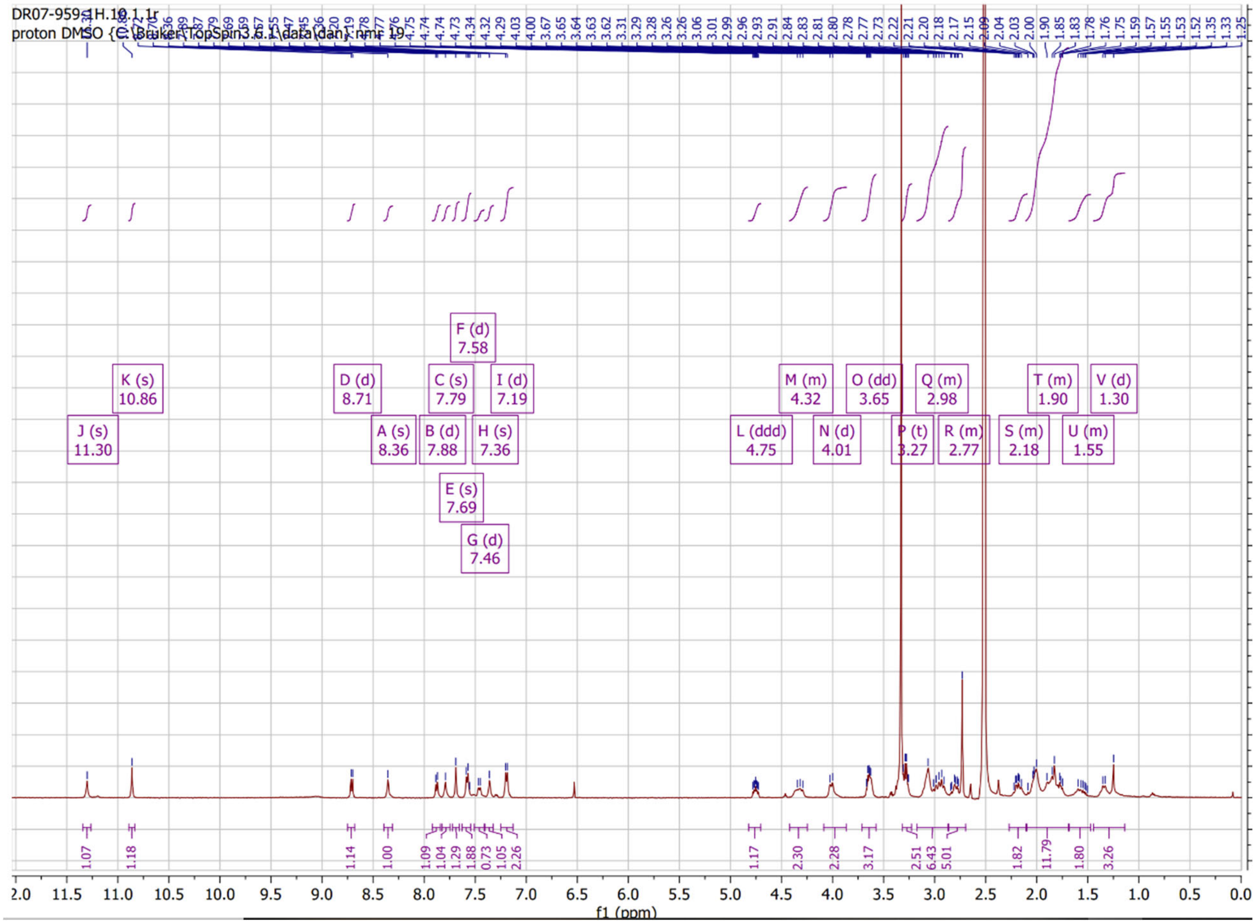

Example 10 LCMS

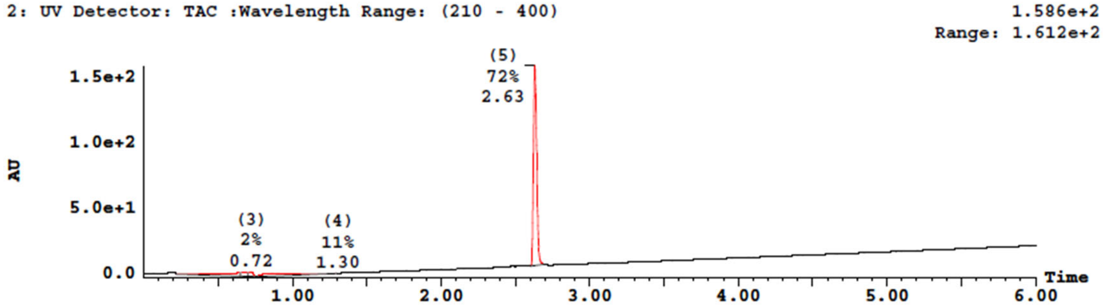


| Peak Number | Compound | Time | AreaAbs | Area %Total | Width | Height | Mass Found |
|---|---|---|---|---|---|---|---|
| 1 | | 0.64 | 6e+005 | 11.95 | 0 | 3e+006 | Not Found |
| 2 | | 0.67 | 1e+005 | 2.65 | 0 | 3e+006 | Not Found |
| 3 | | 0.72 | 1e+005 | 2.33 | 0 | 4e+006 | Not Found |
| 4 | | 1.30 | 6e+005 | 11.34 | 1 | 2e+005 | Not Found |
| 5 | | 2.63 | 4e+006 | 71.74 | 0 | 2e+008 | Not Found |

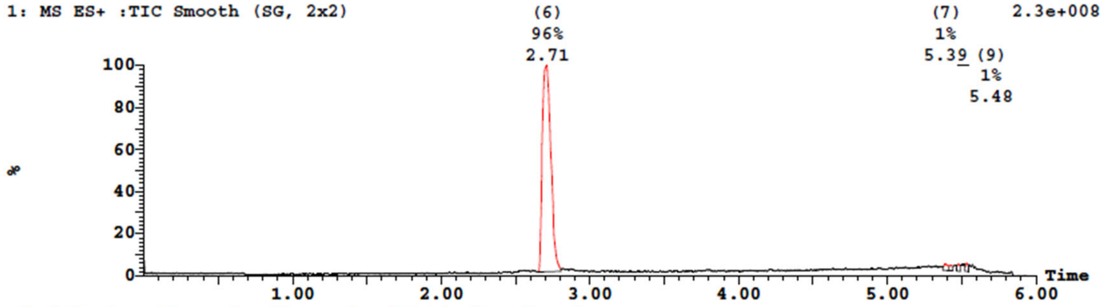


| Peak Number | Time | AreaAbs | Area %Total | Mass Found |
|---|---|---|---|---|
| 6 | 2.71 | 2e+007 | 95.60 | Not Found |
| 7 | 5.39 | 2e+005 | 1.38 | Not Found |
| 8 | 5.42 | 2e+005 | 0.96 | Not Found |
| 9 | 5.48 | 2e+005 | 1.05 | Not Found |
| 10 | 5.53 | 2e+005 | 1.00 | Not Found |

| Peak ID | Time | Mass Found |
|---|---|---|
| 3 | 0.72 | Not Found |

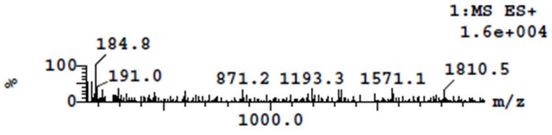


| Peak ID | Time | Mass Found |
|---|---|---|
| 4 | 1.30 | Not Found |

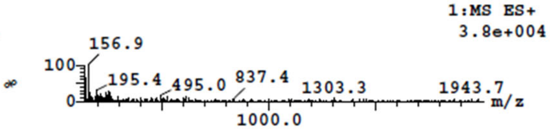


| Peak ID | Time | Mass Found |
|---|---|---|
| 5 | 2.63 | Not Found |

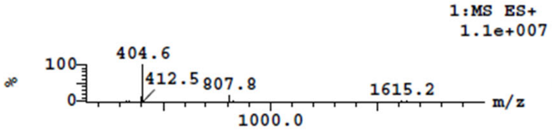


| Peak ID | Time | Mass Found |
|---|---|---|
| 6 | 2.71 | Not Found |

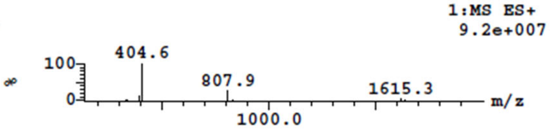

Example 10 $^{1}$H NMR

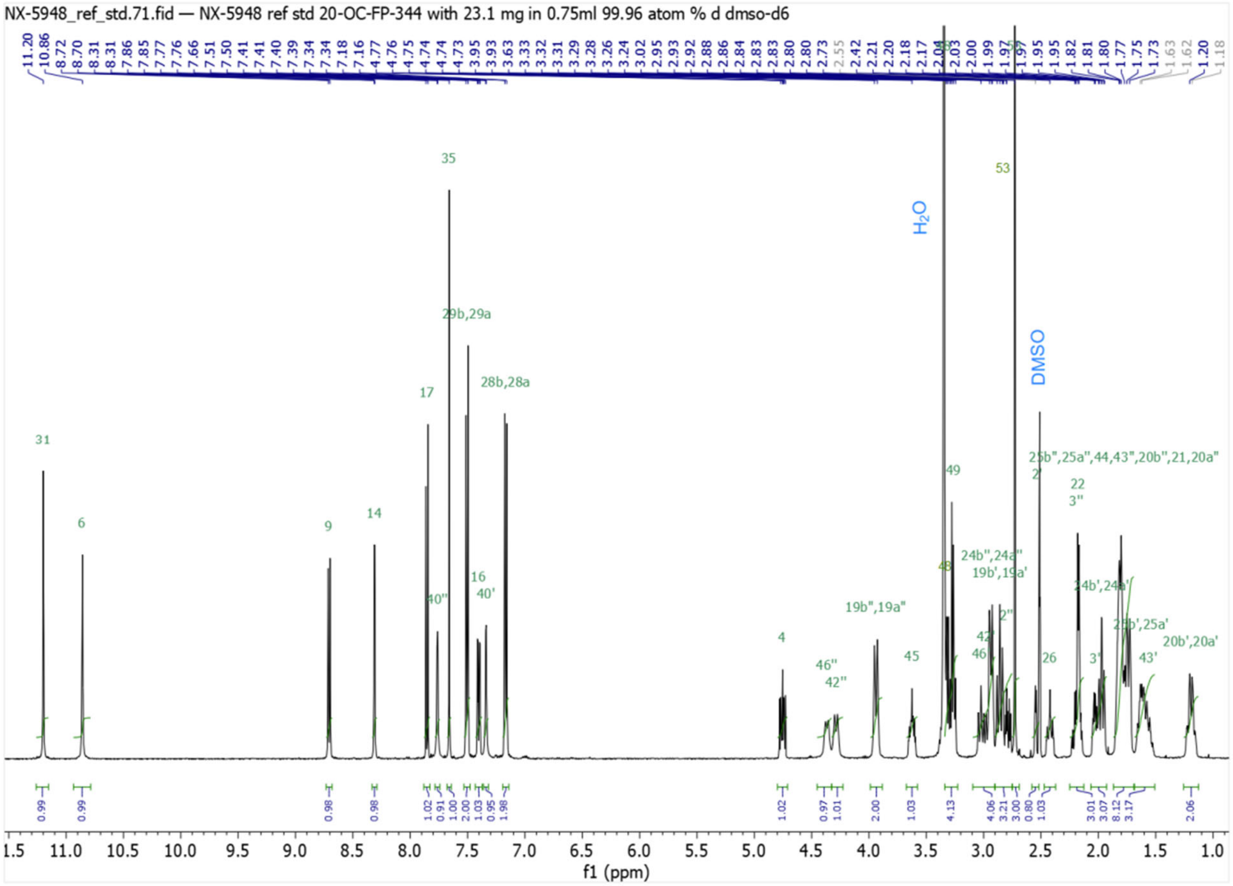

Example 10 $^{13}C$ NMR

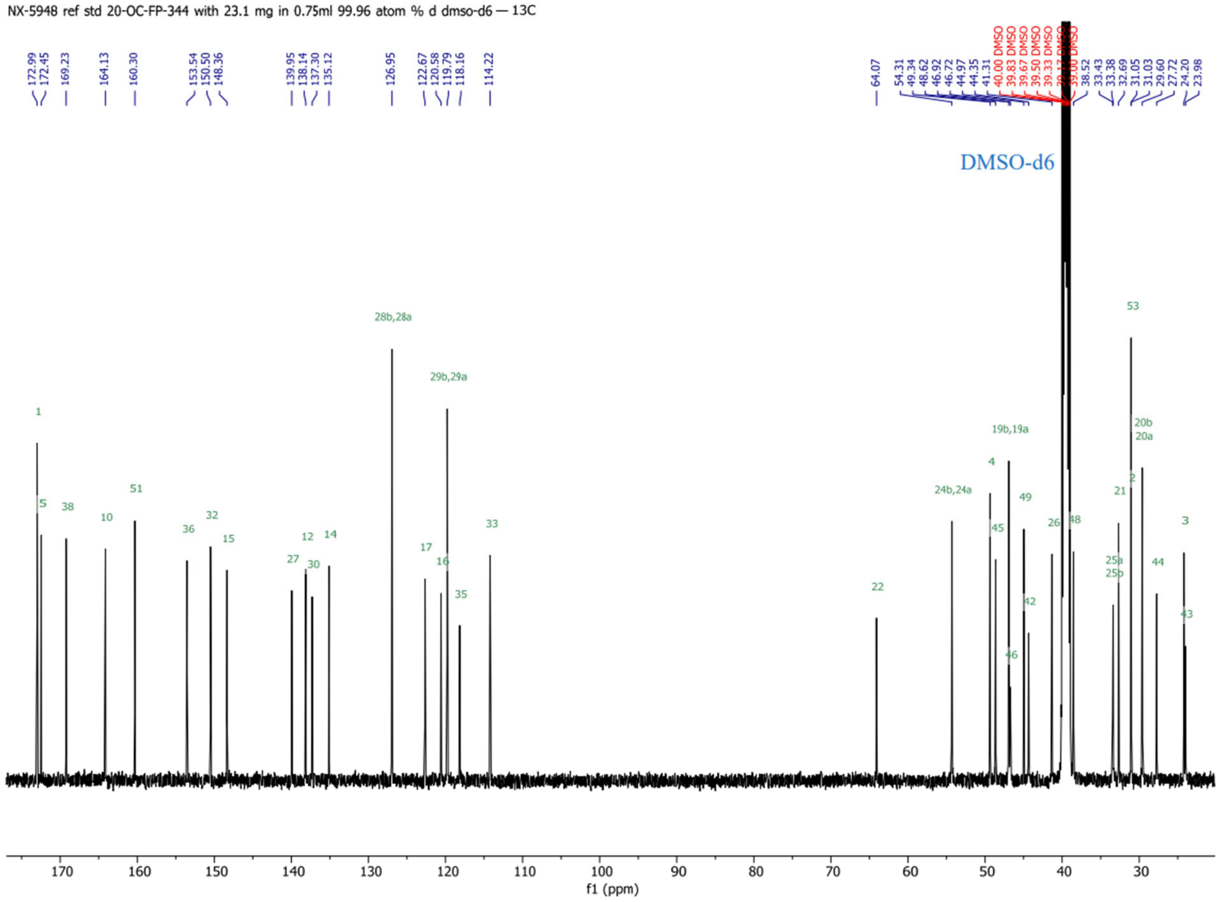


Compound **2603**

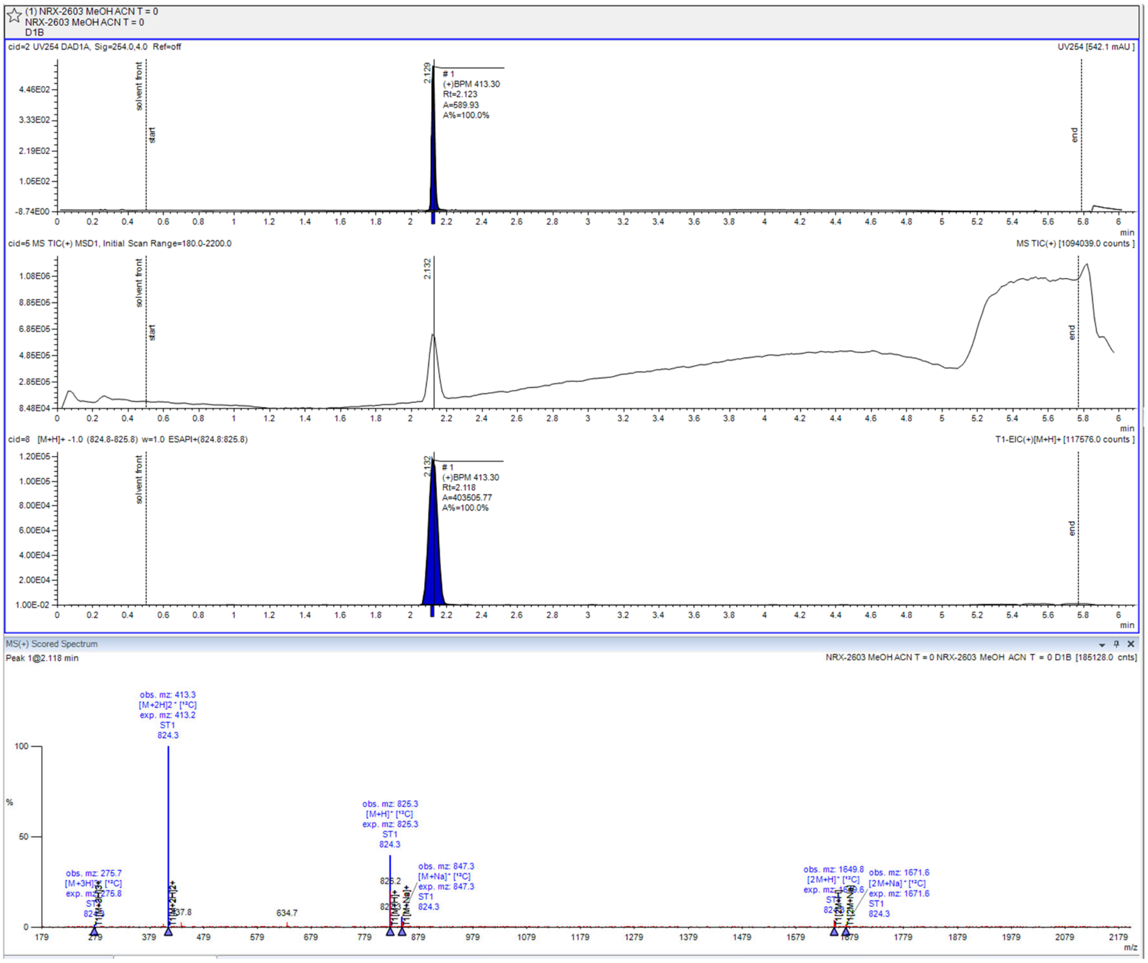
(1) NRX-2603 MeOH ACN T = 0
NRX-2603 MeOH ACN T = 0
D1B
cid=2 UV254 DAD1A, Sig=254.0,4.0 Ref=off
UV254 [542.1 mAU ]
#1
(+)BPM 413.30
Rt=2.123
A=589.93
A%=100.0%
cid=5 MS TIC(+) MSD1, Initial Scan Range=180.0-2200.0
MS TIC(+) [1094039.0 counts ]
cid=8 [M+H]+ -1.0 (824.8-825.8) w=1.0 ESAPI+(824.8:825.8)
T1-EIC(+)[M+H]+ [117576.0 counts ]
#1
(+)BPM 413.30
Rt=2.118
A=403505.77
A%=100.0%
MS(+) Scored Spectrum
Peak 1@2.118 min
NRX-2603 MeOH ACN T = 0 NRX-2603 MeOH ACN T = 0 D1B [185128.0 cnts]
obs. mz: 413.3
[M+2H]2+ [12C]
exp. mz: 413.2
ST1
824.3
obs. mz: 825.3
[M+H]+ [12C]
exp. mz: 825.3
ST1
824.3
obs. mz: 847.3
[M+Na]+ [12C]
exp. mz: 847.3
ST1
824.3
obs. mz: 275.7
[M+3H]3+ [12C]
exp. mz: 275.8
obs. mz: 1649.8
[2M+H]+ [12C]
obs. mz: 1671.6
[2M+Na]+ [12C]
exp. mz: 1671.6
ST1
824.3
634.7
m/z
min

Compound **2604**

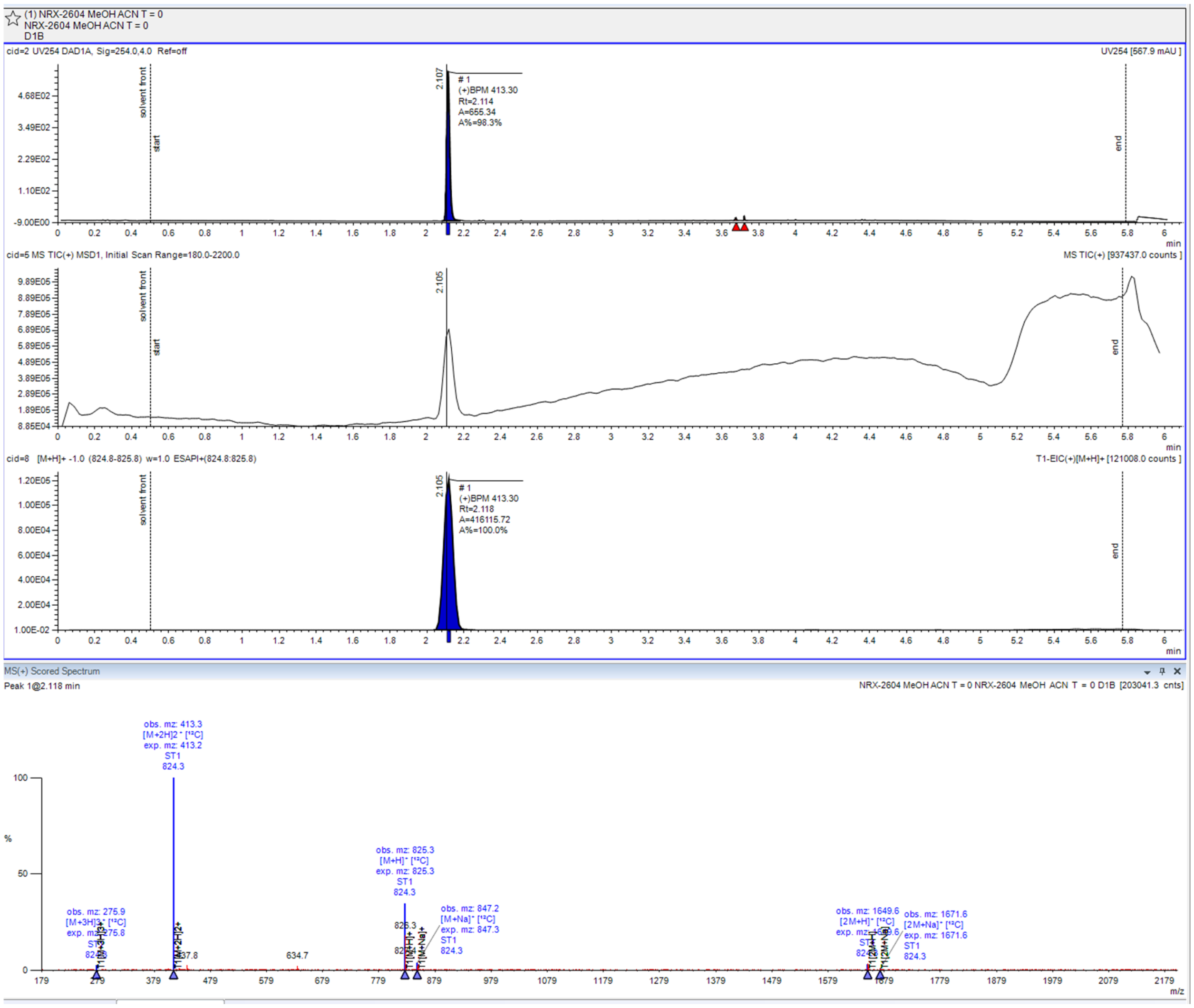
